# Nucleolar and Chromatin Remodeling during γ-Irradiation-Induced Mitotic Catastrophe: Live-Cell Imaging Correlated with UBTF and Fibrillarin 3D Redistribution

**DOI:** 10.1101/2025.08.20.671252

**Authors:** N. Makhatadze, L. Nadaraia, M. Gogebashvili, M. Nanikashvili, V. Okuneva, N. Ivanishvili, L. Rusishvili, P. Tchelidze

**Affiliations:** Carl Zeiss Education and Scientific Center, New Vision University (NVU), Tbilisi, Georgia; Republic Center of Structure Research, Georgian Technical University (GTU), Tbilisi, Georgia; I. Beritashvili Center for Experimental Biomedicine, Laboratory of Radiation Safety Problems, Tbilisi, Georgia; Georgian Scientific Industries (GSI), Tbilisi, Georgia; Iv. Javakhishvili Tbilisi State University (TSU), Tbilisi, Georgia

**Keywords:** γ-Irradiation, Mitotic Catastrophe, Nucleolar Remodeling, NAC Mobility, UBTF and Fibrillarin Redistribution, Volume LM Imaging

## Abstract

Study explores the detailed chronology of nuclear and nucleolar pre-mortal 2D dynamics and 3D structural remodeling during 30 Gy γ-irradiation-induced mitotic catastrophe and apoptosis. Using post-irradiation time-lapse imaging over 72 hours, we analyzed nuclear deformation, nucleolar components and nucleolus-associated condensed chromatin remodeling in histone H2B-GFP tagged He-La cells. To assess the 3D redistribution of major nucleolus-specific proteins and related resident sub-compartments we visualized the most stress-sensing nucleolar components using anti-UBTF and anti-fibrillarin immunolabeling. Post-irradiation time-lapse imaging revealed a three stages of multinucleation: (i) progressive nuclear invagination leading to a lobulated shape, (ii) asymmetric nuclear fragmentation into unequal-sized micronucleoli, via amitosis and (iii) endomitotic division of nuclear fragments, followed by apoptotic nuclear degradation. γ-Irradiation-induced nucleolar changes resembled nucleolus associated chromatin rearrangements observed with chemical inhibitors. Unlike classical pattern of rRNA synthesis inhibition, nucleolar segregation or capping were not observed. Even after 12 and 24 hours, nucleoli remained large, irregularly shaped, revealing decrease of UBTF-positive nucleolar components number accompanied with sufficient enlargement of their sizes. These changes suggest a pre-segregated state rather than complete nucleolar segregation. During 48-72 hours we observed emergence of giant UBTF-positive structures, however typical pattern that finalizes nucleolar segregation by formation of crescent like UBTF- and fibrillarin-positive caps, have never been registered. Our findings provide new look into the impact of γ-irradiation on nuclear and nucleolar architecture, with implications for understanding chromatin and major nucleolar components dynamics in radiation-induced damage response leading to MC and apoptosis.

**Graphic Abstract:** 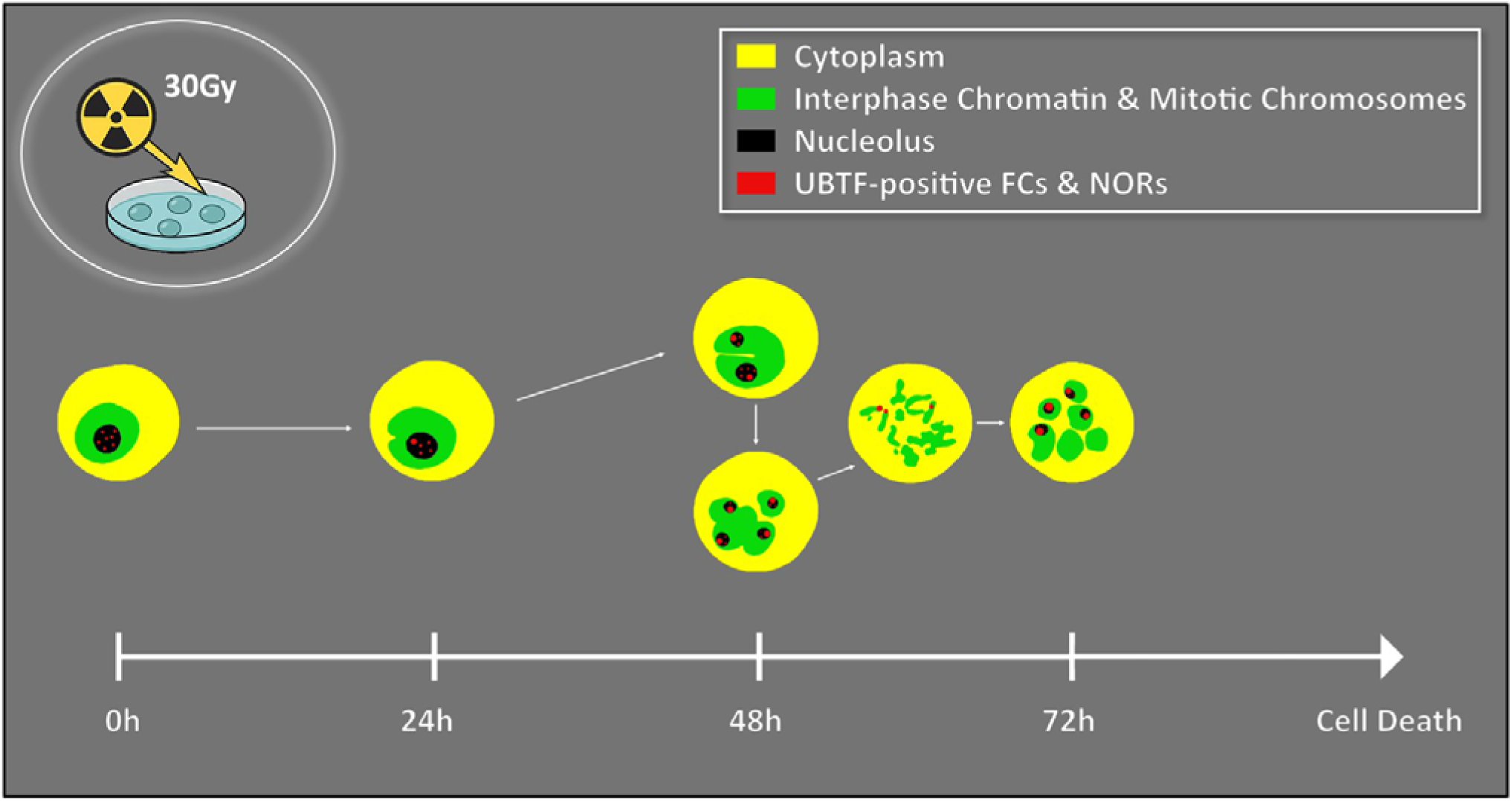

## 1. Introduction

Induced by ionizing radiation, ultraviolet (UV) light, chemotherapeutic drugs, certain viral infections, and other physical or chemical agents, DNA damage leads to a broad range of cell-death-associated pathologies, among which mitotic catastrophe (MC) is considered a major form [1–43]. Genotoxic insults such as γ-irradiation disrupt the integrity of the genome and activate diverse damage response pathways that eventually lead to cell death. Each pathway is distinguished by its specific morphological signature of pre-mortal degradation. These abnormalities often culminate in aberrant mitosis, mitotic arrest, or premature entry into mitosis, ultimately expressed through apoptosis, necrosis, autophagy, or mixed forms of post-MC death [4, 6, 8, 9, 13, 15, 18, 22, 28, 31, 37, 39, 41, 43].

Ionizing radiation inflicts DNA single- and double-strand breaks that activate checkpoint mechanisms, aiming to halt the cell cycle and facilitate repair. However, when the damage is extensive and the repair system is overwhelmed or impaired, cells may bypass checkpoint arrest and enter an aberrant mitosis. As a result, distorted chromosomal segregation leads to micronucleation, multinucleation, and the formation of deformed or fragmented nuclei [5, 6, 10, 24, 27, 37, 41, 43]. The interest in understanding the molecular and structural basis of MC has been intensified by its relevance to the clinical outcomes of radiotherapy (RT) in solid tumors. Developing therapeutic strategies that enhance MC (EMC) holds considerable promise for improving RT effectiveness through EMC, particularly in cases involving apoptosis-resistant malignancies [3, 13, 21, 25, 26, 29, 32].

The progression toward MC and its associated aberrant mitosis is tightly linked to disorders in cellular biosynthetic capacity, particularly ribosome production. Ribosome biogenesis is coordinated by molecular machineries, governing transcription of rRNA genes (r-genes), pre-rRNA processing, and assembly of pre-ribosomal particles. These machineries are structurally integrated to form the nucleolus, a central nuclear compartment essential for maintaining cellular metabolism. The nucleolus houses ribosomal DNA chromatin (r-chromatin) - large head-to-tail tandem arrays of r-genes, structurally organized by hundreds of transcriptional unit electron microscopically distinguished in form of classical “Christmas Trees” [44–67]. Therefore, structural framework of the nucleolus is anchored by extensive strands of r-chromatin, which— despite its immense linear length—is compactly folded into nanostructured subdomains or nucleolar components (NCs). These include both non-nucleosomal transcriptionally active r-chromatin and nucleosomal, condensed chromatin comprising the nucleolus-associated chromatin (NAC) system [54–57, 61, 62, 65–67]. Although some NCs are visible using light microscopy (LM), its intricate organization has historically been deciphered by transmission electron microscopy (TEM) [44–67]. In TEM the nucleolus consists of five substructures: (i) fibrillar centers (FCs), regions of low electron density involved in rDNA transcription initiation; (ii) the dense fibrillar component (DFC), which surrounds FCs and mediates early rRNA processing; (iii) the granular component (GC), which contains maturing pre-ribosomal particles; (iv) the NAC system, which includes perinucleolar (PCC) and intranucleolar (ICC) condensed chromatin; and (v) nucleolar vacuoles (NVs), which appear in LM as optically clear regions are filled with nucleoplasm and must be distinguished from large FCs by TEM. Among these, NAC plays a pivotal role in integrating chromatin architecture and the nucleolar transcriptional apparatus. The PCC shell and ICC cords together form a spatially unified system that supports the compartmentalization and redistribution of NCs under physiological and stress conditions [61, 62].

As a biosynthetic and structural hub, the nucleolus is a key cellular sensor, whose morphology and organization adapt in response to differentiation, metabolic shifts, and genotoxic stress. Numerous studies have exploited the nucleolus as a model for understanding r-gene expression, r-chromatin plasticity, targeting nucleolar micro and nanostructure in processes such as proliferation, mitosis, neoplastic transformation, and cellular senescence. Morphological changes—such as nucleolar segregation, NAC compaction, and nucleolar shrinkage—serve as reliable indicators of cellular status under both normal and pathological conditions [57, 58, 61, 62, 65–80]. The best-characterized nucleolar stress response occurs during chemical inhibition of rRNA synthesis, such as treatment with actinomycin D (AMD), which leads to classical picture of nucleolar segregation/nucleolar capping, that manifests in pronounced contraction of NAC and transformation of FC/DFC complexes into crescent-like nucleolar caps. These alterations have been demonstrated using live-cell imaging, post-fixation immunodetection, and ultrastructural analysis [57, 61, 62, 68–70, 72].

In contrast, the effects of ionizing radiation—particularly γ-rays—on nucleolar architecture and dynamics remain much less understood. Given the high density of rDNA and the extended size of r-chromatin arrays, the nucleolus constitutes a vulnerable target for irradiation-induced rDNA damage [81–94]. However, it is unclear whether the damage inflicted by γ-irradiation initiates the same structural response observed with chemical inhibitors or induces an alternative pathway of nucleolar remodeling. In this context, the most intriguing question was: whether irradiation damaged NAC retains contraction ability, thus playing a crucial role in the re-localization of

FC/DFC complexes [61]. Moreover, the role of nucleolar proteins in mediating these responses remains an open question, with particular interest to Pol I associated architectural transcription factor UBTF and pre-rRNA early processing factor fibrillarin, in this context [61, 62, 64, 95–97].

To address this gap, we analyzed the nuclear and nucleolar response to 30 Gy γ-irradiation in HeLa cells permanently expressing histone H2B-GFP. This cell model has been previously validated for studying nuclear chromatin behavior during apoptosis and MC [8, 11, 39, 40, 98]. We employed live-cell 2D time-lapse microscopy and fixed-cell volume imaging over a 72-hour period, in combination with immunodetection of UBTF and fibrillarin, to monitor the temporal dynamics of nuclear deformation, multinucleation, and nucleolar reorganization.

Our goal was to determine whether γ-irradiation induces classical nucleolar segregation and whether NAC, despite sustaining DNA damage, retains its contractile ability and continues to mediate the positioning of FC/DFC assembly [61]. While we expected nucleolar behavior similar to that observed with AMD, our findings revealed an entirely distinct pattern. Although UBTF- and fibrillarin-positive domains underwent substantial rearrangement, the characteristic nucleolar caps were never formed. Instead, the nucleolus maintained its enlarged and irregular shape, while FC/DFC complexes remained integrated with the compacted NAC system. These results suggest that γ-irradiation provokes a non-canonical, yet structurally cohesive nucleolar stress response, possibly reflecting the resilience of nucleolar chromatin organization under ionizing damage.

## 2. Materials and Methods

### 2.1. Cell Culture

As a widely used model for studying cell death-related morphology, we utilized cultured HeLa cells stably expressing the histone H2B-GFP fusion protein, which was obtained courtesy of Prof. O. Piot (University of Reims Champagne-Ardenne, France). In parallel experiments, double immunolabeling of non-transfected HeLa cells with anti-UBTF and anti-fibrillarin monoclonal antibodies has been used. We used a non-transfected HeLa culture as a control, because the GFP fluorescence of the nucleoplasm could hinder the discrimination of the precise intra-nucleolar position of the anti-fibrillarin label, which also emits green light (for details, see Section 2.5).

Moreover, the HeLa culture, expressing histone H2B-GFP, was selected for its bright, UV-resistant nucleoplasm and intra-nucleolar fluorescence, enabling extended LSM observations with minimal photo-bleaching for up to 72 hours. This resistance proved crucial for the observation of living cells. Notably, the nuclear chromatin and NAC exhibit early apoptosis-associated changes, making this culture suitable for cellular stress experiments induced by chemical and/or physical factors [61, 62, 74, 98].

By targeting the entire nucleoplasm and NAC, which express histone H2B-GFP fluorescence indicative of condensed chromatin, we investigated structural modifications during 72-hour acquisition of 2D-time series. Hence, the choice of the histone H2B-GFP permanently transfected cell line was based on its specific features, namely: (i) these cells demonstrated high stability of green fluorescence in the nucleoplasm, crucial for maintaining a satisfactory level of brightness during up to 72 hours of time-lapse imaging of γ-irradiated cells. This characteristic was vital for our study; (ii); these cells exhibited prominent intra-nucleolar fluorescence, identified as nucleosomal domains with the ultrastructural appearance of intranucleolar condensed chromatin (ICC); (iii) HeLa cells are known for their prominent FC/DFC assembly [61, 62] that facilitates discrimination of nucleolar sub-territories involved in r-gene transcription as well as pre-rRNA processing while using phase contrast and fluorescent regimes; (iv) in addition to study the nucleolar behavior the model employed provides the possibility to follow dynamics of multi nuclear cell formation – the cellular pathology process which is typical for γ- irradiation induced MC.

### 2.2. Cell Maintenance

Stock cultures were maintained in 40 ml flasks containing DMEM (Gibco, UK) supplemented with 10% calf serum and 1% penicillin/streptomycin mixture. Regular reseeding occurred 2-3 times per week based on the cell monolayer reaching late pre-confluent or early confluent stages. Monthly tests for mycoplasma detection were conducted. To study nucleolar dynamics during γ- irradiation-induced remodeling, we prepared samples in a way that ensured cultures remained sub-confluent even after 72 hours of post-irradiation incubation and/or post-irradiation acquisition of living cell images.

### 2.3. Cell Seeding Methods

Method (1): The collagen-coated glass surface of Ø35 mm “MatTek” Petri dishes, featuring 14 mm wells as the growth area (MatTek, USA), was pre-conditioned by pouring a few drops of media at 37°C for 15-30 minutes. The exact amount of meticulously homogenized cell suspension was added. Cell density and distribution were assessed using a phase-contrast microscope after incubation at 37 °C for 15-30 minutes. The satisfactory cell density within the growth area was achieved by dilution, which yielded ∼10,000-15,000 cells/mL. Medium (1.5-2 mL) was added by pouring it onto the wall of the Petri dishes to avoid disturbing the attached cells. Cells can be repeatedly re-homogenized within the well using a 1.0 mL pipette or syringe. Using this concentration, we regularly achieved a dispersion of ready-to-work cultures after 48 hours of incubation. Following these conditions, we obtained cultures showing attached and flattened cells gathered in small groups, distributed sparsely enough to remain sub-confluent after 72 h.

Method (2): This method yielded a less dispersed pattern, yet suitable for maintaining sub-confluence after 72 hours. The collagen-coated glass surfaces were pre-conditioned, as described above. 1.5-2.0 mL of a properly homogenized cell suspension (∼10,000-15,000 cells/mL) was poured into wells. Desired dispersion was achieved by agitation for 5 minutes. Repeated resuspension using a 1 mL pipette or syringe provided better dispersion. Both methods ensured the availability of isolated cellular groups suitable for post-irradiation incubation and imaging.

### 2.4. Irradiation Setup

The irradiation procedure was carried out at the Center for Applied Research of the E. Andronikashvili Institute of Physics at Tbilisi State University (Tbilisi, Georgia). All experimental procedures involving γ-irradiation exposure of cells were approved by the Ethics Committee of the I. Beritashvili Center for Experimental Biomedicine and were conducted in strict adherence to Georgian National Health and Safety Regulations.

#### Selection and Rationale for Radiation Doses

In a recent study, we tested γ-irradiation doses of 10 and 30 Gy, which were deliberately chosen based on clinically relevant radiotherapy paradigms [99]. The 10 Gy exposure aligns closely with high-dose fractions commonly applied in Stereotactic Body Radiotherapy (SBRT), enabling the examination of MC and apoptosis-mediated DNA damage, repair mechanisms, and cell survival pathways representative of therapeutic scenarios. After 24-48 hours of incubation, the cells were briefly rinsed with PBS (three times for 5 minutes) and then prepared and submitted for γ-irradiation by incubating in fresh medium at 37°C for 2-3 hours. In corresponding experiments, we observed that a higher dose of 30 Gy, which is beyond typical clinical fractionation schedules, produced a much more profound impact on cellular and nuclear morphology caused by MC. Therefore, in this study, we selected to evaluate the potential utility and effects of high dose single fraction exposure to test whether EMC occurs.

#### Irradiation Procedure and Protocol

Cell cultures were delivered to the irradiation facility (irradiation chamber) in a handmade plastic container. Immediately following irradiation, Petri dishes were removed from the irradiation chamber and transported promptly in a regular air atmosphere to the laboratory incubation facility, using the same thermostatic container. The transport duration ranged between 45 minutes and 1 hour.

Cells were exposed to γ-radiation using the GUPOS-3M gamma irradiation system, which employs a fully shielded lead chamber containing a circular arrangement of ^137^Cs rods, providing a uniform gamma radiation field at a dose rate of 1.1 Gy/min. The internal temperature of the radiation chamber was maintained at a stable 25°C under ambient oxygen atmosphere conditions throughout the procedure. Before being submitted to γ-irradiation, cells were briefly rinsed with PBS (3 × 5 min), immersed in fresh media, and the dishes were delivered to the GUPOS-3M γ- installation. Irradiation of cells was conducted directly in Petri dishes (Fig. 1). Isotope ^137^Cs, which provides a dose of 1.1 Gy/min, was utilized as the source of γ-irradiation. Each irradiation procedure was carried out individually. Single Petri dishes containing cultured HeLa cells were sequentially placed in the irradiation chamber. Each dish was irradiated independently, receiving a total exposure of 30 Gy. The exposure time was precisely controlled so that to reach a 30 Gy dose, we treated cells for 33 minutes to deliver the correct cumulative dose according to the established dose rate. Each exposure protocol experiment was repeated in triplicate.

**Figure 1.**
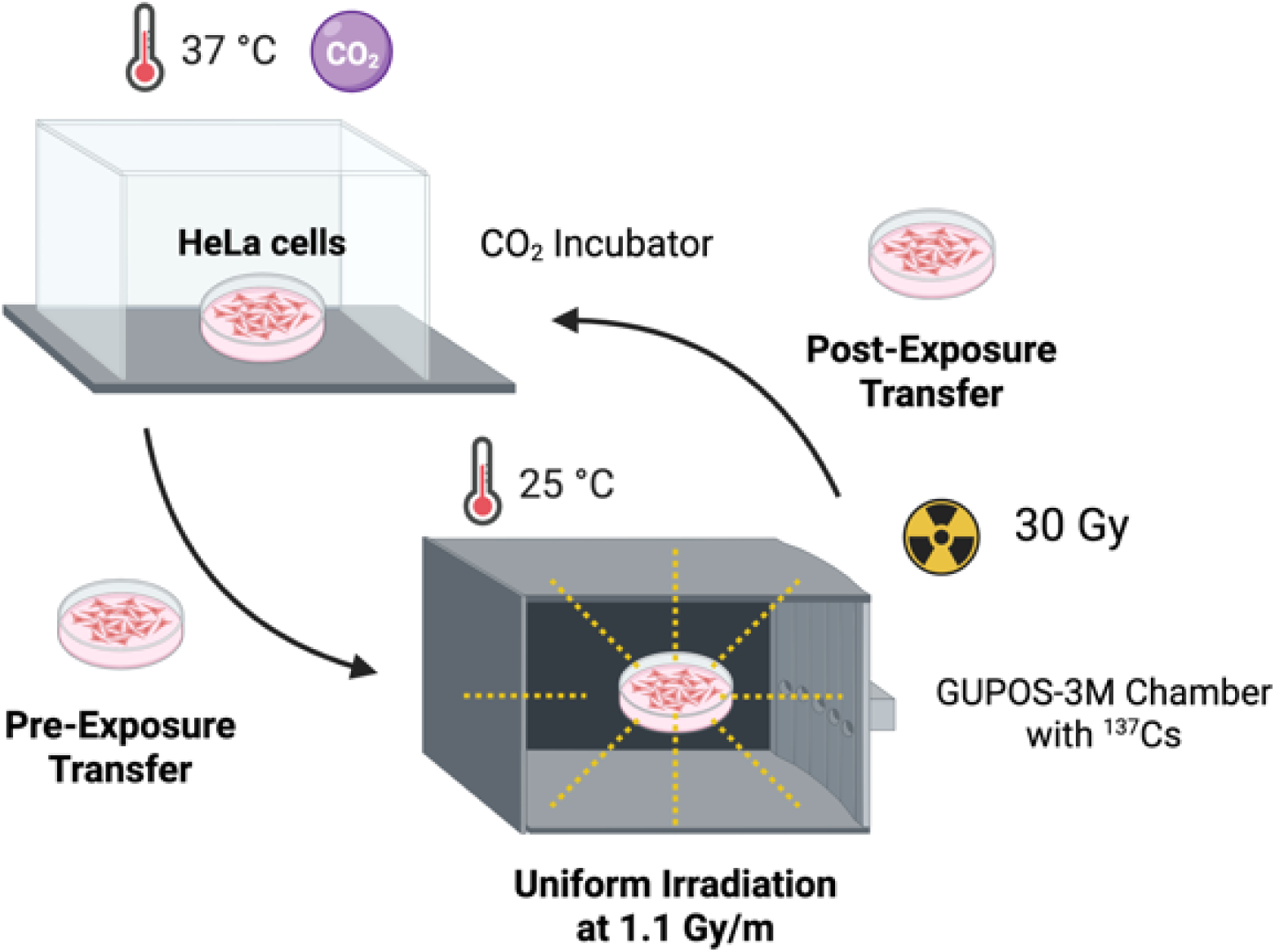
Schematic of the GUPOS-3M γ-irradiation system at the Center for Applied Research, E. Andronikashvili Institute of Physics, Tbilisi State University. The fully shielded lead chamber houses a circular arrangement of ^137^Cs rods, providing a uniform γ-radiation field at 1.1 Gy/min. Cells cultured in Petri dishes were irradiated individually, each receiving a total dose of 30 Gy.

#### Post-Irradiation Incubation and Control Samples

Upon arrival, cells were immediately transferred into a CO incubator, where they were maintained in fresh media under standard cell culture conditions for 30 min. Next, the irradiated cultures were subjected to subsequent microscopic examination and post-irradiation imaging over 72 hours [99]. While the γ-irradiated culture was passing through LSM 2D time-lapse acquisition, the resting part of the irradiated cells were maintained at 37 °C in a CO2 incubator for 72 hours. Fixed after 12, 24, 48, and 72 hours, these γ-irradiated but not laser-beam-exposed samples served as conventional morphological controls for comparison with corresponding points in post-irradiation LSM imaging. In addition, non-irradiated HeLa cells were imaged over 72 hours using the same LSM conditions as those applied for time-lapse visualization of γ-irradiated samples. As a result, during imaging at 1.5% of laser power, no structural changes on the level of cytoplasm and nucleus/nucleolus were detected.

As the temperature in the irradiation chamber was maintained at 25°C, control tests were necessary to assess any potential impact of a lower temperature on cellular and nuclear/nucleolar morphology. Meanwhile, preliminary control tests, conducted using non-irradiated cultures that were exposed to 25^°^C for 120 min, didn’t reveal (at least on the microstructural level) remarkable structural changes that could be attributed to signs of cellular metabolism cessation.

### 2.5. Post-Fixation Immunolabeling of UBTF and Fibrillarin

Imaging of UBTF-labeled cells is widely used to identify under-condensed active, potentially active, and inactive rDNA genes folded into the structure of interphase FC and mitotic NORs [61, 62, 95–97]. Meanwhile, imaging of anti-fibrillarin labeled cells was performed because it is present at high concentrations within the DFC, where its rRNA methyltransferase activity is required for rRNA processing [61, 62]. These characteristics allowed us to follow FCs and DFCs as well as to study their 3D modification induced by γ-irradiation. For post-fixation imaging of anti-UBTF and anti-fibrillarin mono- and double-immunolabeled cells, we used primary monoclonal antibodies conjugated with AlexaFluor 488 and AlexaFluor 594, both purchased from Santa Cruz Biotechnology (USA). This type of antibody enables mono-immunolabeling, allowing for the use of only one block for nonspecific binding in normal goat serum (NGS, Novex, USA), as well as avoiding incubation with a biotinylated secondary antibody through mono- and double-immunolabeling, thereby making the duration of the immunostaining procedure notably shorter. Testing primary monoclonal antibodies conjugated with AlexaFluor594 (anti-UBF/F9 fragment and anti-fibrillarin) and AlexaFluor488 (anti-fibrillarin), we developed two protocols that differ significantly from our earlier used scheme [61].

By mono-labeling both, anti-UBTF as well as anti-fibrillarin protocols included four similar steps, namely: (i) after brief rinsing in PBS γ-irradiated samples were fixed at room temperature (RoT) during 10 min in 4% PAF diluted in PBS and adjusted to pH 7.2-7.4 by 0.1N NaOH and rinsed repeatedly in PBS (3x5 min); (ii) cells were permeabilized by incubation in 1% TritonX- 100 diluted in PBS during 5 min, and extensively washed in PBS; (iii) to block nonspecific binding, cells were incubated in 10% NGS in PBS during 60 min at RoT; (iv) after removing NGS the cells were covered either by mouse anti-UBF/F-9 fragment or by mouse anti-fibrillarin AlexaFluor594 conjugated primary antibodies diluted (1:20) in PBS containing 1% NGS overnight at 4°C. After cells were washed with PBS (3 × 5 min), selected samples were subjected to LCM imaging according to the procedure described in Section 2.4.

For the simultaneous visualization of UBTF (in red) and fibrillarin (in green), we utilized only intact, non-transfected cultures. Double labelling was performed according protocol that includes following 6 steps: (i) as it was described above, briefly rinsed γ-irradiated cells were fixed in 4% PAF and rinsed again; (ii) cells were permeabilized in 1% TritonX-100 prepared in PBS during 5 min, and extensively washed in PBS; (iii) to block nonspecific binding, cells were incubated in 10% NGS in PBS during 60 min at RoT; (iv) after removing NGS the cells were covered with anti-UBF AlexaFluor594 conjugated primary antibodies diluted (1:20) in PBS containing 1% NGS overnight at 4°C; (v) after rinsing in PBS (3x5 min) cells were secondly incubated in 10% NGS during 60 min at RoT; (vi) NGS was removed and cells were covered with anti-fibrillarin

AlexaFluor488 (1:20) dissolved in PBS containing 1% NGS over night at 4°C. After washing in PBS, the cells were subjected to LCM imaging directly in their culture boxes.

### 2.4. 2D Time-Lapse 2D Imaging and 3D Visualization using ZEN software

After irradiation, cells were washed in PBS (three times, 5 min each), returned to the cultivation medium, and then subjected to time-lapse imaging over 72 h using a Carl Zeiss (Germany) LSM 900 microscope (hereafter referred to as LSM), equipped with an AxioObserver Z1/7 inverted microscope and an AiryScan 2 augmented resolution device. All procedures, including post-irradiation behavior observation, were conducted directly on cultures in Petri dishes. The imaging was performed at 37°C in a CO_2_-enriched atmosphere using a special microscope plate holder with a sealed box. Microscopy details included using various objectives at optical zooms of 100x, 200x, and 630x. Hence, images were registered at low and high magnifications using objectives with corresponding numerical apertures. We used Plan-Neofluor/10x with 0.3 M27 numerical aperture and Plan-Apochromat/20x/0.8 M27 for low magnification, including a time-lapse approach. By imaging objects at high magnification, including time-lapse mode, we used Plan-Apochromat/63x/1.4 Oil DIC M27. The time-lapse imaging setup involved capturing images every 10 minutes for every 12-24 h post-irradiation imaging time (i.e., during the first 12 hours, 12-24 hours, 24-48 hours, and 48-72 hours), simultaneously in phase contrast and fluorescence regimes.

Histone H2B-GFP fluorescence was induced and recorded using a 1.5-2.5% Colibri 7 laser device at 488 nm excitation light, while emission light was 517 nm. As a control, we used high-magnification LSM imaging conducted before γ-irradiation treatment. For time-lapse analysis and visualization of the behavior of living cells damaged by γ-irradiation, the obtained time series were transformed into 2D movies using Carl Zeiss ZEN 3.0 software (Germany). Significant points from the dataset were extracted to visualize changes in nuclear/nucleolar quantitative and structural parameters, as well as the movement and coalescence of ICC clumps during γ-irradiation.

For fibrillarin immunodetection, using anti-fibrillarinAlexaFluor488 monoclonal antibodies, fluorescence was induced by 488nm wavelength of 2% of laser power, while emission light was caught at 517nm. Meanwhile, for visualization of both UBTF and fibrillarin using AlexaFluor594 conjugated anti-UBF/F-9 fragment and anti-fibrillarin monoclonal antibodies, fluorescence was excited at 561nm with 2% laser power and an emission wavelength of 618 nm.

By virtual serial sectioning necessary for 3D visualization and following volume reconstruction, corresponding Z-stacks comprising whole nuclear volume were collected at medium scanning speed, using interlayer distance (section plan thickness) of 0.3 µm. Related 3D models of living as well as post-fixed and immunolabeled interphase and mitotic cells were generated by means of ZEN 3.0 software using solid surface and/or transparent volume rendering options.

### 2.5.Image 3D Reconstruction and Rendering using “UCSF Chimera” software

3D Reconstruction and modeling were also performed by means of “UCSF Chimera” (Computer Graphic Laboratory, USA) software in following way. Acquired using LSM Z-stacks of GFP-tagged chromatin, anti-UBTF/AlexaFluor594 labeled and anti-fibrillarin/AlexaFluor488, 594 labeled samples were exported from ZEN 3.0 as 16-bit single-channel TIFF files that retained voxel-size metadata. Stacks were opened in “Fiji/ImageJ” via Bio-Formats, split by channel, and background-corrected. Slice-specific binary masks were generated with the “Otsu” threshold algorithm in stack mode (Image [Z Adjust [Z Threshold; Method = Otsu; Black background). Resulting masks were saved as TIFFs for each channel. Threshold adjusted stacks were loaded into “UCSF Chimera” software with Volume Viewer. The surfaces were created at density 0.5 (appropriate for binary data), pseudo-colored (chromatin - green, UBTF - red, fibrillarin - cyan), and smoothed with three iterations of Gaussian filtering (kernel = 1 voxel). Uniform lighting, 35% transparency on the chromatin surface, and orthogonal clipping planes were applied across all samples. High-resolution stills (2,000 × 2,000 px, 600 dpi) and tilted pictures were exported directly from “Chimera” software. All threshold and rendering parameters were kept identical for every condition to ensure comparability. By 3D rendering UBTF-positive structures of our interest (SOI, hereafter) were always red, while by rendering of anti-fibrillarin labeled SOI, we either kept original red color or changed it to cyan.

For better visualization of NAC system, we utilized AutoCAD 2021 (Autodesk, USA). The LSM Z-stack images were exported in .jpeg format and orientation markers were added in Photoshop 2020 (Adobe, USA) to ensure accurate overlay of layers. Structures within each image were manually outlined in CorelDRAW 2019 (Alludo, Canada) and exported in “.dwg” format for compatibility with AutoCAD. The outlined structures were then inserted into separate AutoCAD layers, sequentially aligned, and elevated along the Z-axis in proportion to the original 0.3 µm interlayer difference. Surface modeling was achieved using the “mesh” and “loft” tools. The visual style was switched to “Realistic” to enhance visualization of the structures by adding transparencies. The models were rendered at Full HDTV resolution and saved in .png format with 32-bit color depth (24-bit RGB + 8-bit alpha channel) at 300 DPI. In the final stage, the processed images were overlaid onto the original LSM images to ensure accurate alignment with the corresponding structures. This workflow enabled the generation of accurate and visually detailed 3D representation of the NAC and other SOI.

## 3. Results

### 3.1. Native and Fixed NAC Integrates ICC Network and PCC Shell into 3D Unit

Experiments were started by LSM imaging of nuclei and nucleoli in control HeLa cells stably expressing histone H2B-GFP (SFig. 1 – 4). While SFig. 1 shows fixed cells, images, displayed in SFig. 2 – 4 reveal the nuclear structural organization in living cells taken at low (SFig. 2) and higher magnification (SFig. 3). At the same time, SFig. 4 demonstrates spatial structure of nuclei and chromosomes after 3D reconstruction. The intensity of fluorescence in different regions of chromatin outside the nucleolus varies depending on the compaction degree, while the inter- chromatin compartment remains unlabeled. The brightest, i.e., the most compact regions housing histone H2B-GFP, correspond to chromocenters and PCC shell. Consequently, brightly fluorescent NAC that comprise PCC, and ICC, belonging to SOI are clearly featured. Typically, 2D observations of living and fixed samples reveal mononuclear cells, as shown by the merging of phase contrast (PC) and fluorescent images (SFig. 1, b). Abundant dividing cells at various stages of mitosis, and occasional apoptotic cells were observed (SFig. 1, a, c, d– 3, d, e, 4).

**Figure 2.**
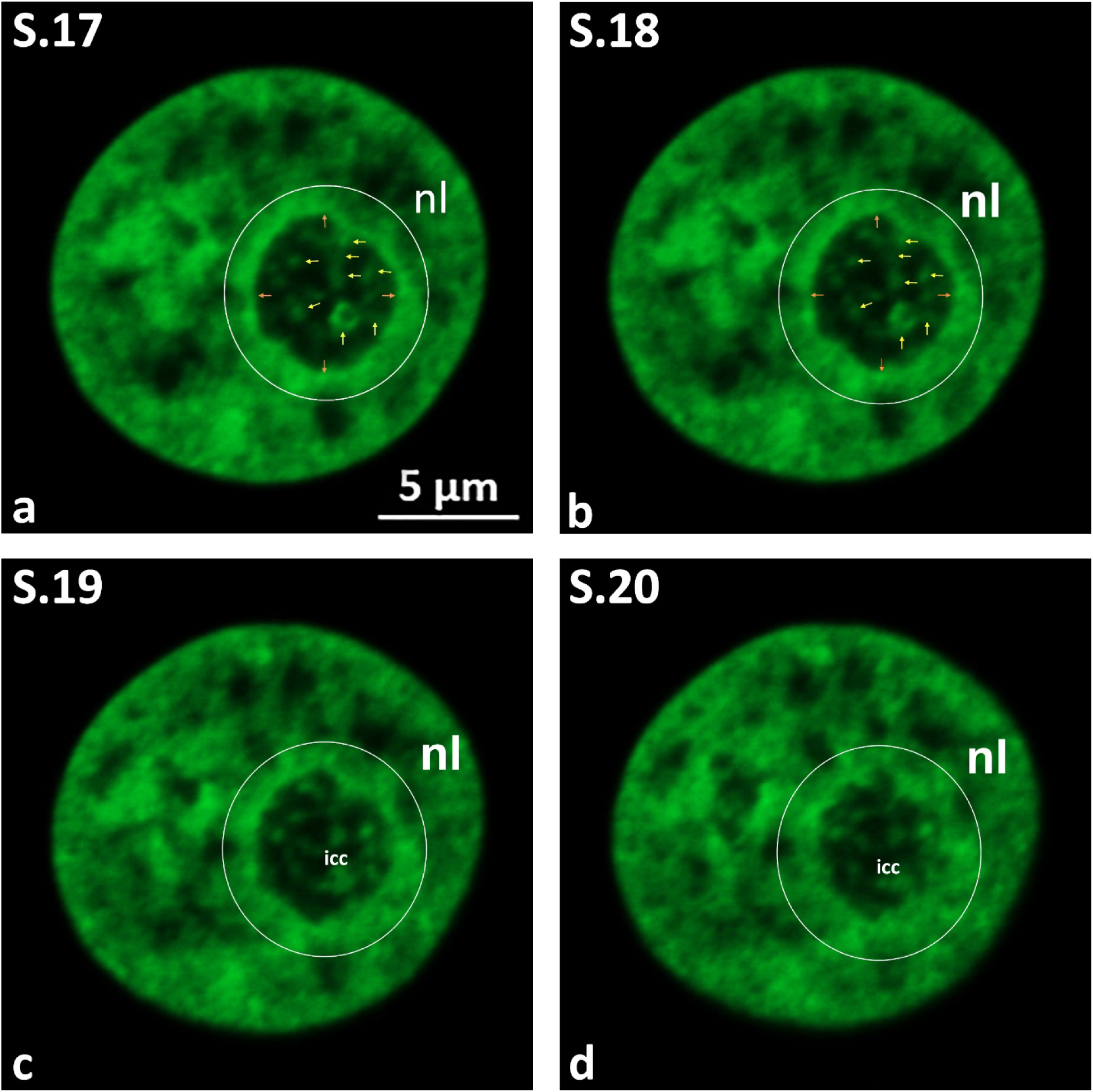
Control: Microstructure of nucleus and nucleolus in HeLa cells stably expressing histone H2B-GFP. (a–d) Successive virtual serial sections (S.17–S.20) from a complete Z-stack, showing nucleoli as large, dark areas against brightly fluorescent chromatin. Note the GFP- positive PCC ring surrounding nucleoli and intranucleolar ICC clumps or anastomosing cords extending from PCC, forming a network-like 3D structure. A large ICC clump displaying a ring- shaped structure is visible (a, b). PCC – perinucleolar condensed chromatin; ICC – intranucleolar condensed chromatin).

**Figure 3.**
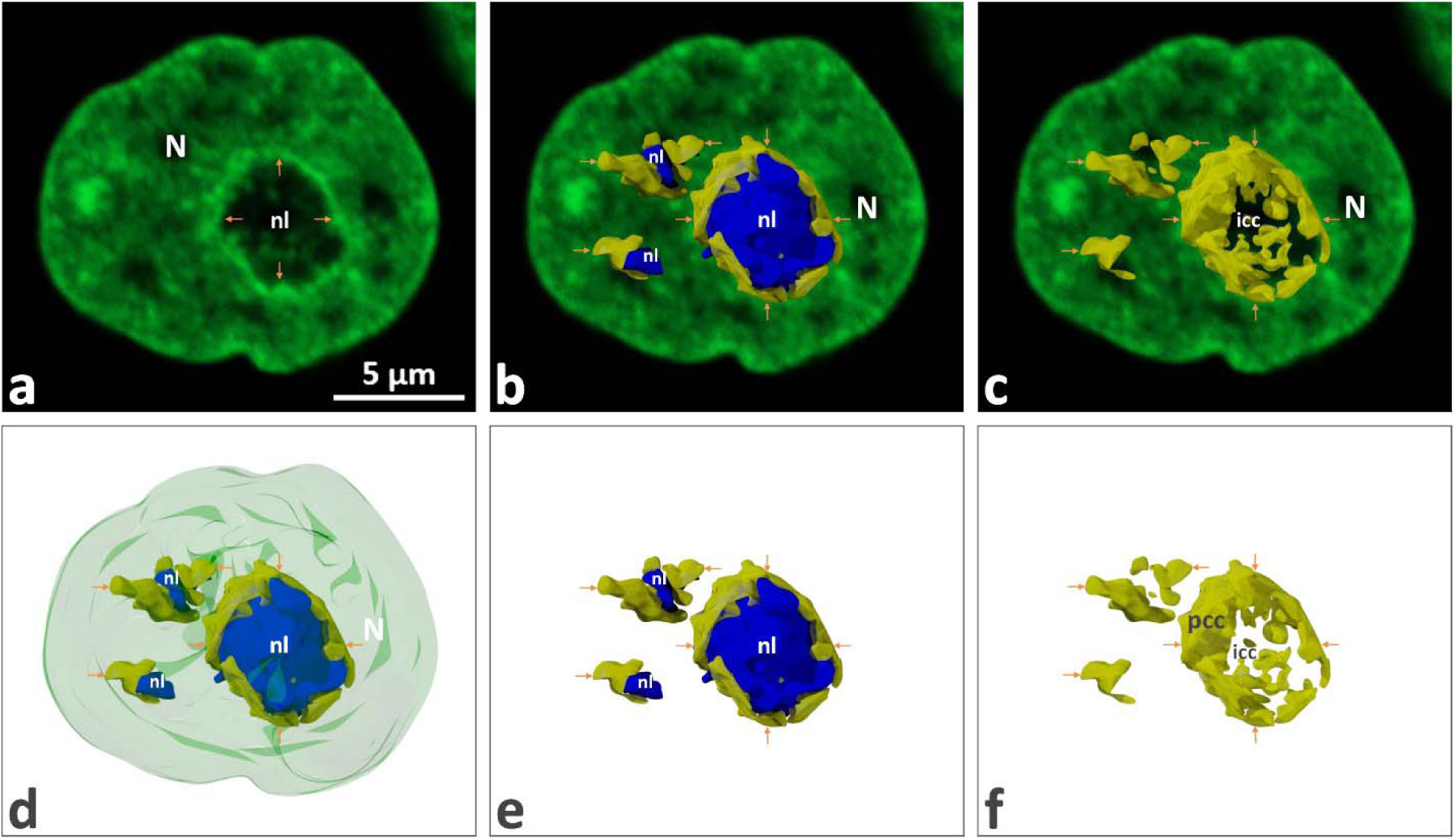
Control: 3D nucleolar model generated using AutoCAD. (a) Single 2D optical section of a nucleus (N) from a histone H2B-GFP HeLa cell. Prominent PCC ring (brown arrows) outlines the nucleolar territory; weaker fluorescent ICC cords are visible within. (b, c) Same plane merged with 3D models of nucleoli (blue, b) and NAC (yellow, c); nucleus contains three nucleoli (nl). (d–f) Sequential 3D models demonstrating spatial arrangements of the entire nucleus with nucleoli and NAC (d), nucleoli and NAC (e), and NAC alone (f). The integrated shell-like PCC and ICC network is evident. (Abbreviations as in previous figure, N – nucleus; NAC - nucleolus-associated chromatin; nl – nucleolus).

**Figure 4.**
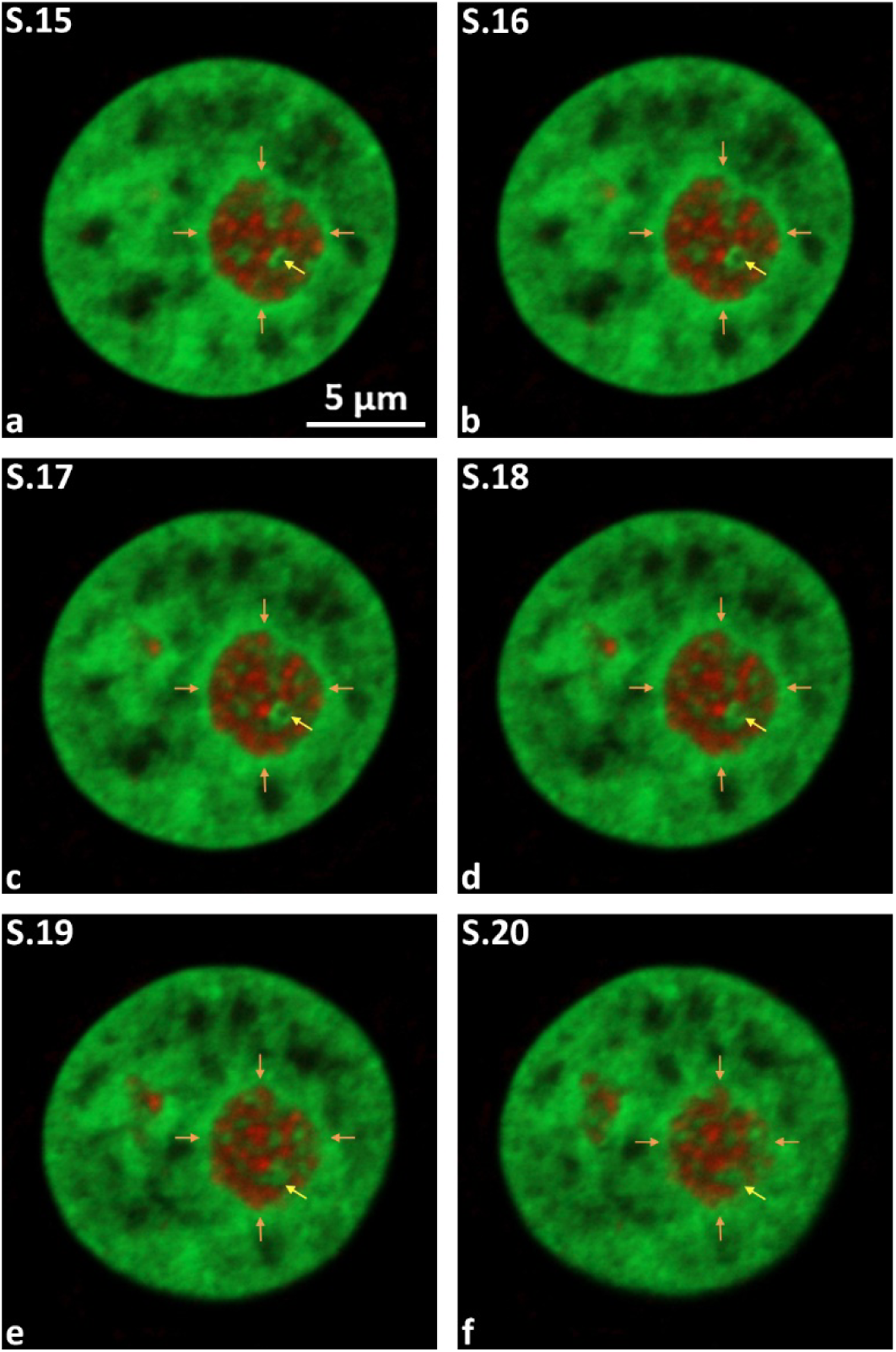
Control: Anti-UBTF immunolabeling corresponding to the nucleus in Fig. 2. Virtual serial sections (S.15–S.20, a–f) demonstrate nuclear and nucleolar microstructure. UBTF (red) is concentrated in multiple intranucleolar foci (∼0.3–0.5 µm diameter), arranged as "folded necklaces." Numerous ICC (yellow arrows) contacts with UBTF-positive entities are clearly visible. PCC ring indicated by brown arrows.

Predominantly, nuclei have a roundish or ovoid shape with slightly irregular contours well recognizable on 3D models (SFig. 4), while the average diameter after fixation ranged between ∼15 - 25 μm (Fig. 1, b).

Nucleoli appeared against the brightly fluorescent chromatin as large, dark, spherical, or elongated territories. The nucleolar interior regularly showed fluorescent inclusions as ICC clumps/cords extending from the PCC shell deep inside the nucleolus (Fig. 2; SFig. 2,3). PCC consistently exhibits higher fluorescence than ICC. The nucleolar territories are well recognizable as round, dark zones with a diameter ranging from ∼4 to 7 μm (SFig. 2, f, h). Even at low magnification (SFig. 3) clearly that nucleoli contain histone H2B-GFP-positive structures of varying sizes and appearances, attributed to ICC. In turn, ICC inclusions have slightly less intense labeling than nucleoplasm chromatin fluorescence. These features become evident when analyzing various individual or random virtual serial 2D planes at higher magnification (Fig.2; SFig. 3, d, e). Corresponding images display the 2D structure, sizes, and localization of ICC zones, which are consistently present in all cells and vary in size and prominence. As a consequence, ICC inclusions were appearing in individual sections as distinct clumps or extended anastomosing cords (Fig. 2; SFig. 3, d, e). Images taken at high magnification (Fig. 2) confirm the persistent presence of numerous ICC inclusions and PCC ring in all observed cells. Here, ICC is either visible as discrete entities or forms well-recognizable network-like structures. Furthermore, the analysis at higher magnification confirmed that PCC is organized into a locally open ring, ∼0.25-0.55 μm in thickness, that encircles a significant part of the nucleolar territory. In addition, these images provide definitive evidence confirming the direct physical link between the PCC ring and ICC clumps as reported earlier from observations on ultrathin sections, including those treated for preferential demonstration of NAC [62, 66–68].

In 3D, clearly visible that PCC forms around confines of the nucleolar territory, a prominent, solid, or locally disrupted shell as revealed by rotation and gradual slicing of corresponding models (Fig. 3). Moreover, 3D visualization and rotation of models reveal multiple cord-like structures emanating from the PCC shell (Fig. 3, a, c, f). Thus PCC protrude inside the nucleolar territory, creating on certain virtual sections the impression of discrete ICC clumps, while in fact these clumps correspond to sections of ICC cords. All together strongly support the existence of a united ICC network that is continuous with the PCC shell, representing protrusions of the latter (Fig. 3, a, c, f). To capture NAC in its native state, we imaged its random sections immediately before starting time-lapse acquisition, registering the PCC shell in continuity with the ICC forming a network.

When measuring the average diameter of nuclei in living HeLa cells, we noted that it’s higher than in fixed samples, ranging between 19 and 25 μm. Importantly, SOI in fixed cells exhibited fluorescence comparable to or slightly less intense than in living cells. Nucleolar sizes were also slightly higher than in fixed cells, reaching from 6.5 to 8 μm. On 2D images, living cells were rich in discrete histone H2B-GFP-positive labeling within the nucleolar territory. As expected, in 3D, ICC appeared as larger clumps or anastomosing cords that all complexed in unit network-like structures. Accordingly, we can conclusively state that the native structure of ICC is network-like. The extensively branched ICC network consists of anastomosing cord-like protrusions emanating from the PCC shell. The PCC resembles a highly compacted part of the juxtanucleolar nucleoplasm, forming a shell enveloping the large part of the nucleolar territory (Fig. 3).

### 3.2. In Control Cells, UBTF and Fibrillarin Positive NCs are Integrated into NAC Unit

Here, we refer to the 2D distribution and 3D organization of UBTF and fibrillarin-positive NCs in fixed control cells. Thus, to visualize the relationships between the NAC and FCs, we imaged immunolabeled UBTF inside H2B-GFP positive intra-nucleolar network formed by naDNA condensed chromatin branches. Like in recently imaged samples of control histone H2B-GFP transfected HeLa cells [61, 62], UBTF was solely concentrated in numerous intranucleolar foci of ∼ 0.3-0.5 µm in diameter, disposed as so-called “folded necklaces” by 3D visualization (Fig. 4-6, a; SFig. 5). The size and topography of UBTF-positive sites correspond to FCs, as confirmed by a plethora of studies, in which UBTF was stably localized exclusively in mitotic NORs (SFig. 6, 7) and in their interphase counterparts, i.e., FCs [61, 62, 95–97]. In 3D views, the spatial interplay between GFP-positive NAC and UBTF-positive NCs became especially obvious, so that an intensively branched ICC network always appeared intermingled with UBTF- positive foci (Fig. 5, 6). Moreover, ICC clumps, which were linked to UBTF-positive structures, were also attached to a PCC shell, creating a bridge between these chromatin components (Fig. 5, a, I, j, 6, b). The proximity of ICC and UBTF-positive NCs, as well as their spatial integration into the entire NAC network, confirms the early-described tight link between both naDNA fractions, belonging to: (i) relaxed r-chromatin and (ii) nucleosomal, most probably non-ribosomal chromatin [61].

**Figure 5.**
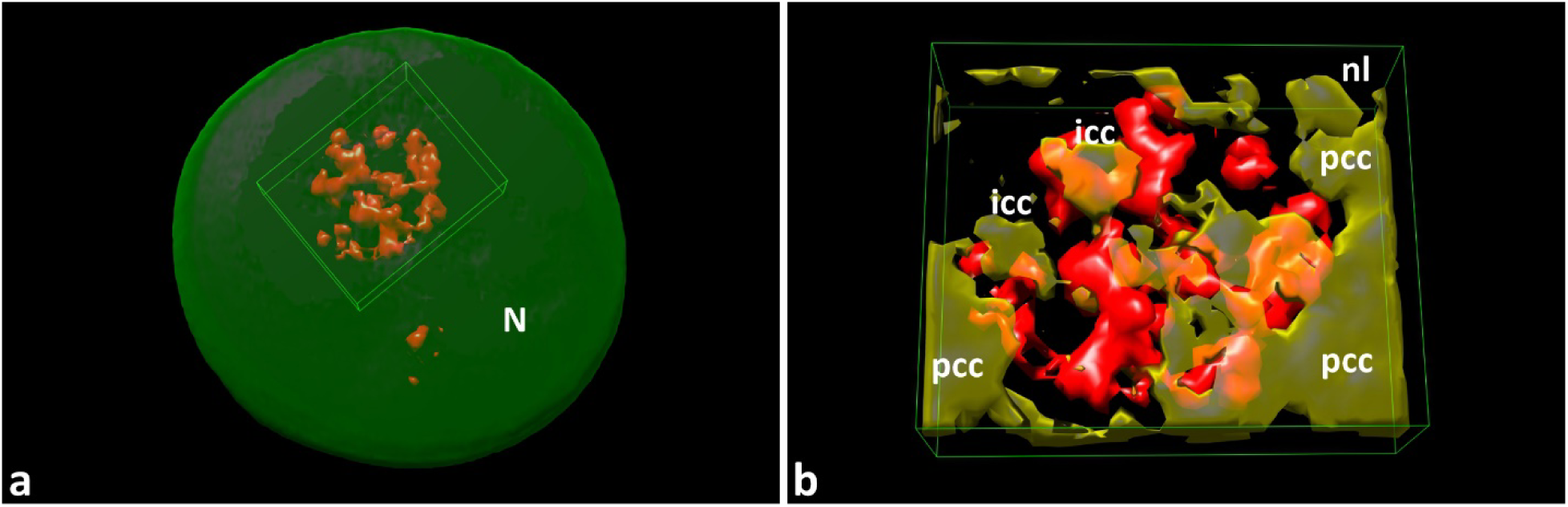
Control: 3D nuclear model following anti-UBTF immunolabeling (UCSF Chimera software). (a) Intranucleolar distribution of UBTF-positive structures (red), arranged in necklace-like chains; nucleolar territory marked by rectangle. The nucleus contains two nucleoli of distinctly different sizes; some smaller UBTF-positive structures belong to the smaller nucleolus. (b) 3D model extracted from the nuclear interior, demonstrating UBTF-positive structures integrated into the NAC system.

**Figure 6.**
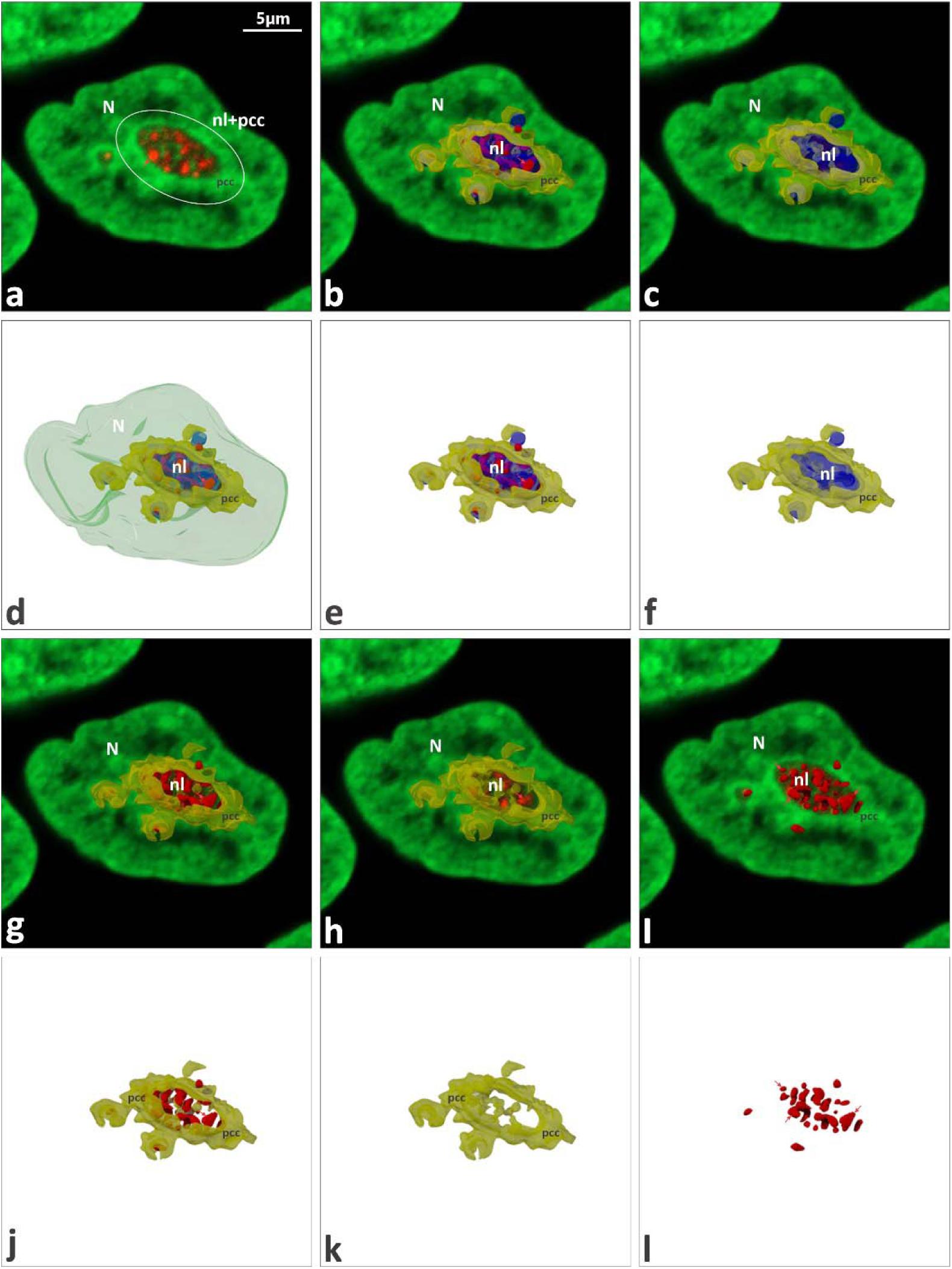
Control: Anti-UBTF immunolabeling, 3D nuclear and nucleolar models (AutoCAD). (a) Single 2D nuclear plane, showing multiple UBTF-positive entities within nucleolar territory and prominent PCC ring; nucleolus and PCC indicated by white circles. (b) Same plane merged with 3D models of nucleoli (blue), UBTF-positive structures (red), and NAC shell (yellow). (c) Same plane with nucleolus and PCC shell. (d–f) Sequential 3D models demonstrating spatial interactions of nucleus with nucleoli, UBTF structures, and PCC shell (d), nucleoli with UBTF and PCC shell (e), and nucleolus with PCC shell only (f). (g, h) Structural interaction of PCC shell with UBTF-positive structures; nucleolus sectioned at different depths for clarity. (i) 2D plane and 3D models of UBTF-positive structures, demonstrating intranucleolar UBTF distribution. Note additional UBTF signals from smaller neighboring nucleoli. (j–l) 3D models illustrating spatial interaction of NAC with UBTF-positive structures (j), NAC only (k), and UBTF-positive structures alone (l, red arrows). Clear integration of UBTF structures into NAC. Abbreviations as in previous figures.

To analyze the control 3D organization of Fibrillarin positive DFC in compliance with the spatial distribution of FCs and NAC, we applied anti-fibrillarin mono-labeling (Fig. 7-9; SFig. 8, 9) as well as simultaneous anti-fibrillarin and anti-UBTF double labeling (Fig. 10). Results obtained in these co-distribution experiments are displayed in Fig. 10. By 2D observation as well as by 3D analysis the intranucleolar fluorescence of Fibrillarin regularly appeared organized as juxtaposed curls (of ∼1.2 µm in diameter) and xtended cords (Fig, 10, a, b). The cord-like morphology of the anti-Fibrillarin labeling suggests that these entities essentially correspond to the extended DFC strands of the nucleolonema, which is an obligatory component of highly active reticulated nucleoli of HeLa cells [62]. 3D model displayed on Fig. 6 and Fig. 9 we used to confirm tight integration of UBTF- and fibrillarin-positive NCs into ICC network. The close structural link between UBTF- and Fibrillarin-positive nucleolar subdomains suggests the putative position of FC/DFC assembly within the ICC network. Concomitant 3D visualization of the couple fibrillarin and UBTF, as well as their relationship, produces clear evidence that these nucleolar proteins are closely associated. By this, fibrillarin positive DFC “covers” the necklaces-like organized groups of FCs. Therefore, inside these units, discrete FCs (or their groups) appear to be embedded in a cord-like mass of DFC that constitutes a ring-like structure around the FCs by virtual sectioning (Fig. 10, c, d). In summary, analyzing simultaneously nucleolar relative 3D repartition of UBTF-positive non-nucleosomal r-chromatin folded in structure of FC and nucleosomal condensed chromatin labeled by histone H2B-GFP we concluded: (i) ICC is always organized as an intra-nucleolar unit meshwork that interconnects FC/DFC assembly with PCC shell enveloping the nucleolus; (ii) presence of both proteins firmly indicates the proximity of ICC and FCs as well as spatial integration of FC/DFC assembly into the whole NAC in form of unit dynamic system.

**Figure 7.**
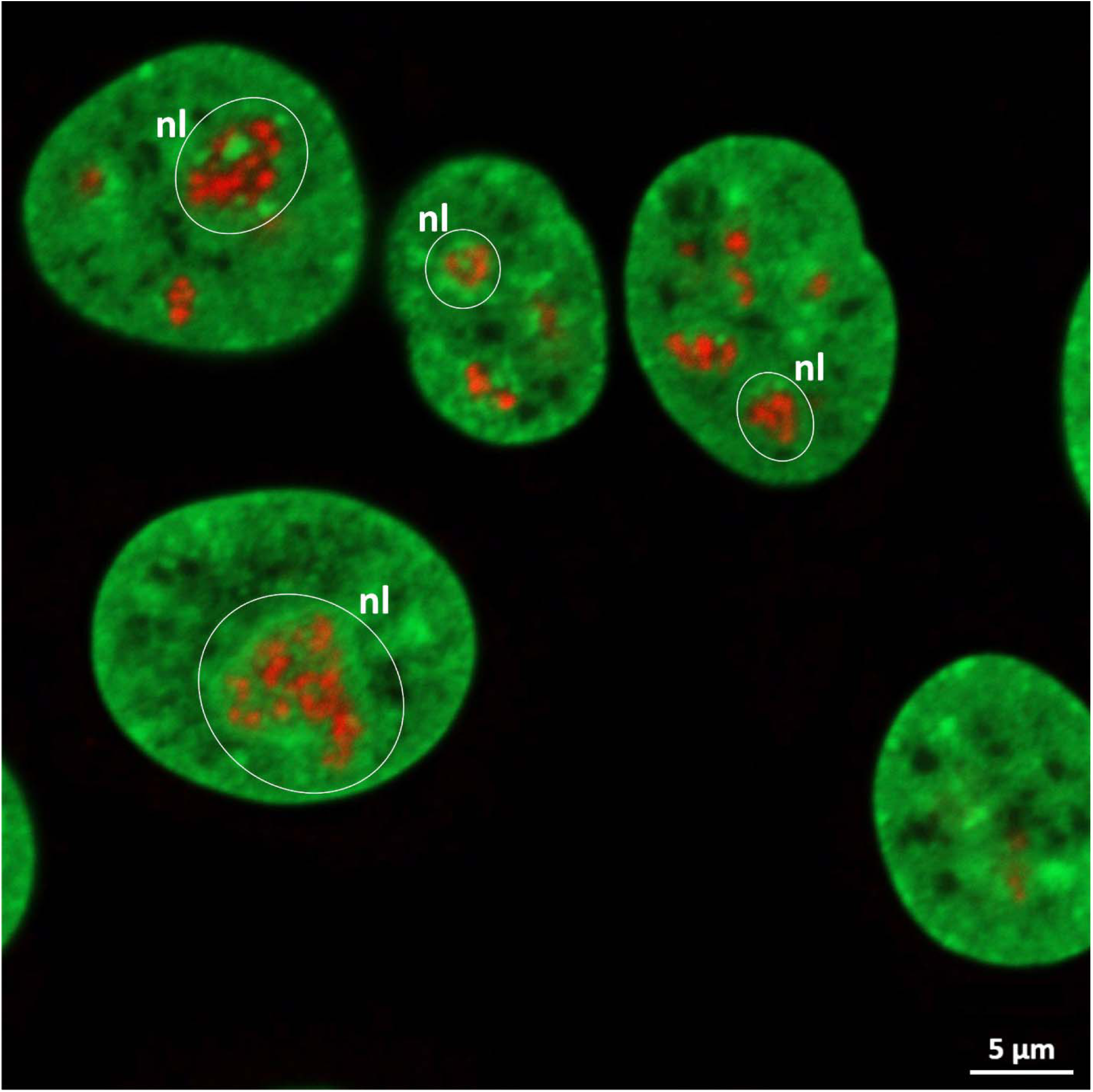
Control: Nuclear and nucleolar microstructure following anti-fibrillarin immunolabeling. Nucleolar territories are outlined by white circles. Fibrillarin (red) is localized in NCs forming extended cord-like structures.

**Figure 8.**
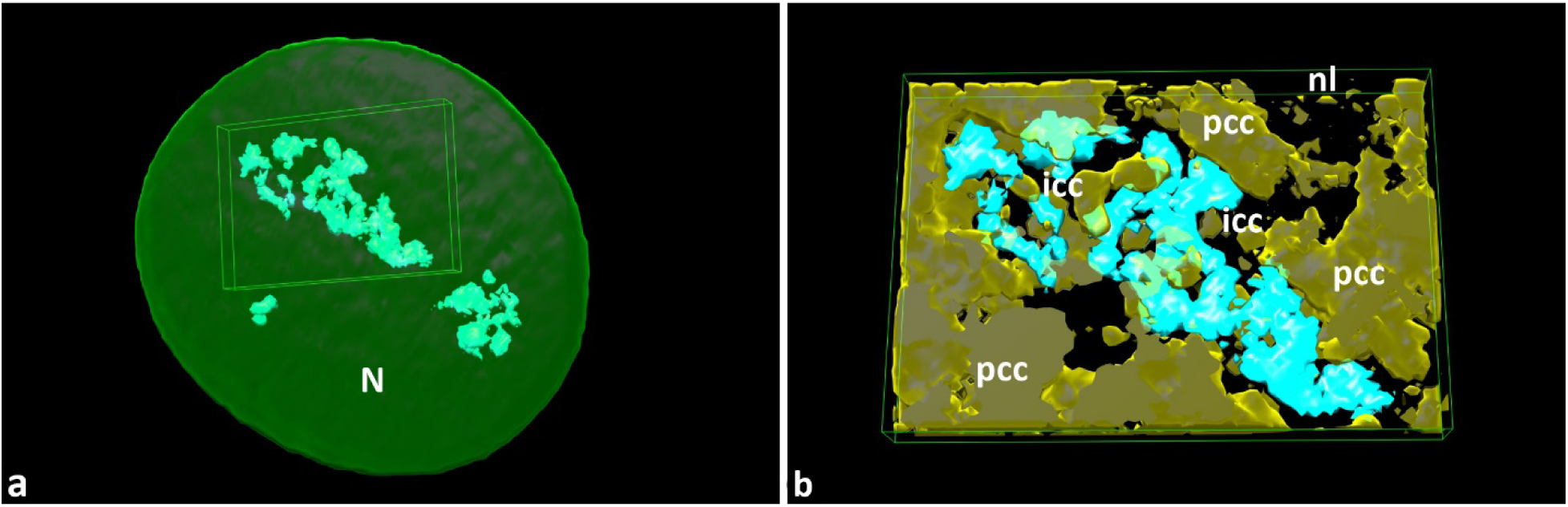
Control: 3D nuclear and nucleolar model (UCSF Chimera software) following anti- fibrillarin immunolabeling. (a) Intranucleolar distribution of fibrillarin-positive structures (cyan) displaying cord-like organization; nucleolar territory marked by rectangle. The nucleus contains three nucleoli of varying sizes; the smallest nucleolus is located lower left. (b) Extracted 3D model highlighting fibrillarin-positive cords integrated within the NAC system. Abbreviations as in previous figures.

**Figure 9.**
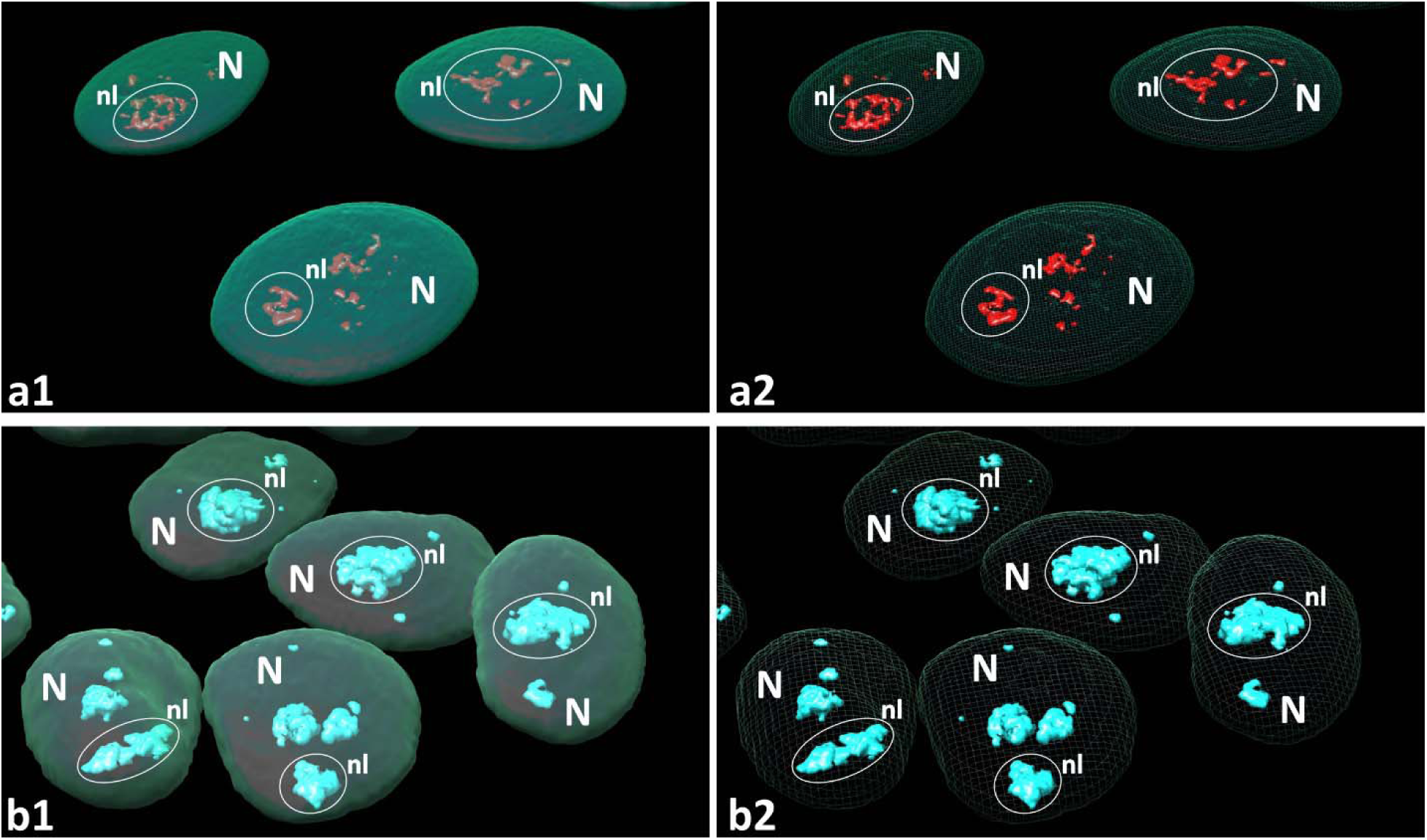
Control: 3D models (UCSF Chimera software) demonstrating spatial organization of nuclei and nucleoli after anti-UBTF (a1, a2, red) and anti-fibrillarin (b1, b2, cyan) immunolabeling. Models generated using (i) solid/transparent (a1, b1) and (ii) outlined/transparent rendering (a2, b2), enhancing nuclear volume visualization. Solid rendering alone clarified intranucleolar content. Structural differences clearly depicted: UBTF-positive structures arranged in chain-like assemblies (a1, a2), fibrillarin-positive structures appeared fused into cloud-like formations (b1, b2). Abbreviations as in previous figures.

**Figure 10.**
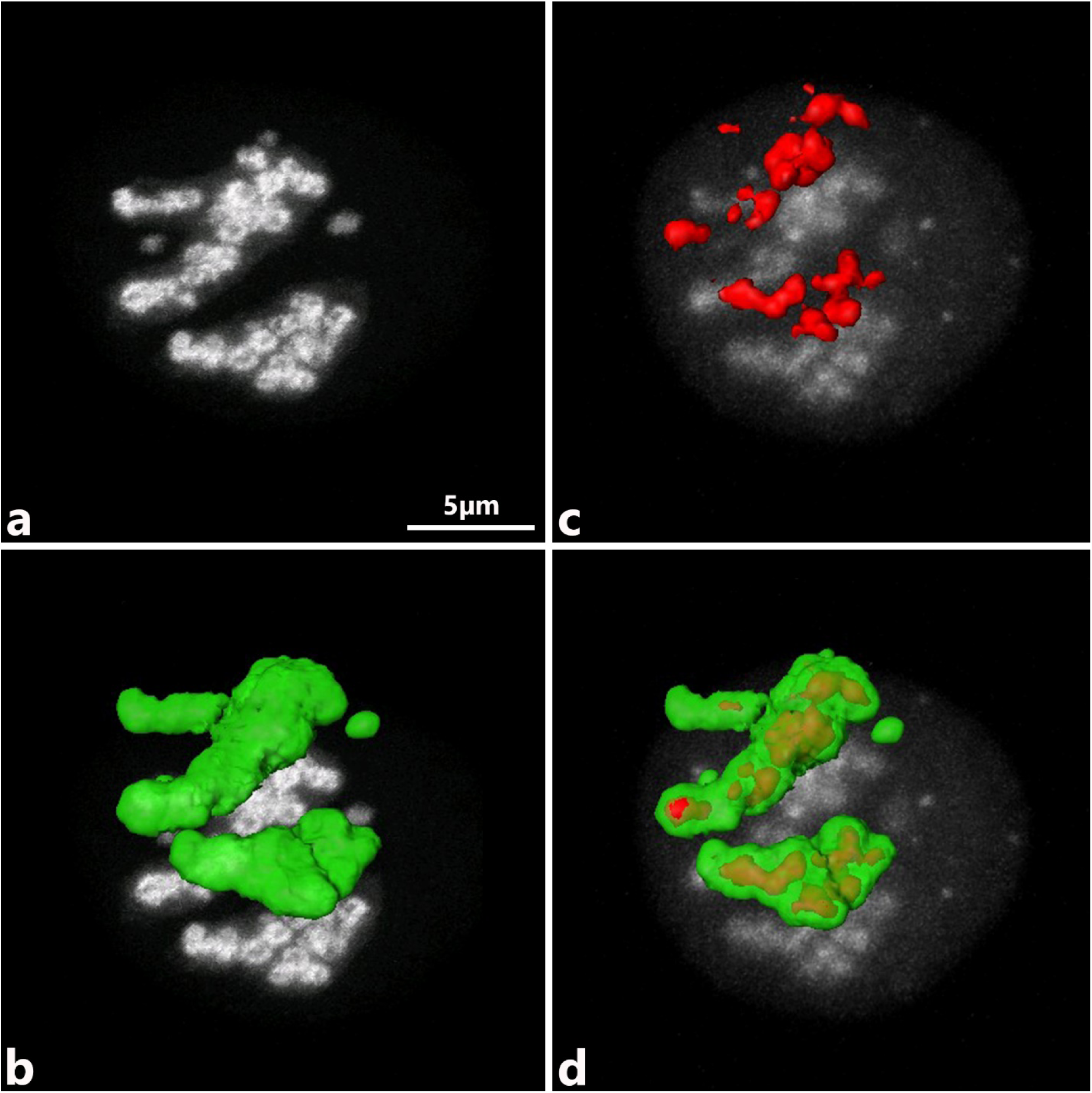
Control: Simultaneous anti-UBTF (red) and anti-fibrillarin (green) double immunolabeling in non-transfected HeLa cells, aligned with corresponding 3D models (UCSF Chimera software). Fibrillarin-positive structures show distinct cord-like 3D organization (a, b); UBTF-positive structures form necklace-like chains of similar-sized spherical entities (c, d). Fibrillarin-positive cords envelop UBTF-positive chains, forming integrated FC/DFC assemblies [61].

**Figure 11.**
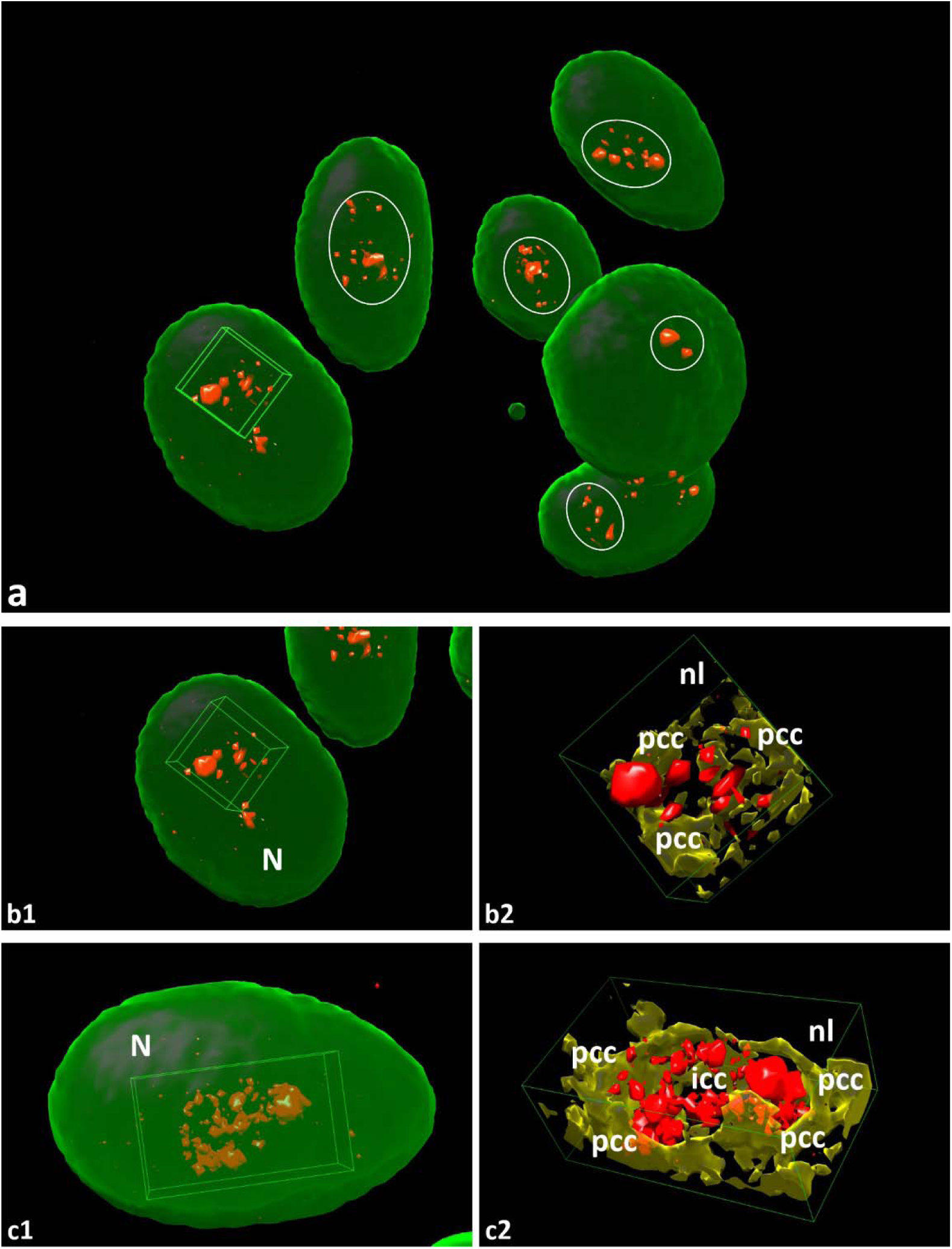
30 Gy γ-irradiation, 24 hours of post-irradiation image acquisition: anti-UBTF immunostaining with 3D nuclear/nucleolar renderings (UCSF Chimera). (a) Marked asymmetric enlargement of UBTF-positive bodies (red): one-to-three giant spheroids plus numerous smaller foci within the nucleolar territory (outlined in white). (b1, b2) Higher-magnification models show a nucleolus that contains one giant and several minor UBTF-positive bodies seamlessly embedded in the NAC shell (yellow). (c1, c2) Another nucleus with two giant and many minor UBTF bodies, again fully integrated into the NAC.

### 3.3 2D Time-lapse Imaging of Irradiated Cells: Striking Changes, delay until 48 – 72 Hours, while ICC Remains Mobile

Time-lapse imaging was conducted to show the detailed chronology of the pre-MC and pre-mortal dynamics of DNA-containing SOI under γ-irradiation with a dose of 30 Gy. This approach facilitated simultaneous imaging of whole nuclei, nucleoplasm, and nucleoli, allowing for 2D analysis of modifications over time (SMovies 1-6). Generally, after irradiation, the significant part of initially ovoid nuclei in surviving cells transforms into irregular shapes, undergoes drastic deformation or lobulation, and eventually and eventually fragments through a cleavage process followed by endomitotic division, resulting in the formation of multinuclear cells (SFig. 10-12). Likely, we did not find clues of cell division arrest before 48 hours, as different stages of mitosis (including its pathological forms) were also observed even in samples collected over 48-72 hours of the post-irradiation image acquisition period.

**Figure 12.**
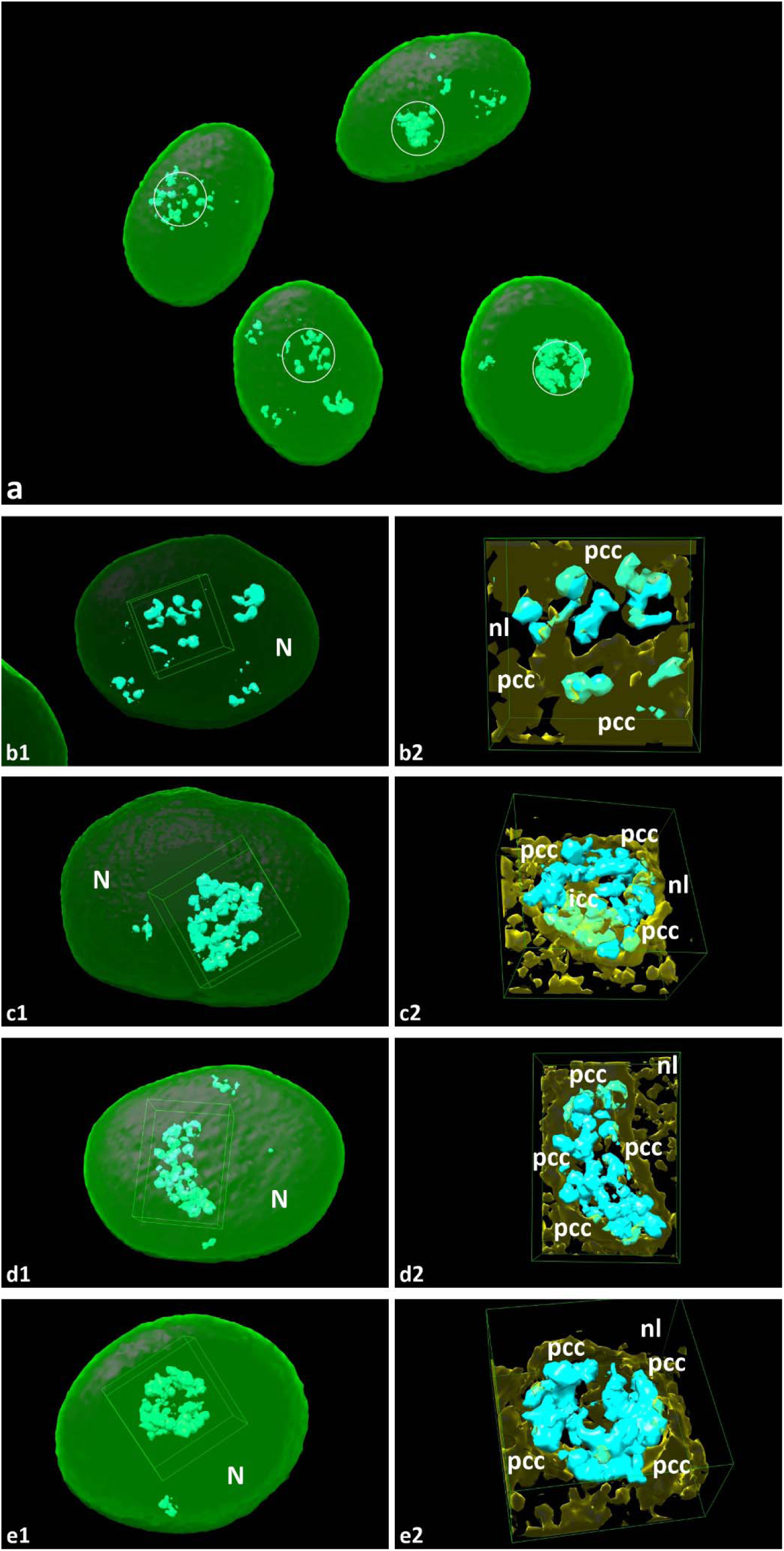
30 Gy γ-irradiation, 24 hours of post-irradiation image acquisition: mononuclear cells, 24 hours after 30 Gy γ-irradiation; anti-fibrillarin immunostaining with 3D nuclear modeling/renderings (UCSF Chimera software). (a, b1–e1) Fibrillarin-positive structures (cyan) form cord-like arrays within the nucleolar territory (outlined in white). (b2–e2) Extracted 3D views reveal these fibrillarin-positive cords seamlessly anchored in the NAC shell (yellow). Abbreviations as in previous figures.

According to the time-dependent changes developing in irradiated and surviving cells, we divided the entire post-irradiation time-lapse imaging period into four stages: stage 1 (the first 12 hours of the post-irradiation image acquisition period). In fact, stage 1 became longer due to 45 min - 1hour of post-irradiation transportation time and 30 min of irradiated cells recovery at 37 °C. The following stages, namely, stage 2, stage 3, and stage 4, comprised periods of 12 – 24 hours, 24-48 hours, and 48–72 hours post-irradiation for image acquisition. Notably, comparing the structural events across all four post-irradiation stages from different experiments, we consistently observe similar dynamics of nuclear/nucleolar changes that become increasingly profound after 24-48 hours [SFig. 10-12]. Meanwhile, survival of cells over 72 hours was demonstrated in SMovie 1.

#### Stage 1: Post-irradiation changes, over the first 12 hours

Related changes were analyzed using a 2D movie time series at medium magnification. Individual LSM images reflecting different stages of post-radiation nuclear evolution were presented and analyzed either in fluorescent regime or merged PC and fluorescence mode (SMovie 2). The latest has been used for better discriminate between mononuclear and multinuclear cells. Key time points were identified and extracted for detailed analysis in galleries of individual/random sections, revealing cellular and nuclear shapes by treatment under a 30 Gy regime. During stage 1, the culture maintained its nuclear and nucleolar shape and structure without drastic changes over the 12-hour acquisition period, resulting in pictures that were largely similar to the control ones (SFig. 13, 14). Mitotic cells could be detected quite often.

**Figure 13.**
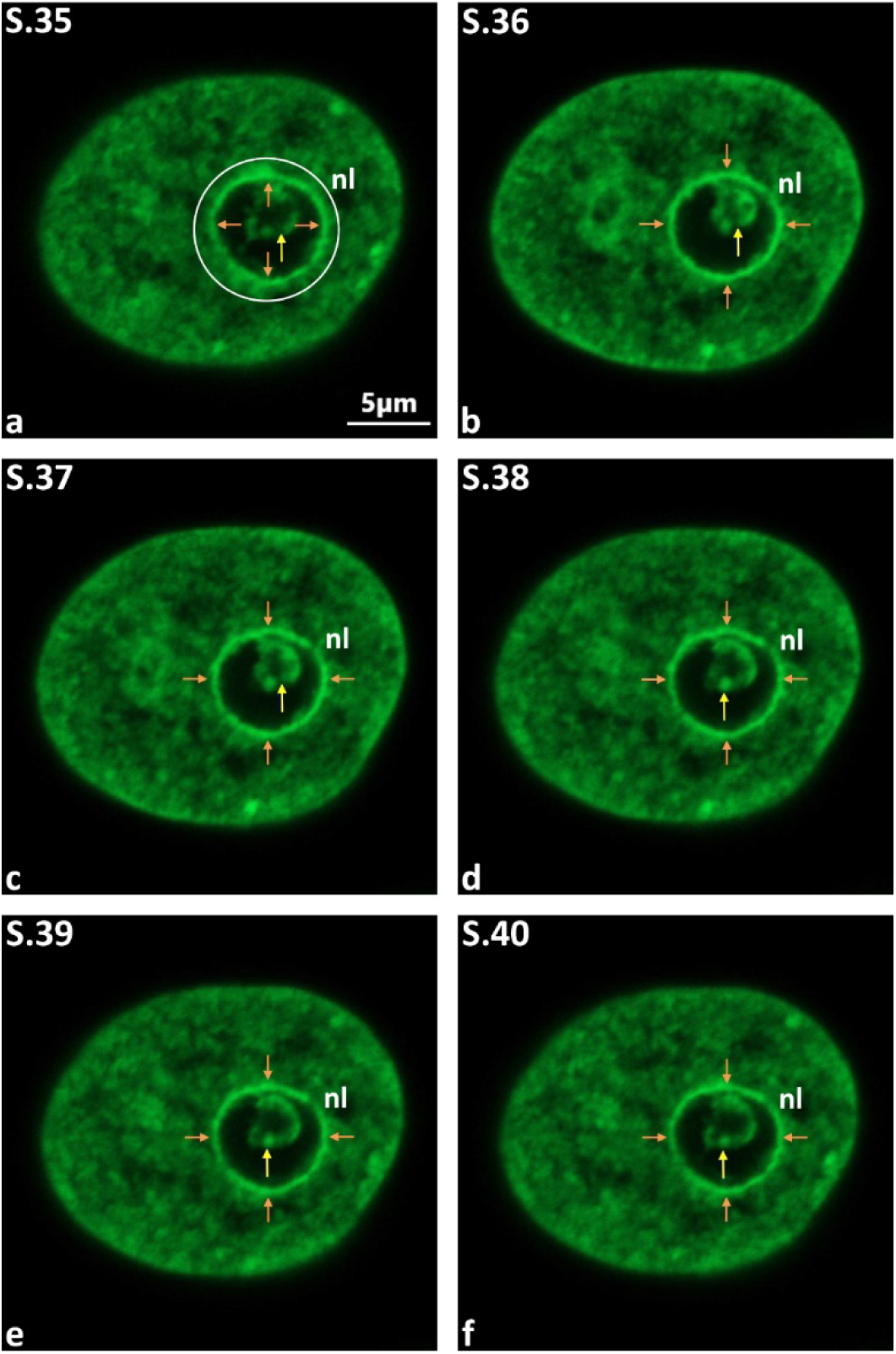
30 Gy γ-irradiation, 48 hours of post-irradiation image acquisition: mononuclear histone H2B-GFP HeLa cell. (a–f) Sequential Z-section images (S.35–S.40) reveal a densely packed ICC aggregate forming a ring-like structure (indicated by yellow arrows) within the nucleolar region (outlined in white). Additionally, a distinct PCC ring (marked by brown arrows) surrounds the nucleolus, indicating significant chromatin condensation. Abbreviations as in previous figures.

#### Stage 2: Post-irradiation changes over 12 - 24 hours

During the 12 - 24 hours (Fig 11, 12; SFig. 15-18; SMovie 3) of the post-irradiation period, massive nuclear/nucleolar changes were not detected in surviving cells. However, the minor cell population, containing nuclei with profoundly deformed contours as well as apoptotic and multinuclear cells, has occasionally been registered. The bulk of survived/surviving cells retain mononuclear appearance, while nuclei maintain roundish or ovoid shape with nearly smooth outlines. Over a 12-hour period, cells with slightly deformed outlines were sparsely observed; however, their quantity progressively increased to 24 hours. Slightly deformed nuclei exhibit wave-like periphery, so that deep invaginations were absent.

#### Stage 3: Post-irradiation changes over 48 hours

During the 24 – 48 hours (Fig. 13-18; SFig. 19- 27; SMovie 4) of post-irradiation period, the nuclear diameter progressively increased, ranging an average between 23.5 and 30.5 μm. However, the majority of cells remain mononuclear with nuclei maintaining ovoid shape and smooth outlines. Despite the prominent cellular destruction, rare mitotic cells could be observed (SFig. 22). However, after 48 hours, strongly deformed cells increase in number, while multinuclear and apoptotic cells appear in abundance (SFig. 19, 20). The deformed nuclei exhibit deep invaginations, ranging from one to multiple. Multinuclear cells can be regularly observed, while apoptotic cells (including early stages) can be easily recognized due to the dramatically enhanced nuclear brightness of increasingly compacted chromatin. The number of multinuclear and apoptotic cells significantly increases by the end of the acquisition time, i.e., after ∼40 hours of the post-irradiation period. In parallel, the nucleolar changes gradually increase, so that the intranucleolar fluorescence becomes significantly brighter and more prominent than in previous samples, due to the disorganization of the ICC network and its transformation into large clumps. Often, ICC clumps seemed to be reshaped, acquiring a ring- like appearance. Moreover, this experimental point is characterized by notable thickening of the PCC ring (Fig. 13, 14, 16; SFig. 21, 23, a1).

**Figure 14.**
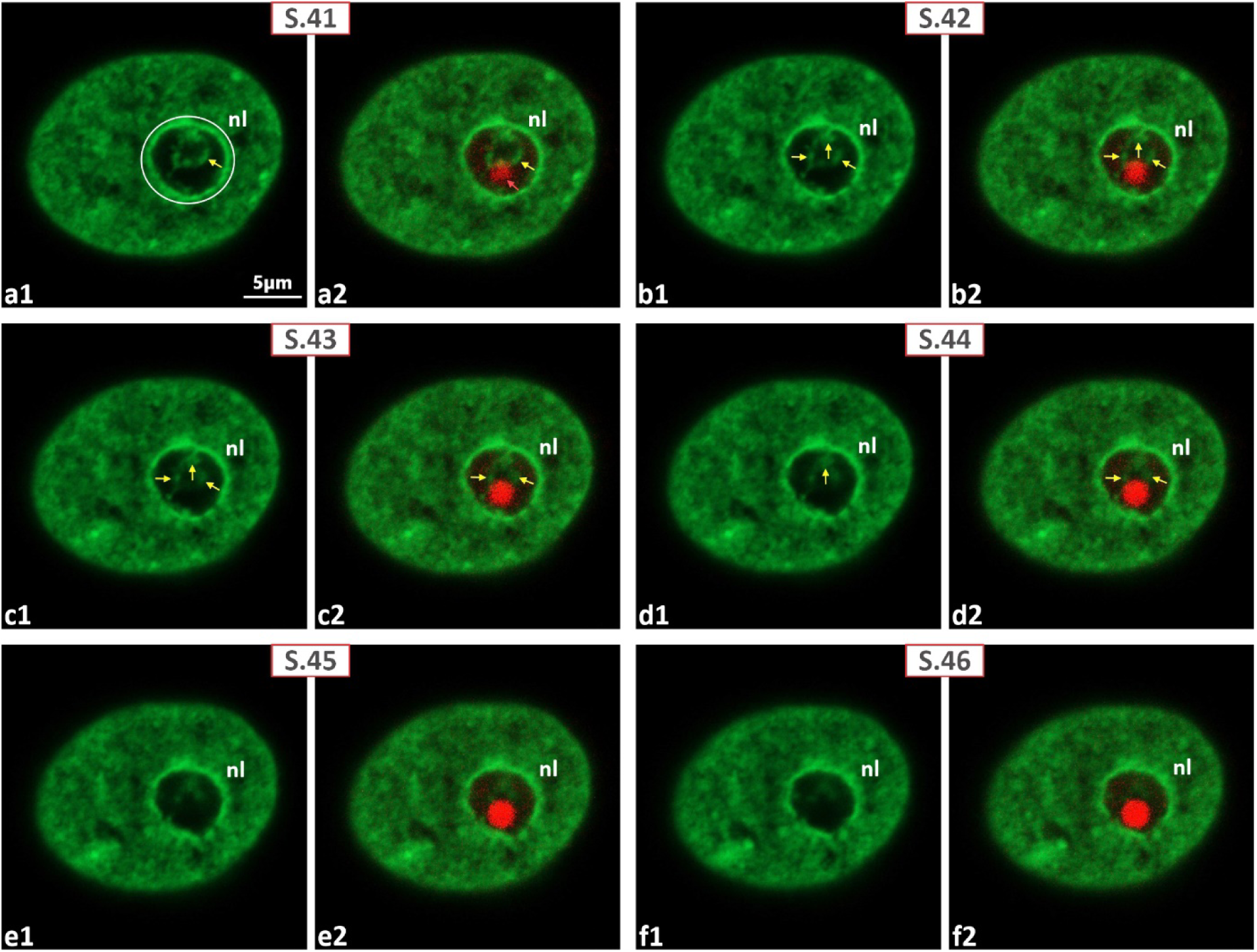
30 Gy γ-irradiation, 48 hours of post-irradiation image acquisition: same nucleus as Fig. 13. Histone H2B-GFP signal merged with anti-UBTF immunostaining. (a1–f1) Serial Z-sections (S41–S46) show a ring-shaped GFP-positive ICC clump. (a2–f2) The ICC ring forms a tight complex with a giant UBTF-positive spheroid; UBTF contacts the PCC are evident in c2– f2. Abbreviations as in previous figures.

**Figure 15.**
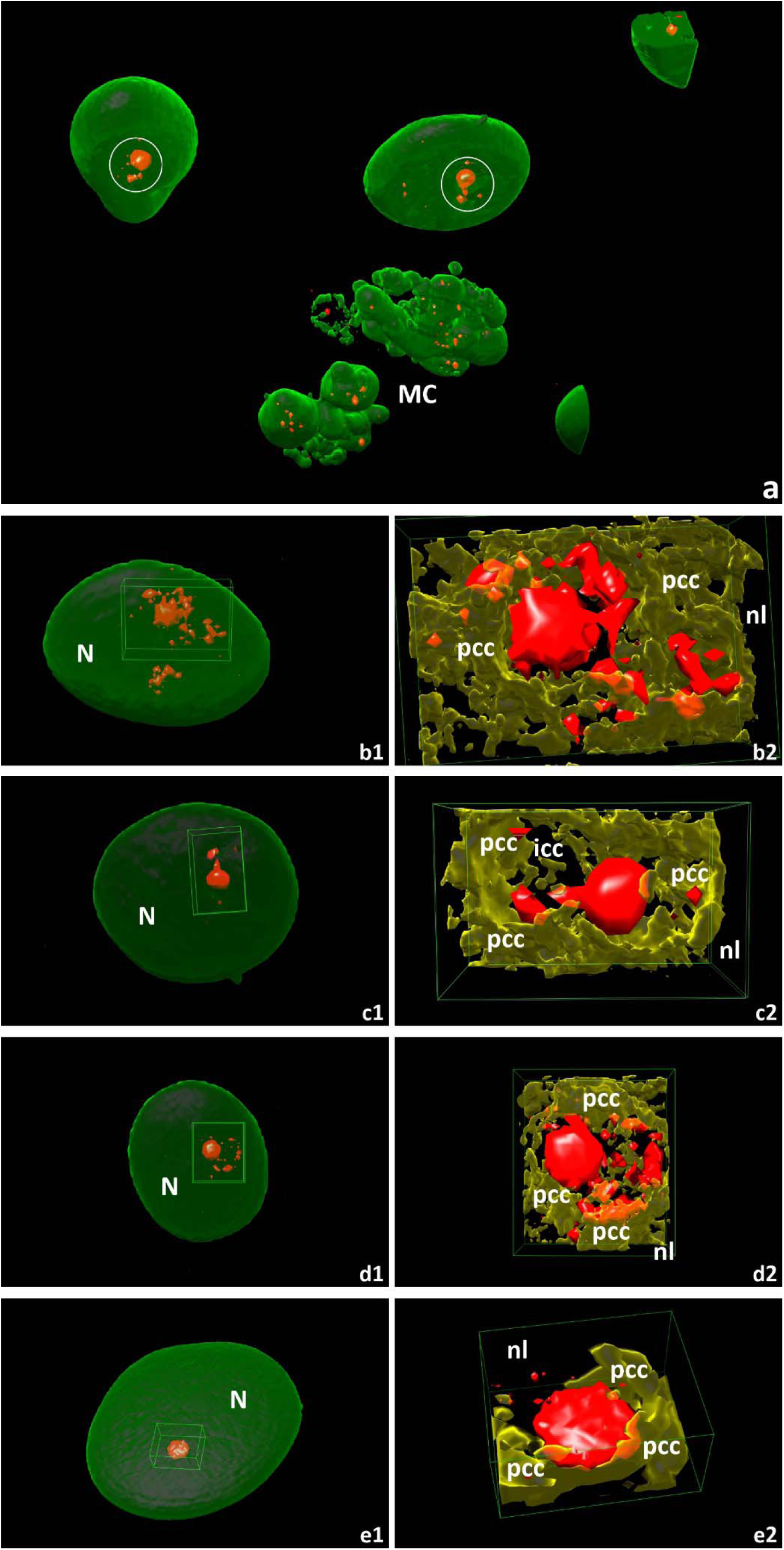
Mononuclear and multinuclear histone H2B-GFP HeLa cells, 48 hours of post-irradiation acquisition after 30 Gy γ-irradiation; anti-UBTF immunostaining with 3D nuclear/nucleolar models. (a) Low-magnification view reveals drastic, asymmetric enlargement of UBTF-positive bodies (red); multinuclear cells show UBTF signal dispersed among nuclear fragments. (b1–e1) 3D overviews of nuclei (nucleolar territories outlined in white) containing one giant and several minor UBTF-positive bodies. (b2–e2) Extracted models highlight these UBTF-positive bodies seamlessly embedded within the NAC shell (yellow).

**Figure 16.**
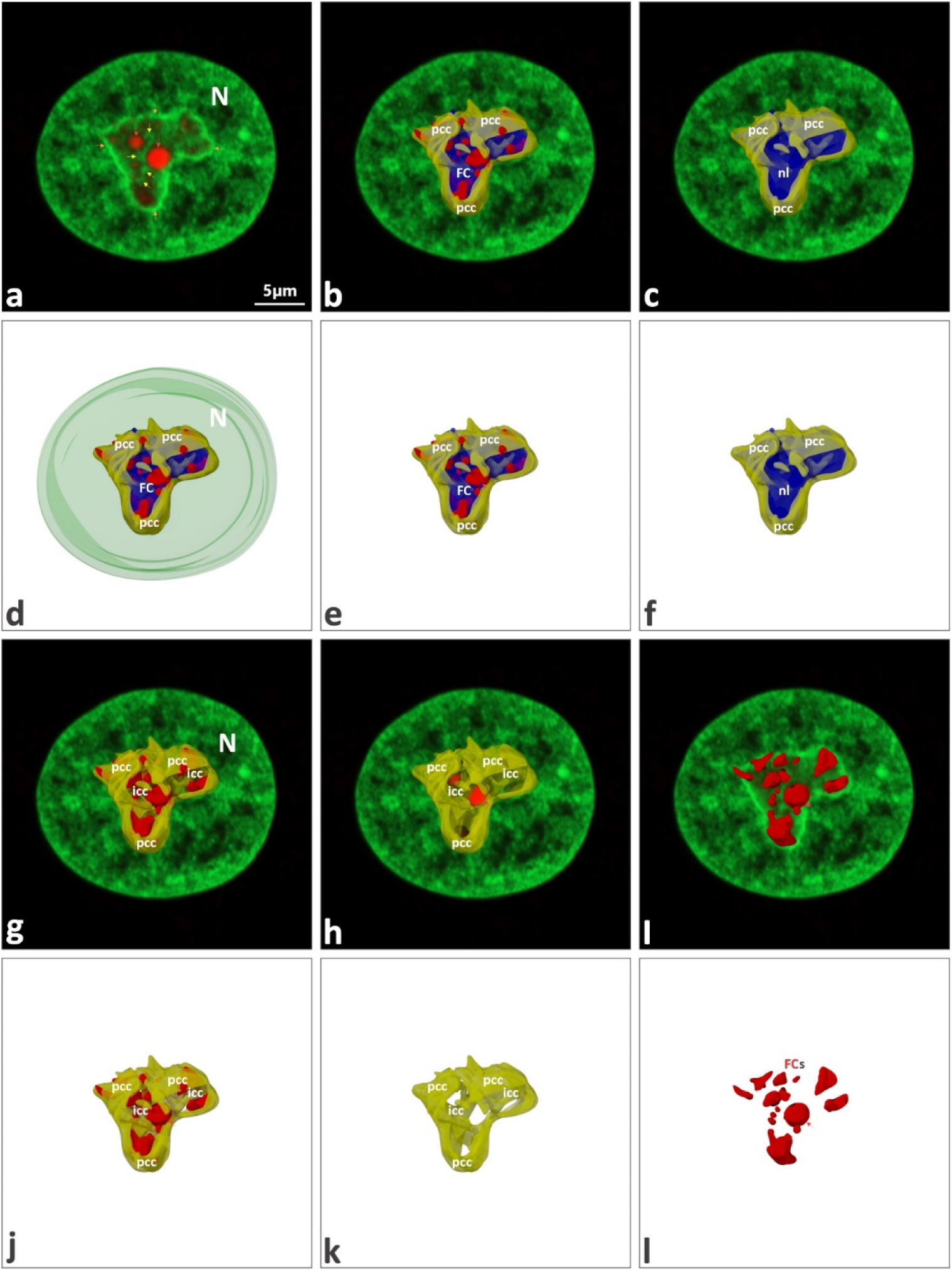
Mononuclear histone H2B-GFP HeLa cell, 48 hours of image acquisition after 30 Gy γ-irradiation; anti-UBTF immunostaining with 3D nuclear/nucleolar renderings (AutoCAD). (a) Single 2D section shows a giant UBTF-positive body (red) abutting the PCC ring. (b) Same plane merged with 3D models of the nucleolus (blue), UBTF-positive bodies, and NAC shell (yellow). (c) Same plane with only nucleolus and PCC. (d–f) Sequential 3D views: all components (d), nucleolus + UBTF + PCC (e), nucleolus + PCC (f). (g, h) Sectioned nucleolar models at two depths highlight tight UBTF–PCC contacts. (i) UBTF distribution alone on the 2D section reveals three giant UBTF bodies firmly anchored to the PCC. (j–l) 3D renderings: NAC + UBTF (j), NAC alone (k), and UBTF-positive bodies alone (l, red arrows). These asymmetrically enlarged UBTF bodies—likely FCs—remain embedded within the NAC. FC - fibrillar center; other abbreviations as in previous figures.

**Figure 17.**
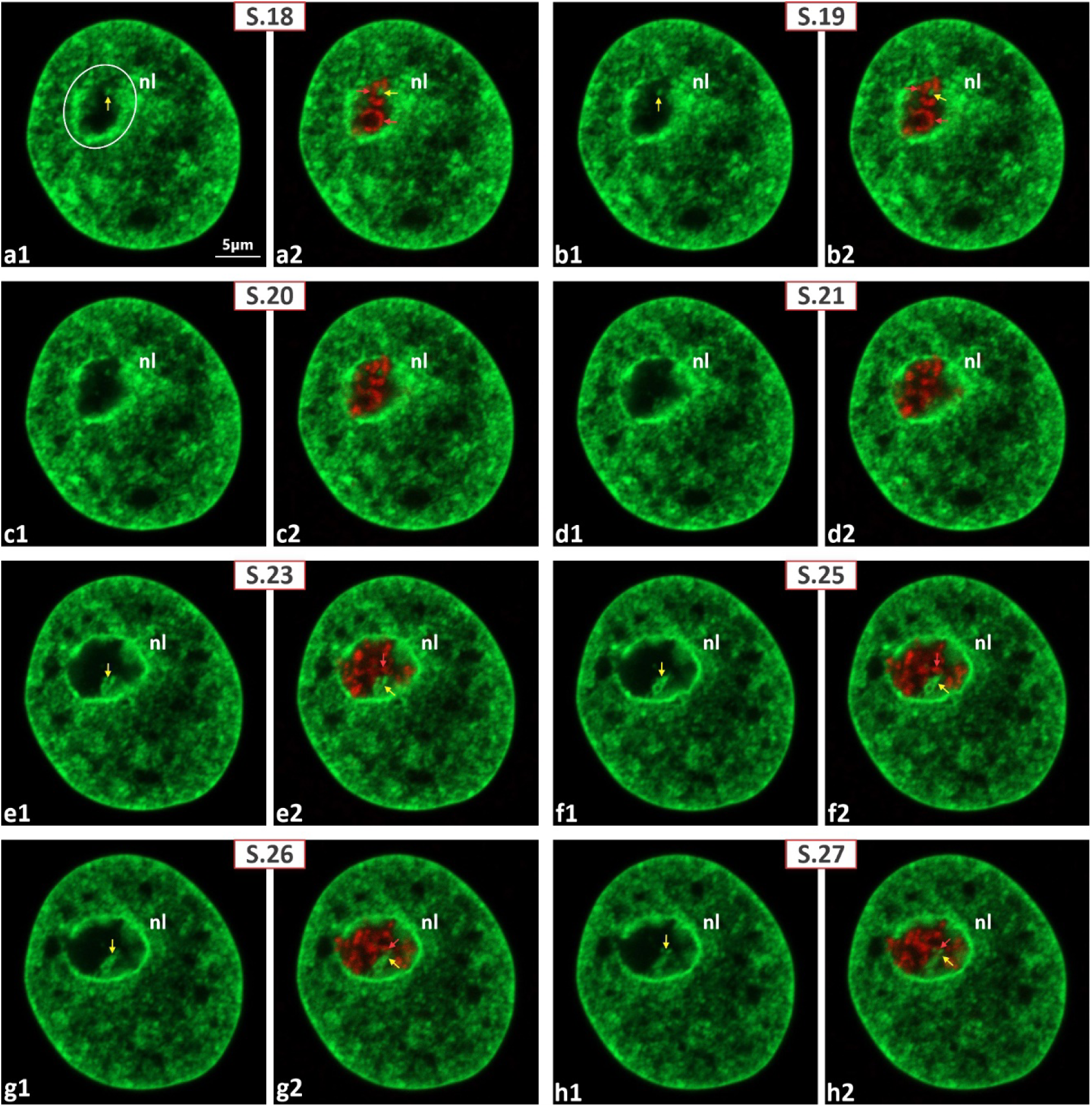
Mononuclear histone H2B-GFP HeLa cell, 48 hours of post-irradiation image acquisition after 30 Gy γ-irradiation; anti-fibrillarin immunostaining. (a1–h2) Gallery of Z-sections (S.18–S.21, S.23, S.25–S.27). The nucleolar territory is outlined in white (17, a). Fibrillarin-positive structures (red) largely preserve their cord-like architecture even 48 hours of post-irradiation image acquisition. Large GFP-positive ICC clumps (yellow arrows) remain tightly coupled to these cords and maintain contact with the prominent PCC ring. Abbreviations as in previous figures.

**Figure 18.**
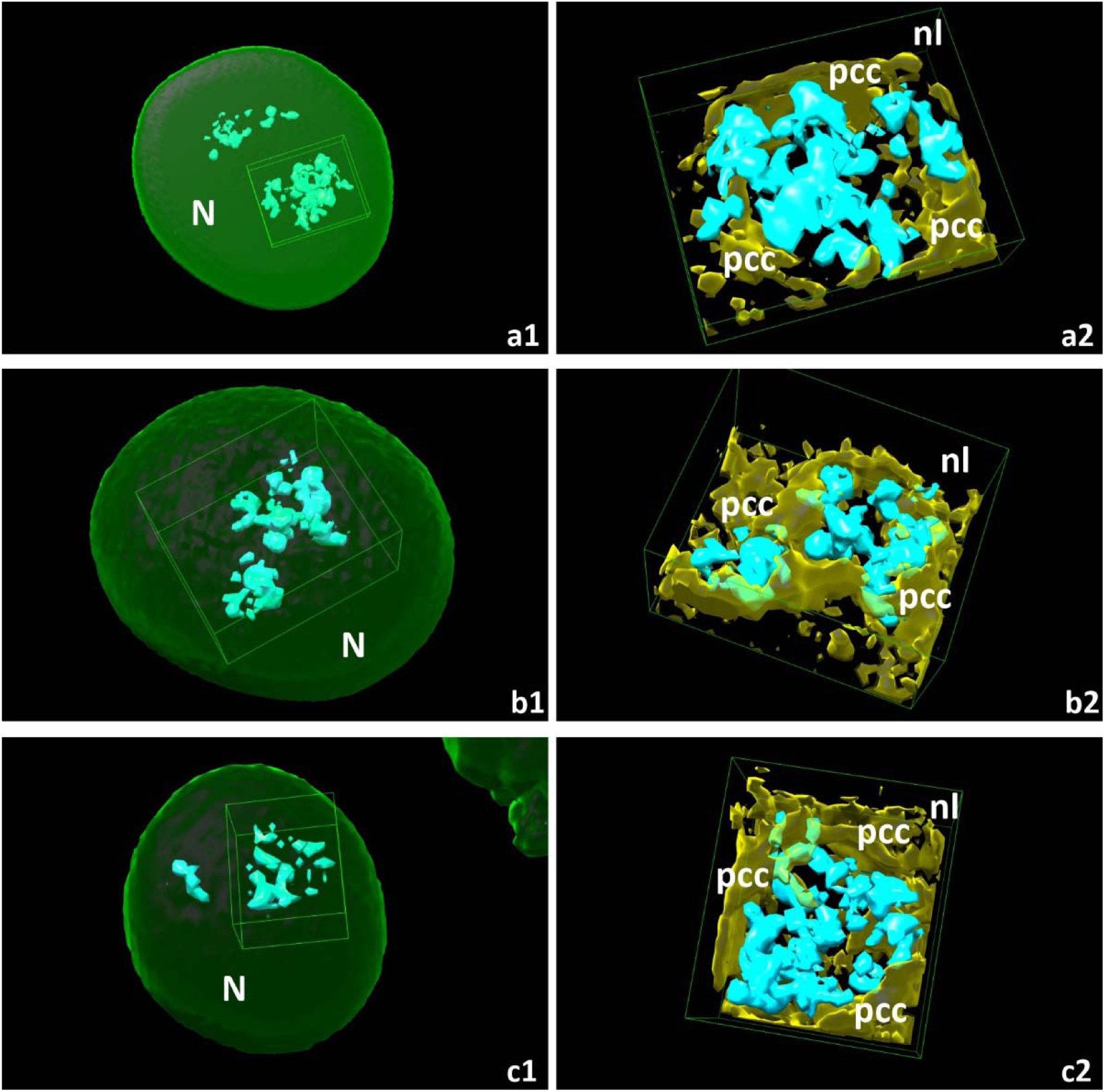
Mononuclear cells, 48 hours of post-irradiation image acquisition after 30 Gy γ- irradiation; anti-fibrillarin immunostaining with 3D nuclear models (UCSF Chimera). (a1–c1) Intranucleolar fibrillarin-positive structures (cyan) retain cord-like organization; nucleolar territories are marked by white rectangles. (a2–c2) Corresponding 3D renderings emphasize fibrillarin-containing cords seamlessly embedded within the NAC shell (yellow). Abbreviations as in previous figures.

**Figure 19.**
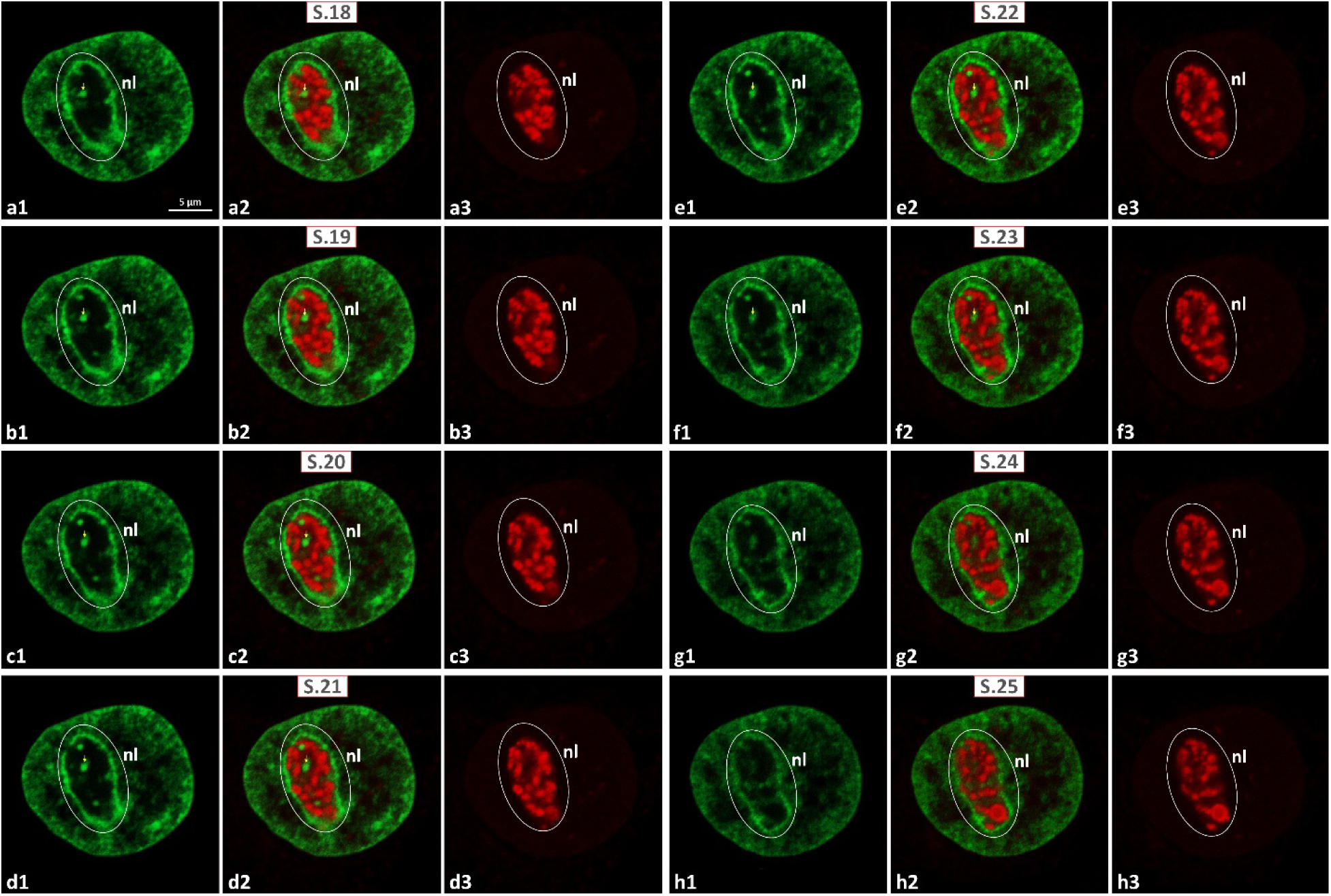
Mononuclear cell, 72 hours of post-irradiation image acquisition after 30 Gy γ- irradiation; anti-UBTF immunostaining. (a1–h3) Consecutive virtual Z-sections (S.18–S.25) with the nucleolar territory outlined in white; GFP channel only (a1–h1): a prominent PCC ring and large ICC clumps are evident (yellow arrows). Merged GFP + UBTF (a2–h2): chain-like arrays of UBTF-positive bodies (red) make frequent contacts with the large ICC clump (yellow arrows, S.18–S.23). Anti-UBTF fluorescence only (a3–h3): numerous spherical UBTF bodies form pronounced chains, indicating that even 72 h post-irradiation surviving cells maintain extensive UBTF organization. Abbreviations as in previous figures.

**Figure 20.**
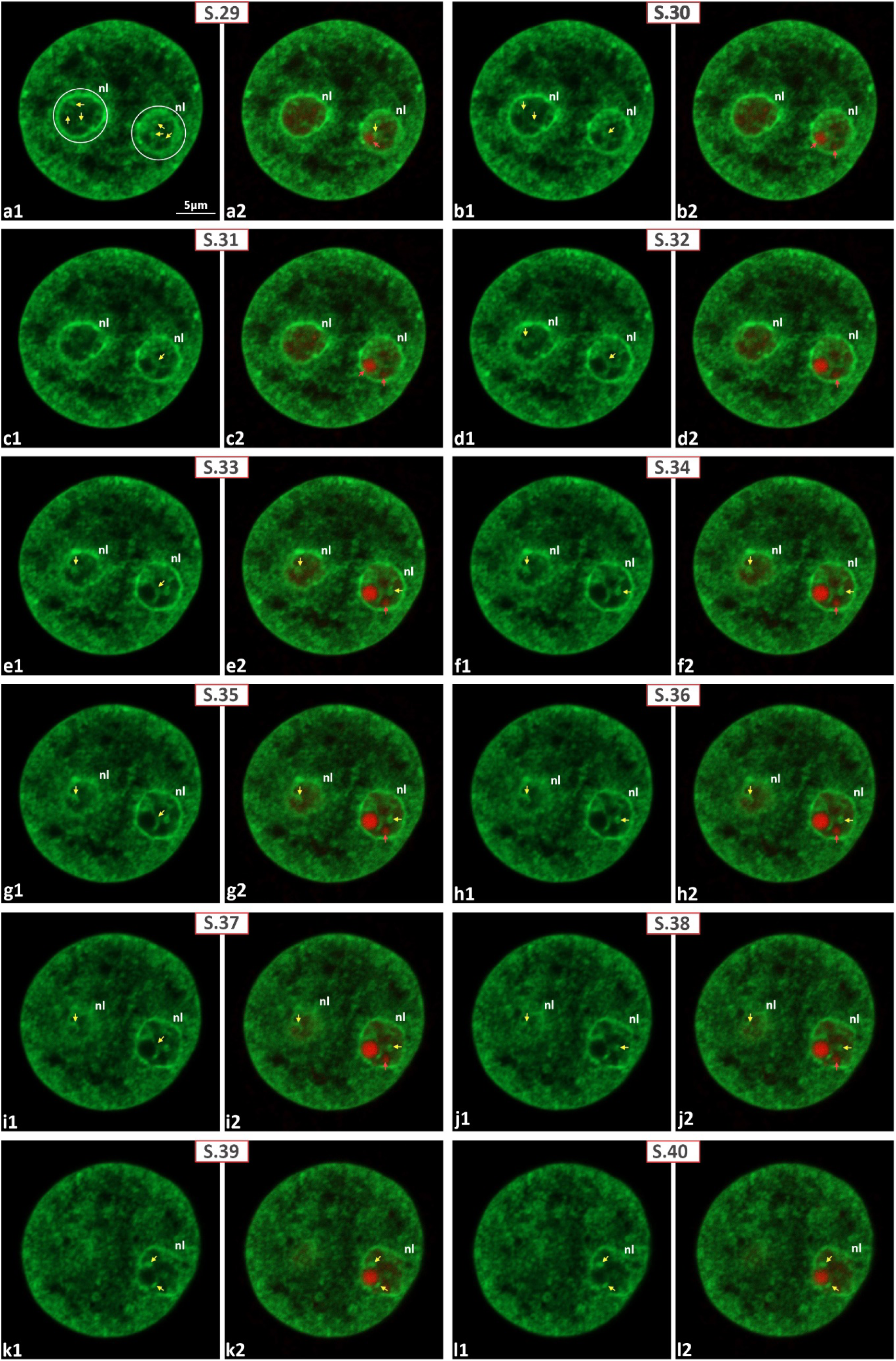
30 Gy γ-irradiation, 72 hours of post-irradiation image acquisition: the gallery of anti-UBTF immunostained nucleus of mononuclear cell images. Nucleolar territory outlined by white circle. (a1–l2) Serial Z-sections (S.29–S.40); GFP channel only (a1–l1): The NAC architecture remains united, with a clear PCC ring and well-defined ICC network; Merged GFP + UBTF (a2– l2): Dramatically enlarged UBTF-positive spheroid (red) is tightly embedded within the NAC and consistently about the PCC ring throughout the stack. Abbreviations as in previous figures.

**Figure 21.**
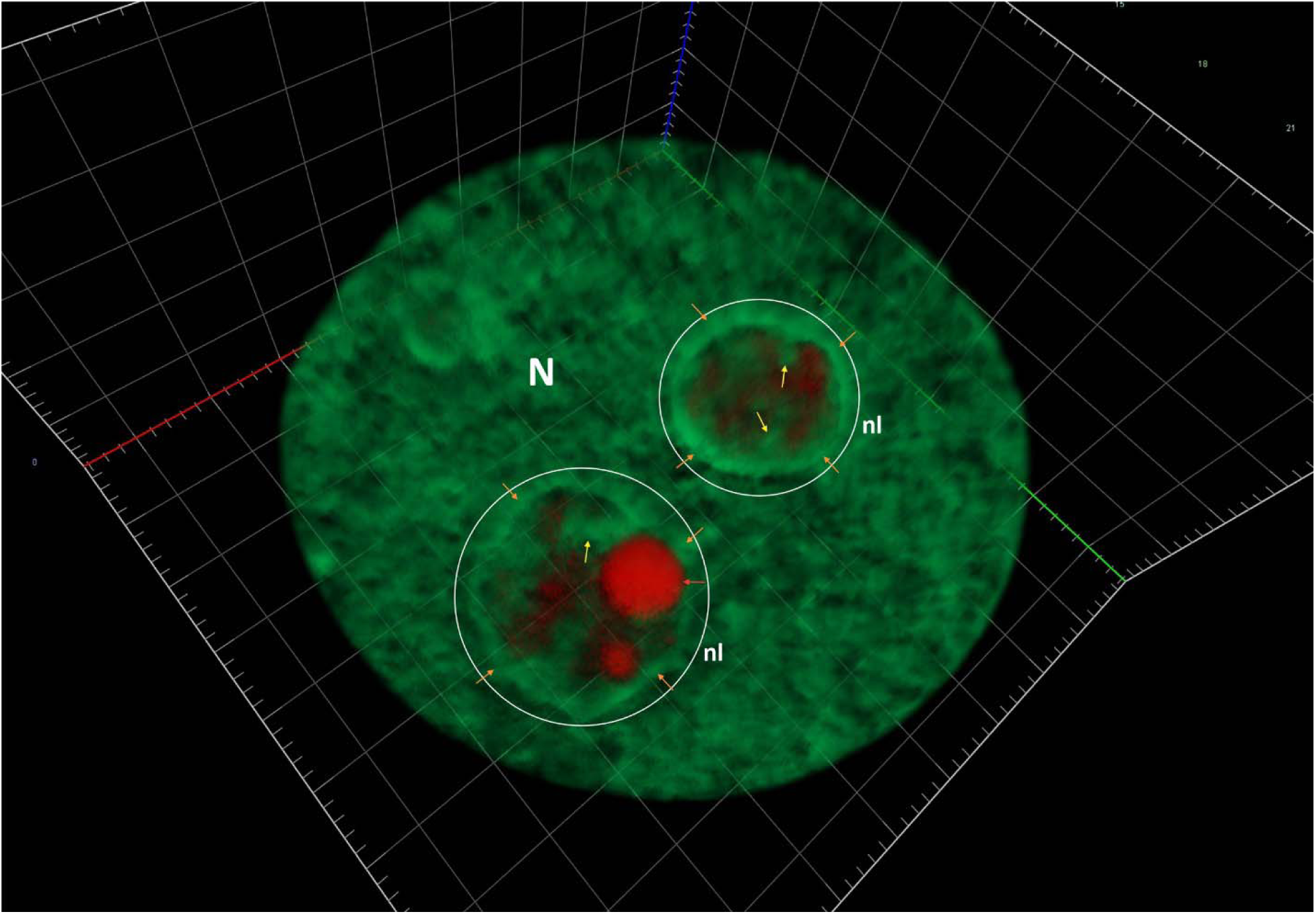
Mononuclear cell, 72 hours of post-irradiation image acquisition after 30 Gy γ- irradiation; 3D nuclear model (ZEN3.0) following anti-UBTF immunostaining. Equatorial nuclear and nucleolar sections reveal structural interplay between large ICC inclusions (yellow arrows) and giant UBTF-positive bodies (red). Three nucleoli of varying size are present, with the two largest outlined in white. All nucleoli are surrounded by a thick PCC ring (brown arrows). In addition to two enlarged UBTF structures, smaller faint UBTF-positive foci are visible. ICC cords originate as extensions of the PCC shell into the nucleolar territory. UBTF bodies—likely fibrillary centers—are clearly integrated into the NAC complex. Abbreviations as in previous figures.

**Figure 22.**
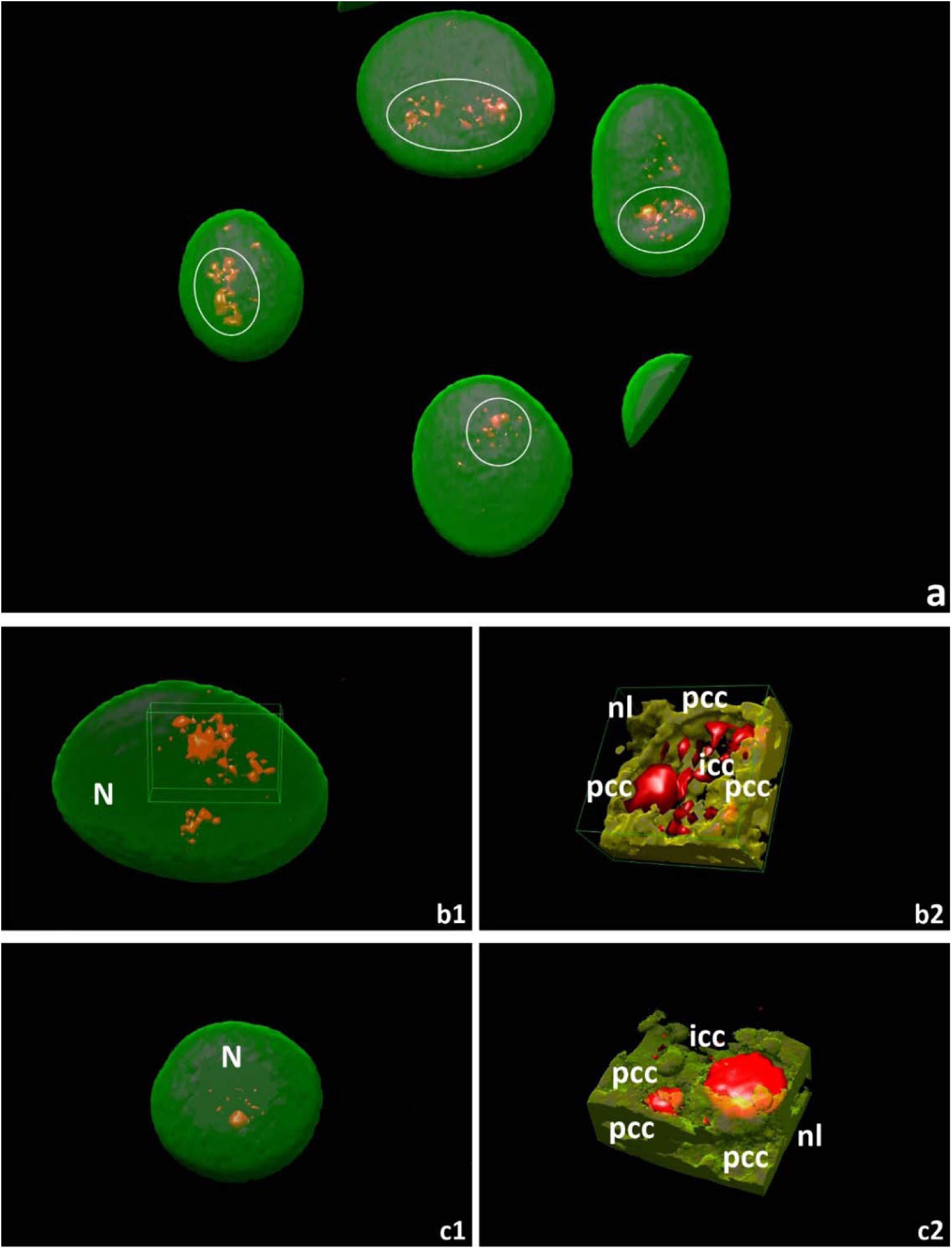
Mononuclear cells, 72 hours of post-irradiation image acquisition after 30 Gy γ- irradiation; anti-UBTF immunostaining with 3D nuclear/nucleolar models (UCSF Chimera). Nucleolar territories are outlined by white circles and rectangles. (a) Nuclei display nucleoli with varying numbers and sizes of UBTF-positive bodies (red). One to three UBTF structures show marked enlargement visible even at low magnification, while smaller UBTF-positive entities maintain chain-like organization. (b1–c1) 3-D models of nuclei highlight nucleolar territories (rectangles in 22, b1) containing one giant and several smaller UBTF bodies; some smaller entities preserve a chain-like spatial arrangement. (b2–c2) Extracted 3-D views demonstrate clear integration of UBTF-positive bodies within the NAC shell (yellow).

**Figure 23.**
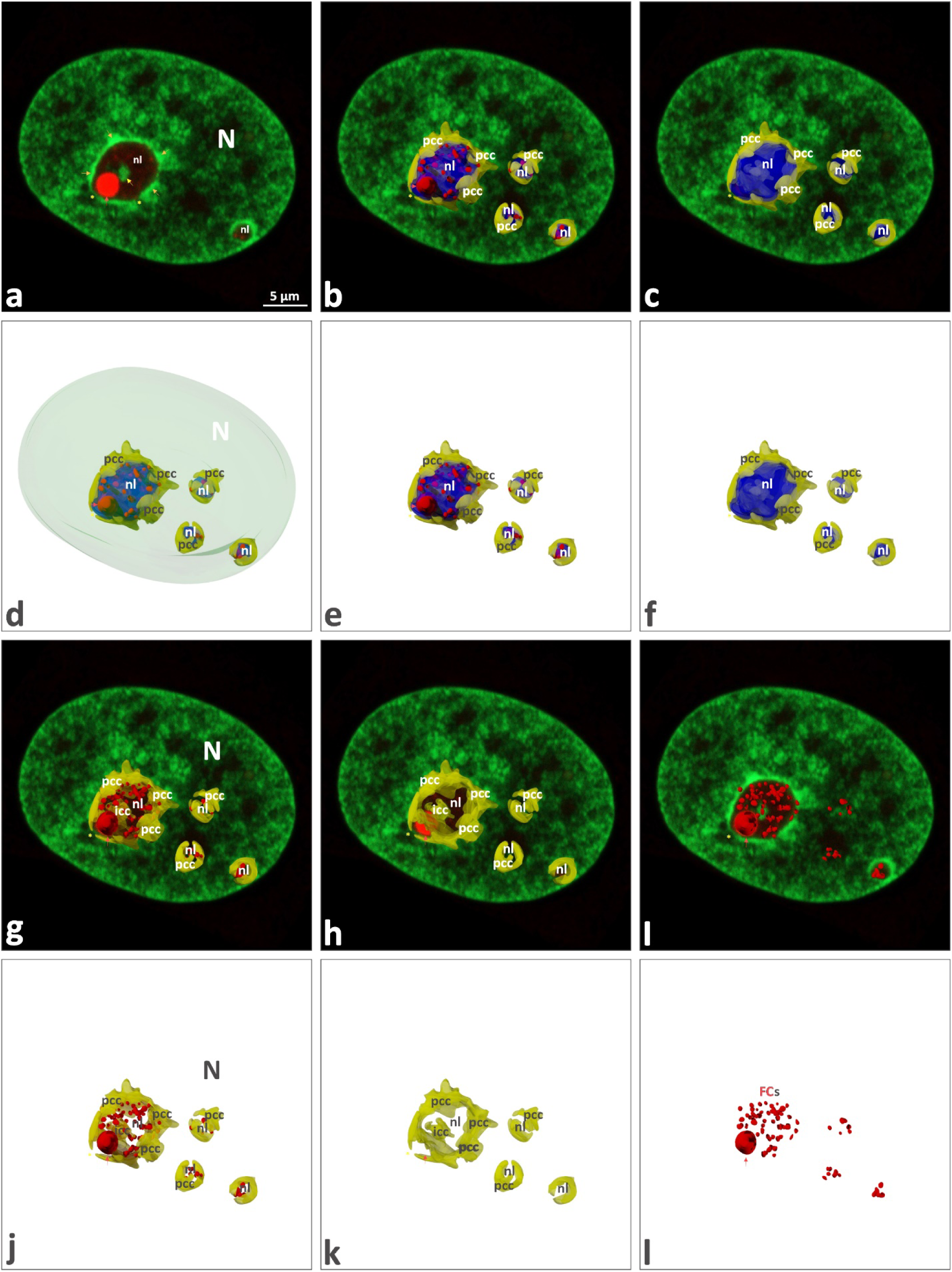
Mononuclear cell, 72 hours of post-irradiation image acquisition after 30 Gy γ- irradiation; anti-UBTF immunostaining with 3D nuclear/nucleolar models (AutoCAD). (a) Single 2-D section shows a giant UBTF-positive spherical structure (red) closely associated with a thickened PCC ring and a prominent ICC clump in contact. (b) Same section merged with 3D models of nucleoli (blue), UBTF bodies, and NAC shell (yellow). (c) Same plane merged with nucleolus and NAC only. (d–f) Sequential removal of model layers illustrates nucleolus position within the nucleus and its structural relations to UBTF-positive bodies and NAC: all components (d), nucleolus + UBTF + NAC (e), nucleolus + NAC (f). (g, h) Sectioned nucleoli reveal tight contacts between PCC shell and UBTF bodies, including the giant UBTF-positive spheroid. (i) 2D section with UBTF-positive bodies alone shows one giant and numerous smaller (likely pseudo-FCs) UBTF-positive structures bridged by a large ICC clump. The giant UBTF-positive body is firmly attached to the PCC ring. (j–l) 3D renderings depict NAC + UBTF (j), NAC alone (k), and UBTF-positive bodies alone (l). All UBTF bodies integrate into the NAC; the large UBTF body likely represents an enlarged fibrillary center, while smaller ones correspond to pseudo-FCs. Abbreviations as in previous figures.

#### Stage 4: Post-irradiation changes over 48 - 72 hours

The most profound effects of 30 Gy γ- irradiation, leading to apparent nuclear and nucleolar structural modifications and drastic changes in global cellular organization, were observed between 48 and 72 hours (Fig. 19-26; SFig. 28-36; SMovie 5). Consequently, a bulk of the cells underwent a transformation from mono- to multinuclear forms. For example, massive apoptosis was detected starting from 56-60 hours into the post-irradiation period, whereas the number of wholly destructed/dead cells dramatically increased to the end of time-lapse imaging, i.e., to 70-72 hours. It should be specially noted that, mechanism of MC-induced multinucleation is passing through: (i) progressive nuclear invagination leading to a lobulated shape; (ii) asymmetric nuclear fragmentation into unequal-sized micronuclei and (iii) endomitotic division of nuclear fragments, followed by apoptotic nuclear degradation (SMovie 6, a). Likely, the apoptotic process starts right after the final appearance of a multinuclear cells (SFig. 30, 32).

Analyzing nucleolar structural changes unraveled by imaging using PC at medium magnification regimes, we registered the emergence of large light zones inside the nucleolus (SFig. 33). At high magnification and in the fluorescent regime, clearly corresponding nucleolar modifications are concomitant with the appearance of extensive ICC/FC unity, so that ICC sometimes acquire a ring-shaped appearance. Concerning γ-irradiation-induced dynamics of GFP positive nucleolar chromatin, we detected that related changes largely resembled NAC rearrangements observed with chemical inhibitors. Importantly, during chemical inhibition, the ICC network was captured in contracted state, forming prominent clumps instead of an intensively branched network (Fig. 20, 21, 23, 24; SFig. 34, 36) [61]. Surprisingly, even after being γ-irradiation damaged, ICC retains intra-nucleolar mobility, as revealed by 2D movies, which correspond to 48-72 (SMovie 6, a) and 56 - 72 (SMovie 6, b) points of the post-irradiation acquisition period. Movie 6, a demonstrates the reshuffling of a prominent ICC clump from the nucleolar interior toward the PCC shell. Moreover, the peripheral migration of ICC was accomplished by the fusion of these two NAC constituents.

**Figure 24.**
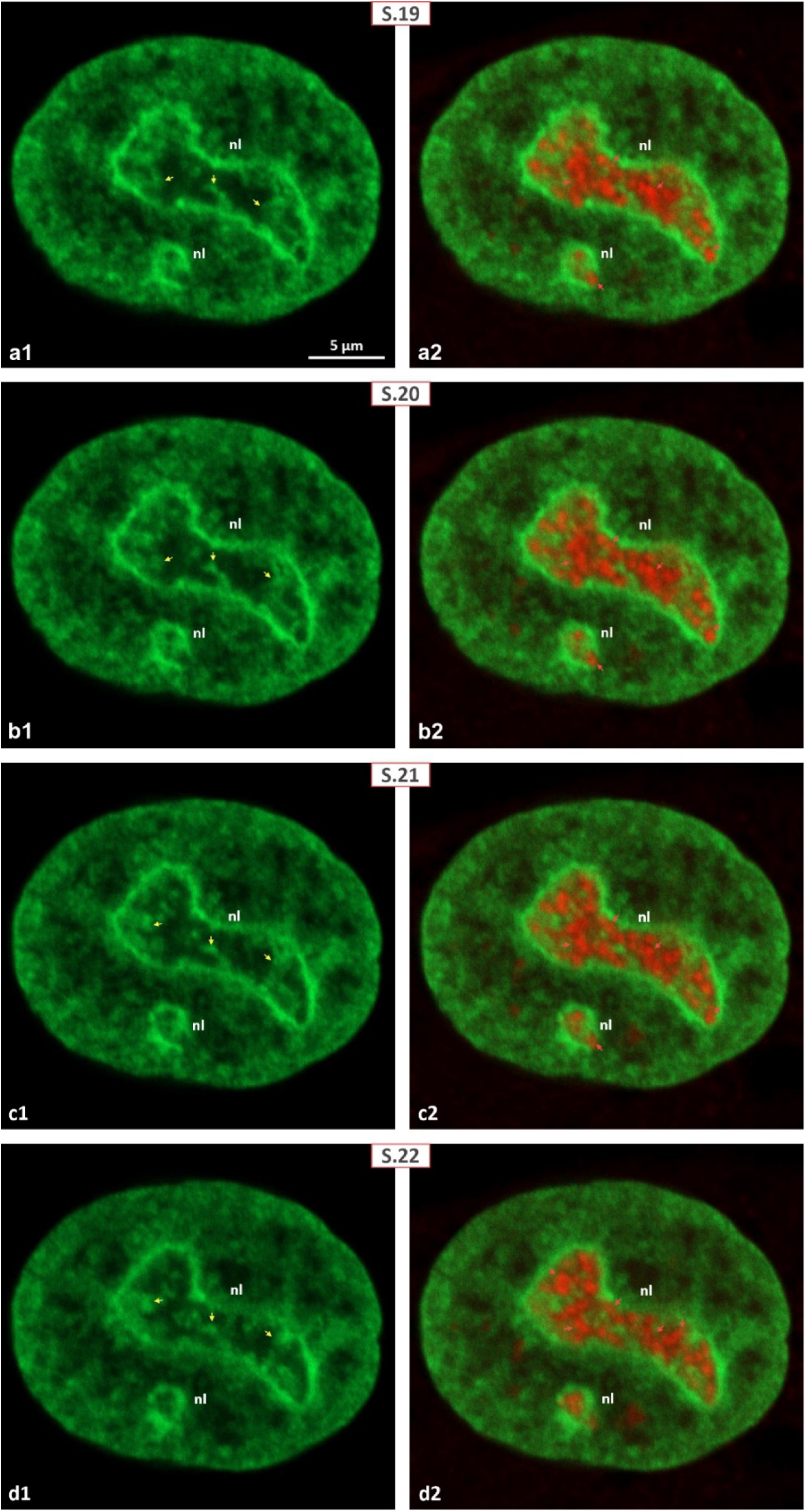
Mononuclear cell, 72 hours of post-irradiation image acquisition after 30 Gy γ- irradiation; anti-fibrillarin immunostaining. Gallery of serial Z-sections (S.19–S.22) shows fibrillarin-positive structures (red) maintaining their cord-like organization. (a1–d1) GFP channel only highlights the NAC system, including a thickened PCC shell and prominent ICC network extending from the PCC. (a2–d2) Merged images show fibrillarin-positive cords tightly integrated with the NAC, consistently contacting the PCC shell across sections. Abbreviations as in previous figures.

### 3.3. Visualizing Changes in FC/DFC Assembly 3D Structure and Relationship with NAC System: γ-Irradiation Doesn’t Provoke Nucleolar Capping

It has been well established that treatment of HeLa cells with AMD and many other chemical inhibitors leads to the classical segregation of nucleoli into distinct nucleolar caps. However, observed nucleolar changes develop in a profoundly different way. Post-irradiation immunolabeling clearly showed that upon γ-irradiation induced nucleolar DNA damage, only UBTF-positive structures became larger while their number decreased gradually (Fig. 11, 13-16, 19-23, 26; SFig. 15, 16, 19-23, 28-32, 34). The behavior of fibrillarin-positive DFC is entirely different, retaining its cord-like organization during the entire 72-hour acquisition period (Fig. 12, 17, 18, 24-26; SFig. 17, 18, 24-27, 35, 36). Only in multinuclear cells massively appearing during 48 – 72 hours fibrillarin-positive cords transform into spherical entities being shared among micronuclei. Below, we have arranged post-irradiation nuclear/nucleolar and NAC reorganization, according to the four stages mentioned.

*<u>Post-irradiation stage 1.</u>* Visually, during the initial 12 hours of γ-irradiation, the nuclei remained unchanged in size, appearing regularly shaped and ovoid in the 3D view, a characteristic typical of control cells (SFig. 13, 14). In addition, such treatment doesn’t induce visible morphological changes in nucleoli. They remain large (reaching up to 8 µm) and irregular, continuously revealing an intensely branching ICC network. In anti-UBTF immunolabeled samples irradiated for 0-12 h, (SFig. 13) its distribution was similar to the control 2D/3D pattern. Correspondingly, UBTF was exclusively visualized within spheres of 0.3–0.5 µm in diameter, organized in a chain-like manner. Anti-fibrillarin labeling (SFig.14) of irradiated cells submitted to 12 hours of post-irradiation image acquisition also yielded a picture close to the protein distribution registered in control samples. Visualization through 2D images and as 3D models, fibrillarin was permanently compartmentalized in “worm-like” cords, largely resembling normal DFC organization (Fig 3, 7-10).

By simultaneously visualizing 2D/3D histone H2B and UBTF in irradiated cells imaged within the first 12-hour period, we did not observe any significant signs of NAC network reorganization. As previously mentioned, profound structural alterations in nuclear shape and chromatin structure weren’t detected at this time, so that registerable minor structural changes were restricted to the nucleolar territory. Although at first glance the sizes and contours of nucleoli looked almost like in control, signs of coarsening/contraction of the NAC network became recognizable, including increased GFP fluorescence.

*<u>Post-irradiation stage 2.</u>* Over 12–24 hours, we observed a gradual increase in the coarsening of ICC inclusions and thickening of the PCC shell. These changes became significantly prominent closer to the 24-hour post-irradiation image acquisition period. In parallel, the sizes of UBTF- positive structures were also gradually increased, so that by the end of stage 2, nucleoli in some irradiated cells revealed much more profound link between PCC, ICC, and prominently enlarged UBTF-positive FCs (Fig. 11, SFig. 15, 16). Here, we found cells with peculiarly appearing nucleoli, where one or two asymmetrically enlarged UBTF-containing spheroids were disposed of the nucleolar periphery. These spheroids could lie close to each other, being deeply immersed in the subjacent PCC shell. Large spherical UBTF- and fibrillarin-positive SOI were permanently in contact with ICC or PCC (Fig. 11, 12). Likely, exposure of cells to γ-irradiation didn’t disorganize DFC as revealed by 2D/3D visualization of anti-fibrillarin labeling. Therefore, presented on Fig.12, SFig. 17, 18 anti-fibrillarin labeling pattern didn’t change much, retaining folded cord-like appearance.

By comparing images of histone H2B-GFP and UBTF as well as UBTF and fibrillarin fluorescence, in nucleoli containing large UBTF-positive spheroids, we also detected a prominent ICC clumps that looked like bridging the gap between the spheroids (Fig. 11). In turn,

H2B-GFP clumps link to the PCC shell, thus joining the UBTF-positive structures or FCs and NAC into a unified system. In all nucleoli analyzed after 24 hours of post-radiation acquisition, UBTF-positive FCs were located inside the nucleolar territory. As a consequence of cell exposure to γ-irradiation, enlarged UBTF-positive NCs were shifted to the margin between the PCC shell. Judging by topography of fibrillarin 3D redistribution, likely nucleoli still contain DFC, organized into “worm-like” cords. In all cells analyzed, we couldn’t observe the formation of nucleolar caps.

*<u>Post-irradiation stages 3 and 4.</u>* Here, beside the bulk of cells looking relatively “healthy” or just slightly altered, cells abundantly bearing severely deformed or cleaved nuclei, multinuclear and apoptotic cells became the most prominently expressed. Signs of cellular destruction and death, also were abundantly pronounced (Fig. 15, 26 SFig. 19, 20, 25, 27, 30, 32). Interestingly, even after a 48-hour post-irradiation period, we observed occasional dividing cells, with well pronounced UBTF-positive NORs (SFig. 22, 24). By analyzing 2D images and, particularly, by rotating 3D models reconstructed from the nuclei of severely damaged cells, we found that many cells contained nuclei that were deformed or even lobbed due to deep invagination. Micronuclei reveal profound UBTF-positive structures, dispersed inside lobes and/or separated micronuclei.

**Figure 25.**
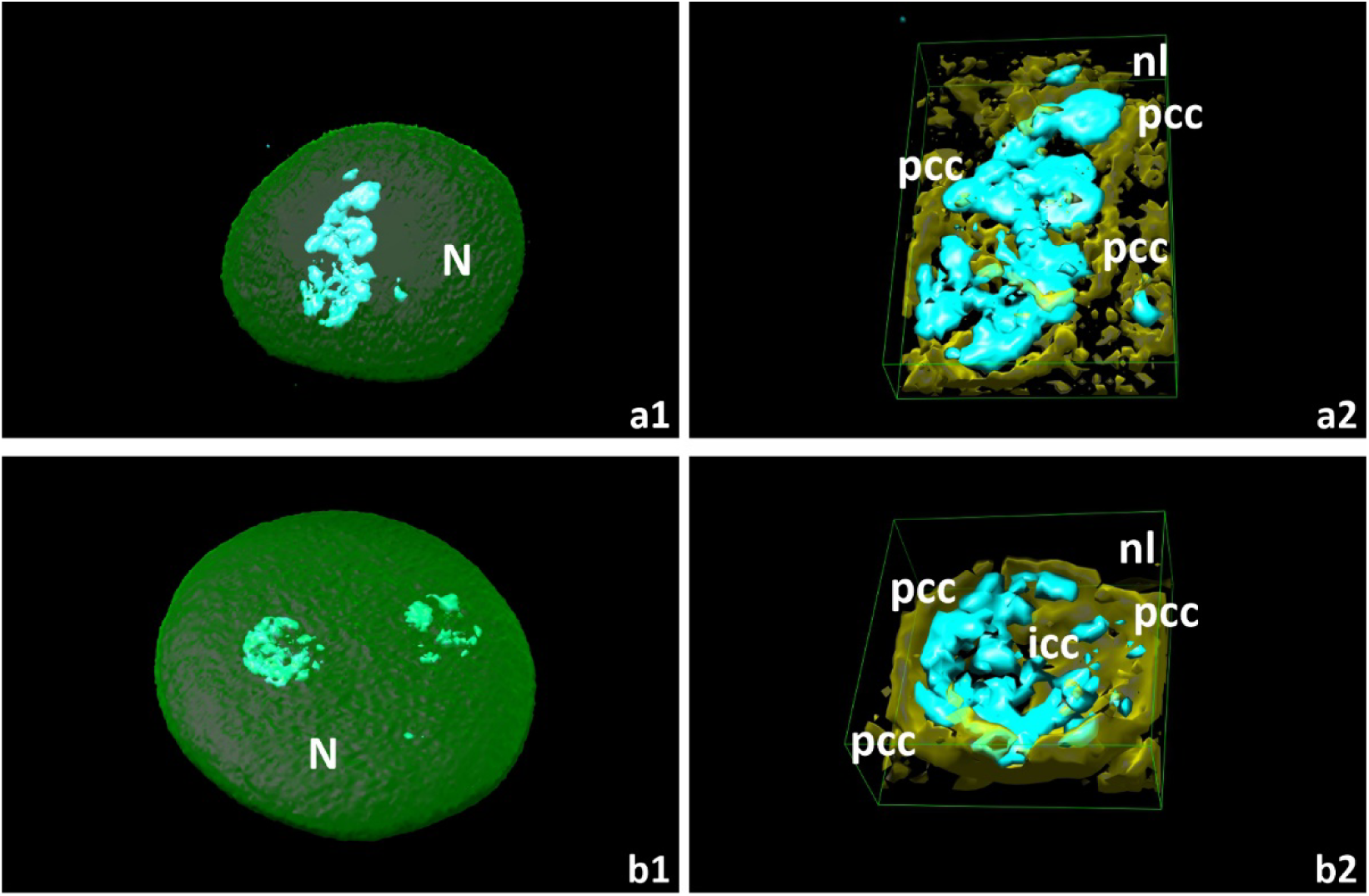
Surviving mononuclear cells, 72 hours of post-irradiation image acquisition after 30 Gy γ-irradiation; anti-fibrillarin labeling with 3D nuclear models (UCSF Chimera). (a1–b1) Fibrillarin-positive structures (cyan) exhibit distinct cord-like intranucleolar organization. (a2– b2) Extracted 3D nucleolar models demonstrate tight integration of fibrillarin-positive cords within the NAC shell (yellow). Abbreviations as in previous figures.

**Figure 26.**
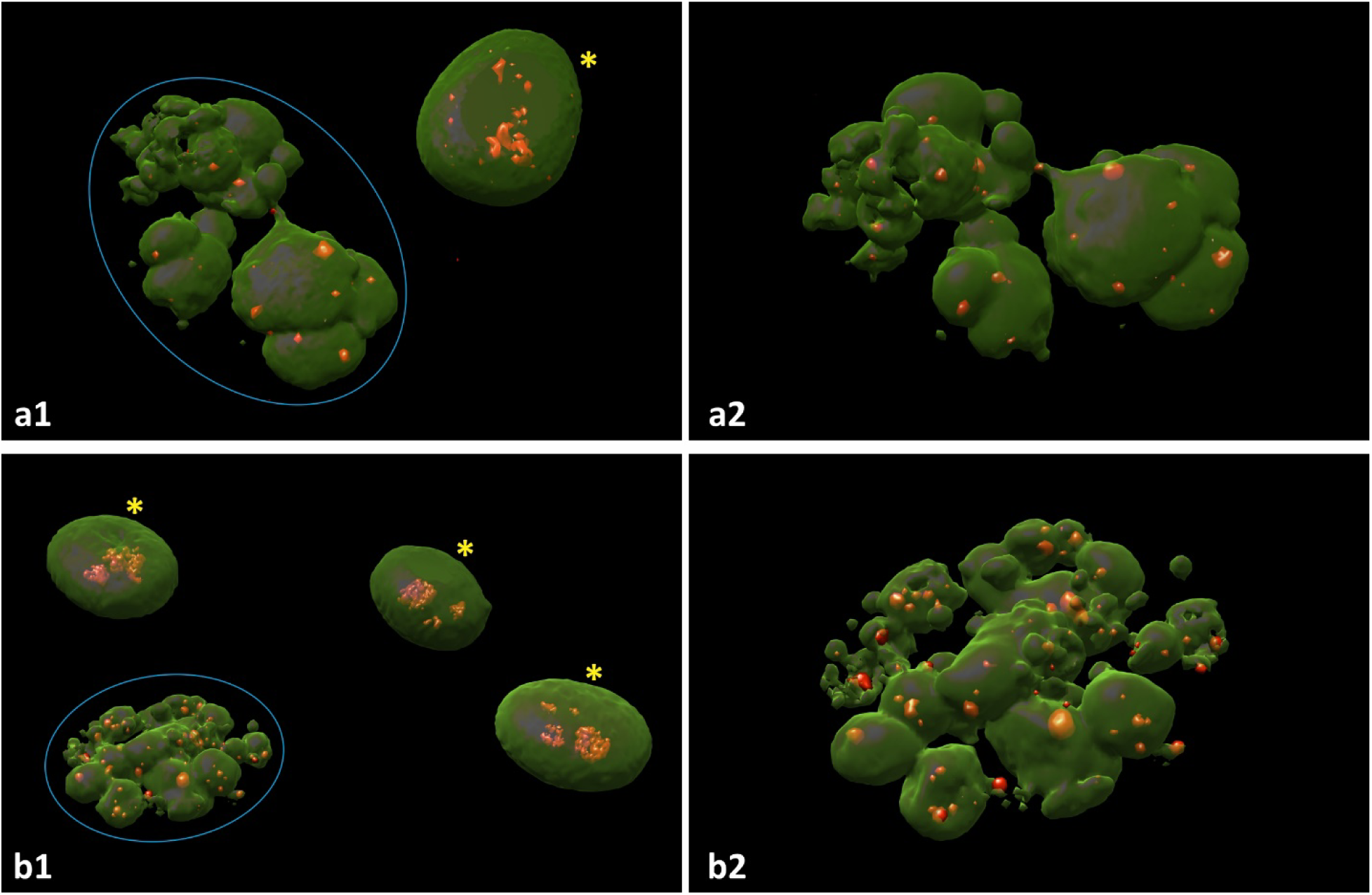
Surviving mononuclear and post-MC multinuclear cells; 72 hours of post-irradiation image acquisition after 30 Gy γ-irradiation; anti-UBTF (a1, a2) and anti-fibrillarin (b1, b2) immunostaining with 3D models (UCSF Chimera). Both labels used AlexaFluor594, retaining red coloration in models. (a1) Low-magnification view shows surviving mononuclear cell (yellow star) and multinucleated/post-MC cells. Surviving mononuclear cell reveals asymmetrically enlarged UBTF-positive bodies and chain-like smaller structures, versus chaotically dispersed round UBTF-positive foci of varying size in nuclear fragments (26, a2, zoomed image). (b1) Low magnification shows similar cell populations. Surviving mononuclear cells retain cord- like fibrillarin organization, while multinucleated cells (26, b2, zoomed image) exhibit disorganized, rounded fibrillarin-positive structures.

The spatial interaction between intra-nucleolar structures marked by UBTF and histone H2B- GFP can be especially well recognized using 3D models of the irradiated nucleoli subjected to post-irradiation acquisition over 48-72 hours. Significantly, light zones exposed on PC images (SFig. 33) match perfectly with the shape, size, quantity, and intra-nucleolar distribution of enlarged but less numerous FCs which are permanently contacting GFP-positive ICC and PCC. Definitely, UBTF-positive FCs never acquire a cap-like structure, being inserted inside the interface area between the nucleolus and the PCC shell. As the compaction of NAC stably increased to this time, the intimate link between PCC, ICC, and UBTF-positive FCs became strictly prominent (Fig. 15, 16, 21-23). The number of nuclei with enlarged to giant sizes UBTF containing spheroids, disposed to the nucleolar periphery, clearly increased during 48 - 72 hours of post-irradiation time. Corresponding 2D images and 3D models particularly represent visual proof of the unity between histone H2B-GFP-positive NAC and FCs. Such a dramatic enlargement of UBTF-positive spheres, accompanied by a strongly increased compaction of ICC, makes the integration of giant FCs into the NAC system well distinguishable. In cases when ICC clump links to PCC shell, one can conclude that γ-irradiation does not disintegrate FCs and the NAC unit system. Importantly, even after 72 hours of post-irradiation time, we never observed the classical pattern of nucleolar segregation i.e. transformation of FC and DFC into UBTF- and fibrillarin-positive nucleolar caps [61, 62]. Interestingly, during 48 and 72 hours of post-radiation period we noted emerging of smaller UBTF-positive structures located in neighborhood with enlarged ones. These tiny structures were profoundly demonstrated on 3D models generated using AutoCAD software (Fig. 23).

Using 2D imaging and 3D reconstruction of immunolabeled fibrillarin-containing DFC, we found that intranucleolar deposits of this protein largely maintain a cord-like organization. As mentioned in the majority of nucleoli submitted to 48 - 72 hours of post-irradiation acquisition, we observed the emergence of giant UBTF-positive FC. Corresponding 3D models clearly expressing different patterns of UBTF and fibrillarin spatial redistribution. Because, by both immunolabellings we did not reveal the transformation of UBTF-positive FCs and-fibrillarin positive DFC into cap-like structures we consider these nucleolar modifications rather as intermediate forms called pre-segregated nucleoli [61].

## 4. Discussion

In this study, we examined nucleolar and nuclear architectural responses to high-dose γ- irradiation, focusing particularly on NAC dynamics and the spatial behavior of FC/DFC assembly. While chemical agents inhibiting ribosomal rRNA synthesis have provided significant insights into nucleolar remodeling [61, 62, 68–81], the structural consequences of physical stressors such as ionizing radiation remain less understood. Here, by employing live-cell 2D imaging combined with fixed-cell 3D reconstruction, we address the gap regarding how higher- than-SBRT doses influence nucleolar structure during the progression of MC and apoptosis.

Unlike chemical inhibition (e.g., Actinomycin D, AMD), which induces classic nucleolar segregation with characteristic crescent-shaped caps, γ-irradiation elicited a distinctly different nucleolar response. Irradiated nucleoli retained enlarged, irregular shapes without typical segregation morphology. Enlarged UBTF-positive FCs were observed alongside smaller UBTF foci, indicating partial segregation or pre-segregated states rather than complete FC relocation. Importantly, despite extensive nuclear deformation, multinucleation, and apoptotic signs, UBTF and fibrillarin remained closely associated within FC/DFC complexes. Fibrillarin preserved its distinctive cord-like organization, demonstrating that γ-irradiation does not fully disassemble these structural nucleolar domains. Thus, our findings reveal significant resilience of nucleolar architecture to severe physical DNA damage.

The stability of FC/DFC association and their persistent linkage to NAC during genotoxic stress suggests an intrinsic protective role provided by nucleolar chromatin compaction. Given that irradiation likely induces rDNA fragmentation, the persistence of small UBTF-positive foci may reflect asymmetrical DNA degradation producing heterogeneous rDNA fragments. If so, these fragments remain associated with UBTF, generating numerous small pseudo-FCs, which supports the concept of nucleoli serving as resilient hubs that maintain structural and possibly functional integrity under physical stress conditions.

### 4.1. Multinucleation and Asynchronous Apoptosis Dynamics

Time-lapse imaging documented multinucleation through sequential nuclear invagination, lobulation, and fragmentation, ultimately culminating in asynchronous apoptotic degeneration. Initially, irradiated nuclei developed intense multifocal invaginations, leading to irregularly shaped nuclei that subsequently fragmented into micronuclei. This nuclear fragmentation likely results from incorrect chromosomal segregation during aberrant nuclear divisions, suggesting amitotic mechanisms. Some micronuclei retained nucleoli with prominent UBTF and fibrillarin- positive regions, while others were anucleolated, likely due to stochastic inheritance of NORs. The presence of both nucleolated and anucleolated micronuclei underscores heterogeneous chromosome distribution events during nuclear fragmentation.

Interestingly, apoptotic changes proceeded asynchronously across micronuclei within the same multinucleated cells. Some micronuclei exhibited “hyper-condensed” perinuclear chromatin blocks - a phenomenon called chromatin marginalization, indicative of advanced apoptosis [99], whereas neighboring nuclei showed relatively normal nucleolar structure. This heterogeneity emphasizes complex nucleolar signaling and regulatory mechanisms activated upon severe DNA damage. Thus, multinucleation and subsequent asynchronous apoptosis represent hallmark features of irradiation-induced MC, with nucleolar remodeling closely coupled yet delayed relative to broader nuclear disintegration.

### 4.2. Nucleolar Remodeling Precedes MC and Apoptosis

A notable finding was the early nucleolar remodeling preceding extensive nuclear alterations. Shortly after irradiation, ICC displayed gradual compaction and redistribution toward the PCC. This behavior resembled nucleolar dynamics triggered by chemical inhibitors [61], suggesting common initial chromatin-based nucleolar stress responses. However, major nuclear deformation and multinucleation were prominent only at later stages (24–48 hours of post-irradiation). Even severely damaged or multinucleated cells maintained bright UBTF and fibrillarin signals, indicating delayed nucleolar disassembly compared to nuclear degradation. These findings highlight that early nucleolar chromatin dynamics may act upstream of MC, setting the stage for subsequent nuclear catastrophe.

Additionally, we observed the formation of optically clear nucleolar regions reminiscent of the previously described “nucleolinus” by Love and Soriano [100, 101]. These giant UBTF-positive FCs likely represent fused FC units resulting from transcriptional inactivation of rDNA. Unlike AMD-induced nucleolar caps, however, these structures remained embedded within the nucleolus without peripheral segregation, suggesting distinct pathways of nucleolar remodeling between chemical and physical DNA damage. Further live-cell studies with RNA Polymerase I (Pol I) -tagged models will clarify the real-time dynamics underlying this unusual structural response.

### 4.3. NAC–FC/DFC Interaction and Structural Integrity

Our results underscore the critical role of NAC, including ICC and PCC sub-compartments, in nucleolar structural responses to genotoxic stress. Although NAC does not contain actively transcribed rDNA, its compact nucleosomal organization facilitates nucleolar compartmentalization. Structural remodeling of NAC following irradiation closely mirrored AMD-induced chromatin rearrangements, including gradual ICC contraction and spatial repositioning toward PCC. Notably, even when severely damaged, ICC retained sufficient mobility to facilitate the spatial redistribution of FC/DFC assemblies without fully segregating them.

This preserved association suggests that despite γ-irradiation-induced DNA breaks, NAC maintains contractile properties essential for sustaining nucleolar organization. Time-lapse observations supported ICC’s active role in nucleolar restructuring, particularly during initial stress response phases. Only later, when nuclear deformation became evident, did ICC form large immobile clumps fully integrated into PCC, further illustrating NAC’s role in nucleolar resilience.

### 4.4. Differential Behavior of UBTF and Fibrillarin under γ-Irradiation

Comparative analysis between chemically and physically induced nucleolar stress highlighted crucial differences in FC/DFC remodeling. Chemical inhibitors like AMD trigger FC/DFC complexes to form compact peripheral nucleolar caps. In contrast, γ-irradiation maintained irregularly shaped nucleoli without cap formation. By this UBTF-positive FCs alongside with fibrillarin-containing DFC behave differentially if compared with action of chemical inhibitors.

UBTF-positive FCs enlarged but did not segregate fully, with small UBTF-positive foci persisting. Fibrillarin distribution remained consistently cord-like and maintained close FC association even after extensive radiation damage.

These results suggest that although γ-irradiation likely inhibits rRNA transcription similarly to chemical inhibitors, it does so via different molecular mechanisms, preventing full nucleolar segregation. Physical damage appears less capable of completely disrupting protein/rDNA-based scaffolds that anchor FC/DFC assemblies. Naturally, rDNA template folded into structure of UBTF-positive FC must be much more sensitive to γ-irradiation-induced damages than r-genes transcription products, presented by fibrillarin-positive protein/pre-rRNA complex. Likely, emergence of giant FCs alongside with tiny pseudo-FCs may result from r-chromatin asymmetric double strand breakages. Consequently, being a protein/pre-rRNA product of r-genes transcription, DFC, can be considered as less sensitive to γ-irradiation-induced damage of rDNA, hence, doesn’t disrupt. Therefore, we hypothesize that even after 72 hours of post-irradiation period giant FCs can have remained at list partially encased in fibrillarin-positive DFC cords, thus remaining complexed as FC/DFC assembly. Future studies employing fibrillarin-GFP permanently transfected HeLa cell models will further elucidate these observations.

### 4.5. Conclusion and Future Directions

Our findings reveal a distinct nucleolar remodeling pattern following severe physical DNA damage, characterized by incomplete segregation, preserved FC/DFC integration, and pronounced chromatin compaction. This pattern underscores nucleolar resilience, supporting its role as a robust structural and functional biosensor during genotoxic stress. The observed nuclear and nucleolar dynamics, including delayed cellular response in form of MC and asynchronous apoptosis, highlight the complexity of cellular responses to ionizing radiation. Hence our results may offer new insights into nuclear and nucleolar resilience under genotoxic stress and suggest that nucleolar features could serve as biomarkers for DNA damage severity. Additionally, results yielded may have clinical value in the context of developing novel RT strategies through EMC. This study also lays the groundwork for future 4D imaging to dissect real-time nucleolar and NAC dynamics in response to different chemical and physical stressors. Therefore, future investigations using advanced live-cell imaging and functional assays (e.g., detecting rDNA single- and double-strand breakages, measuring rRNA synthesis etc.) will further clarify how physical DNA breaks mechanistically influence nucleolar organization, providing insights beneficial for therapeutic strategies aimed at exploiting nucleolar vulnerability in cancer treatments. All these approaches are in progress now in Carl Zeiss Education and Scientific Center at NVU.

## Acknowledgment

This work was financially supported by the New Vision University (Tbilisi, Georgia). Authors express special gratitude to the University authorities, as well as to the members of the Academic Council for their efficient financial and technical engagement.

## Notes

### Competing Interest Statement

The authors have declared no competing interest.

