## Supplementary Figures for "Nucleolar and Chromatin Remodeling during γ-Irradiation-Induced Mitotic Catastrophe: Live-Cell Imaging Correlated with UBTF and Fibrillarin 3D Redistribution": Supplementary Figures.docx

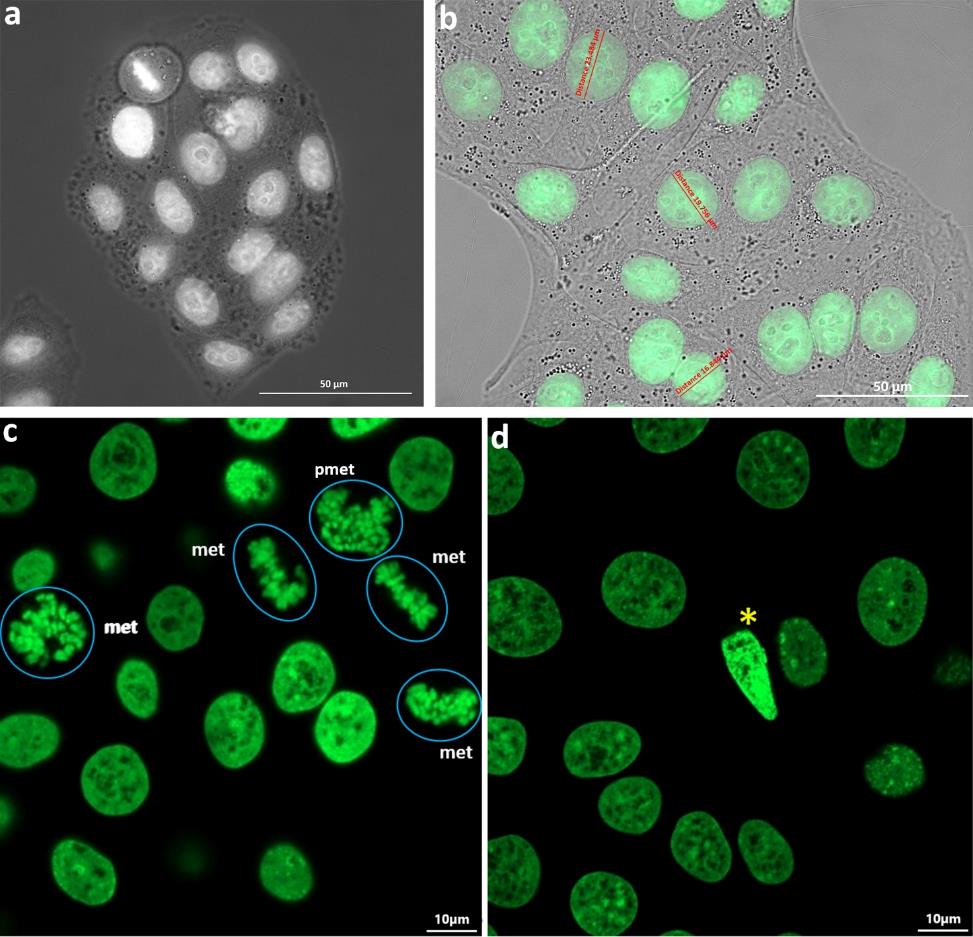

**S1 Figure.** Microstructural features of cultured control HeLa cells stably expressing histone H2B-GFP fusion protein. These cells were taken at low magnification, using phase contrast (PC; 1, a), fluorescent (FL; 1, c, d) and merged PC+FL (1, b) regimes. (b-d) note bright green fluorescence that revealed high resistance to UV-light, retaining sufficient brightness even over 72 hours of post-irradiation image acquisition period. Abundancy of mitotic cells (1, a, c, d) as well as occasional apoptotic cells (1, d, marked by yellow star) were detected.

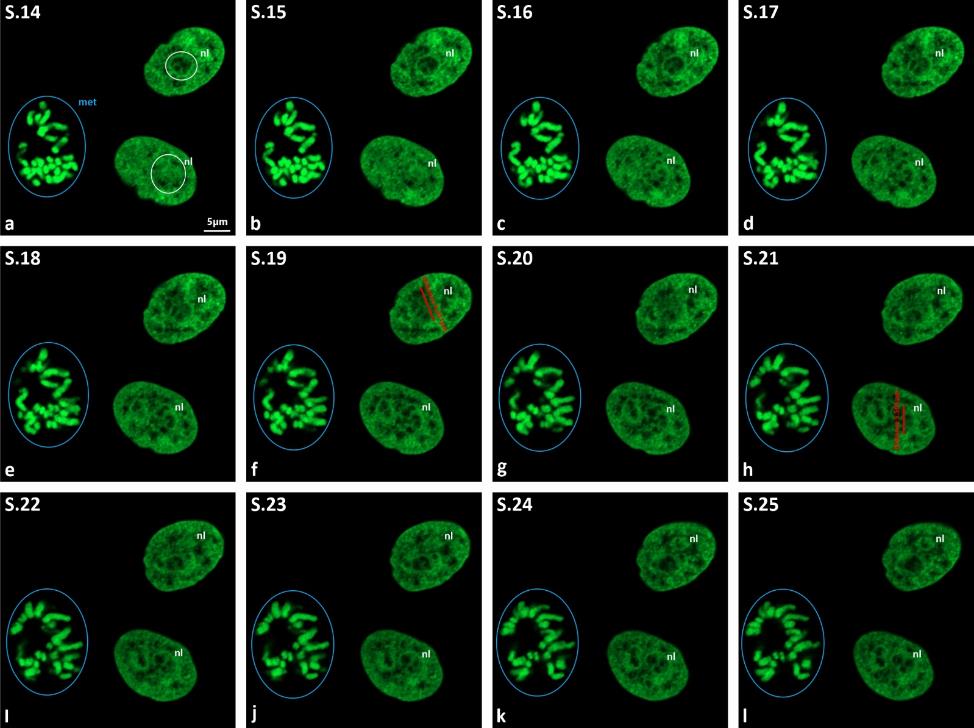

**S2 Figure.** Control: gallery of successive virtual serial sections (S.14-S.25) revealing typical nuclear morphology of interphase mononuclear cells and mitosis (outlined by blue circle). (a-l) Evidence that the majority of histone H2B-GFP transfected HeLa cells uniformly, contain ovoid nuclei with predominantly smooth or slightly wave-like contours, visible on all sections of series presented. Nucleolar territory is outlined by white circles (2, a), while average nucleolar sizes ranged from ~4 to 7 μm (2, f, h). Even at low magnification PCC ring and ICC network were profoundly seen. met – metaphase.

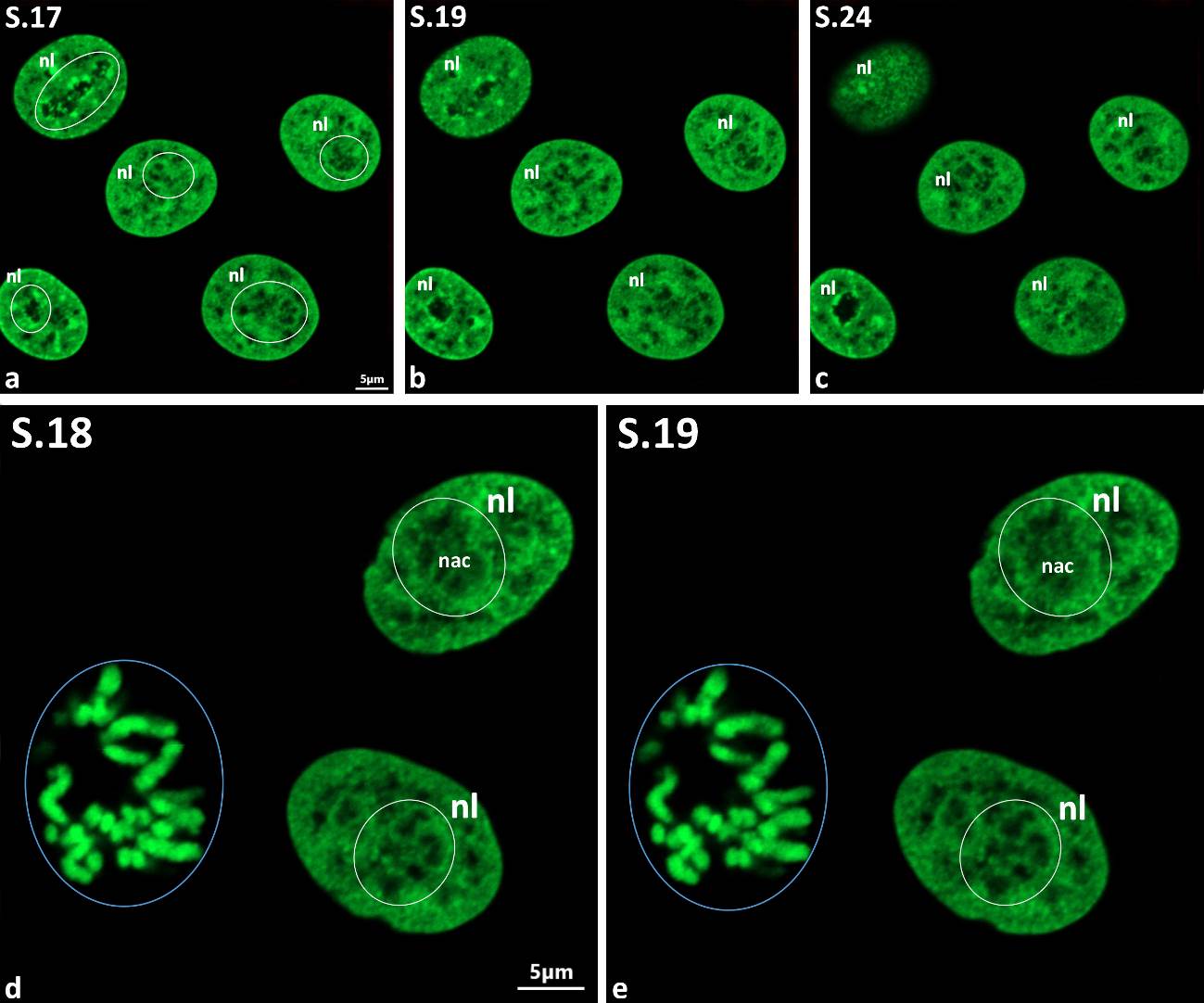

**S3 Figure.** Control: additional evidence that control mononuclear cells are uniform in global nuclear morphology and nucleolar microstructure. (a-c) Gallery of virtual sections (S.17, S.19, S.24) showing nuclei nucleoli (outlined with white circles) stably revealing roundish shape, smooth outlines and profound ICC network; Note also prominent PCC ring. (d, e) Images presented on 2, a-l zoomed for better recognition of NAC system. Nucleolar territory is outlined by white circles. Mitotic cell is outlined by blue circle. NAC – nucleolus associated chromatin; nl – nucleolar territory.

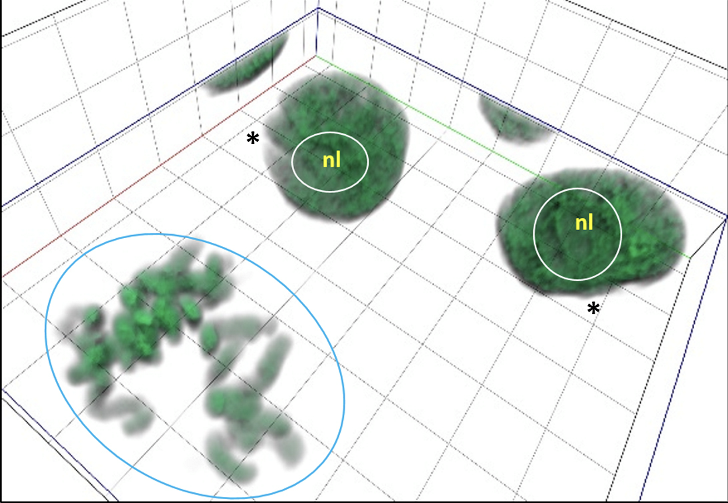

**S4 Figure.** Control: 3D model of nuclei generated by ZEN3.0 software, using aligned: (i) volume rendering; (ii) transparency mode by (iii) variable thresholds (correspond to mitotic cell and nuclei depicted on 2, a-l and 3, d, e). Nucleolar territories is outlined by white circles. Note that nuclear periphery shows small indentations (marked by stars). Abbreviations as on previous supplementary figures.

**S5 Figure.** Control: ant-UBTF immunostaining. Evidence that nuclei with profoundly more irregular (than nuclei demonstrated on 4) contours are also present in cultures used. Although more deformed nuclear shape the anti-UBTF labeling (red) revealed classical chain-like organization (marked by red arrows). Anaphase cell is outlined by blue circle. Abbreviations as on previous supplementary figures.

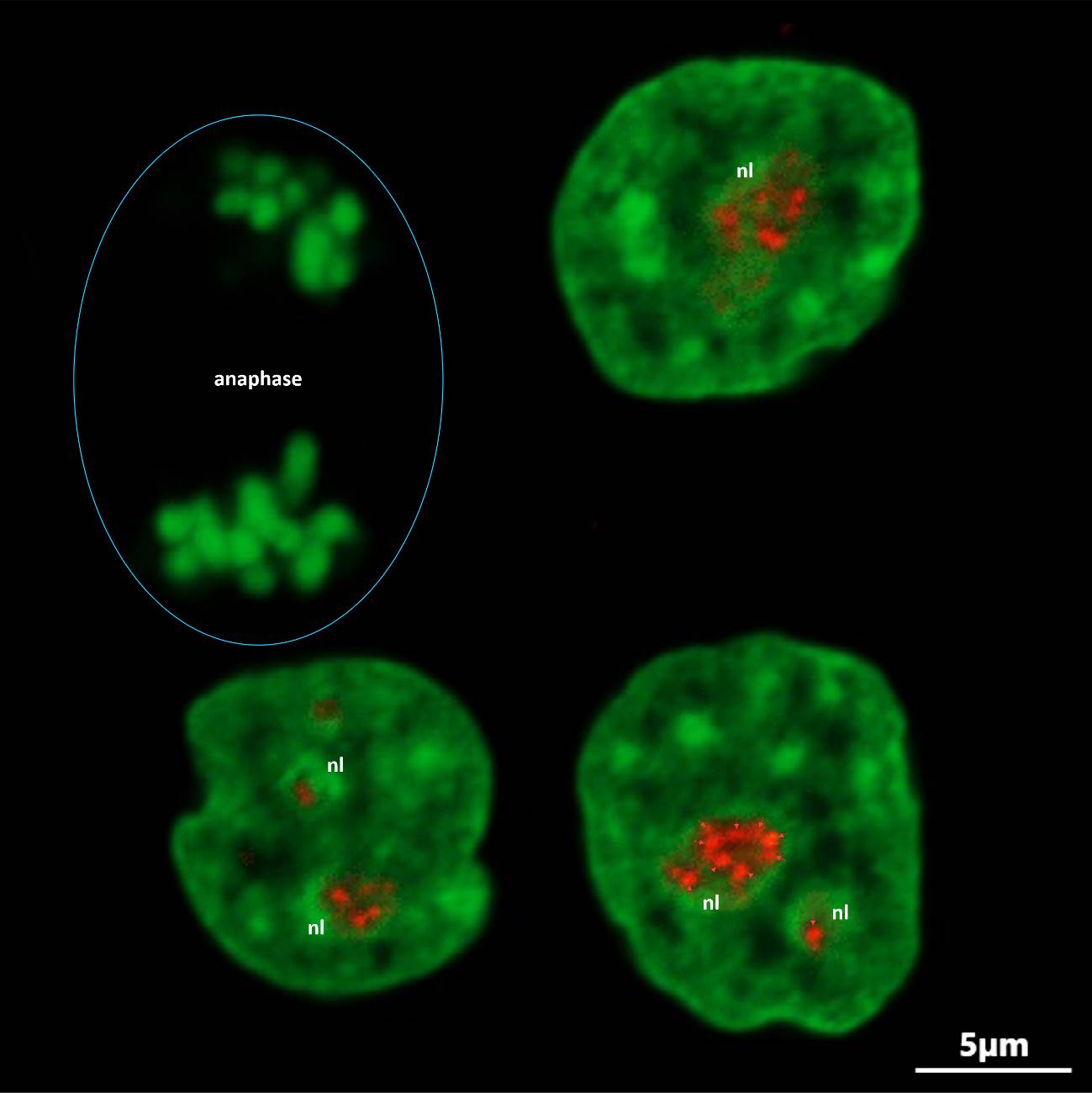

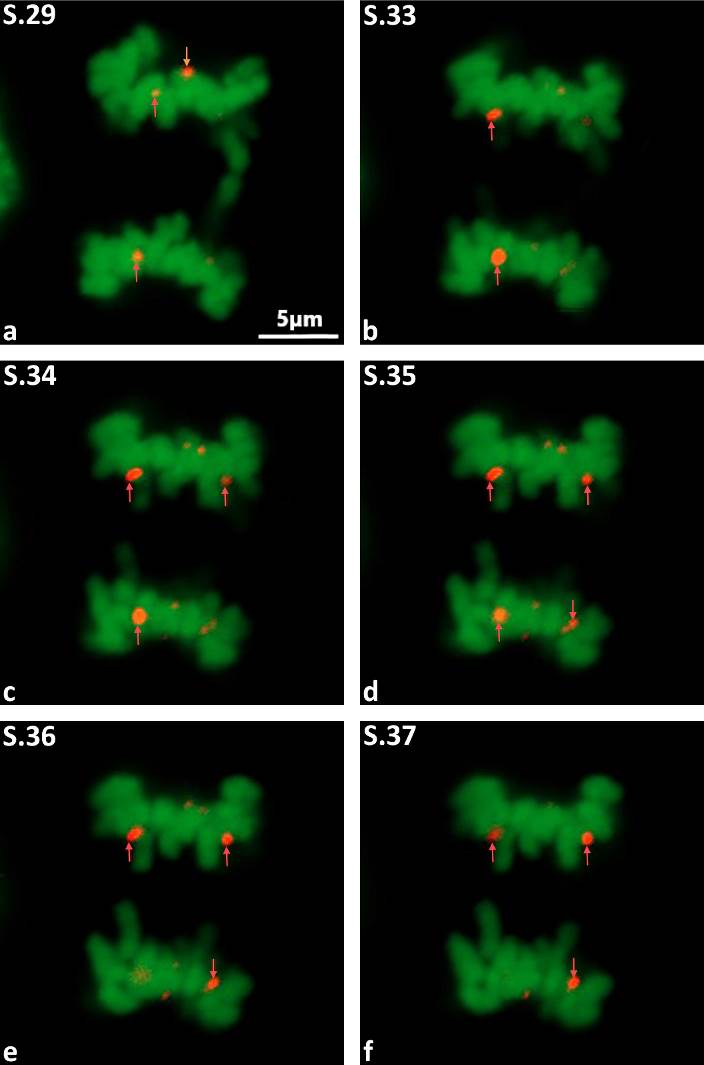

**S6 Figure.** Control: gallery of virtual sections (S.29, S.33-S.37) showing UBTF-positive NORs (marked by red stars).

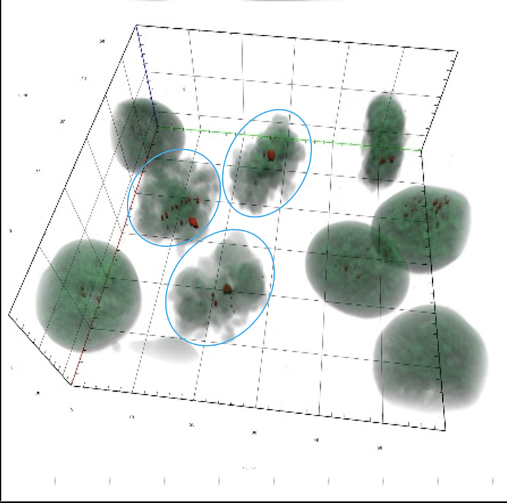

**S7 Figure.** Control: 3D models (ZEN3.0) showing UBTF-positive NORs. Mitotic cells are outlined by blue circles). Note the different 3D pattern of UBTF-positive structures in interphase cells. Evidence that UBTF is exclusively localized in NORs and their interphase counterparts, i.e. FC.

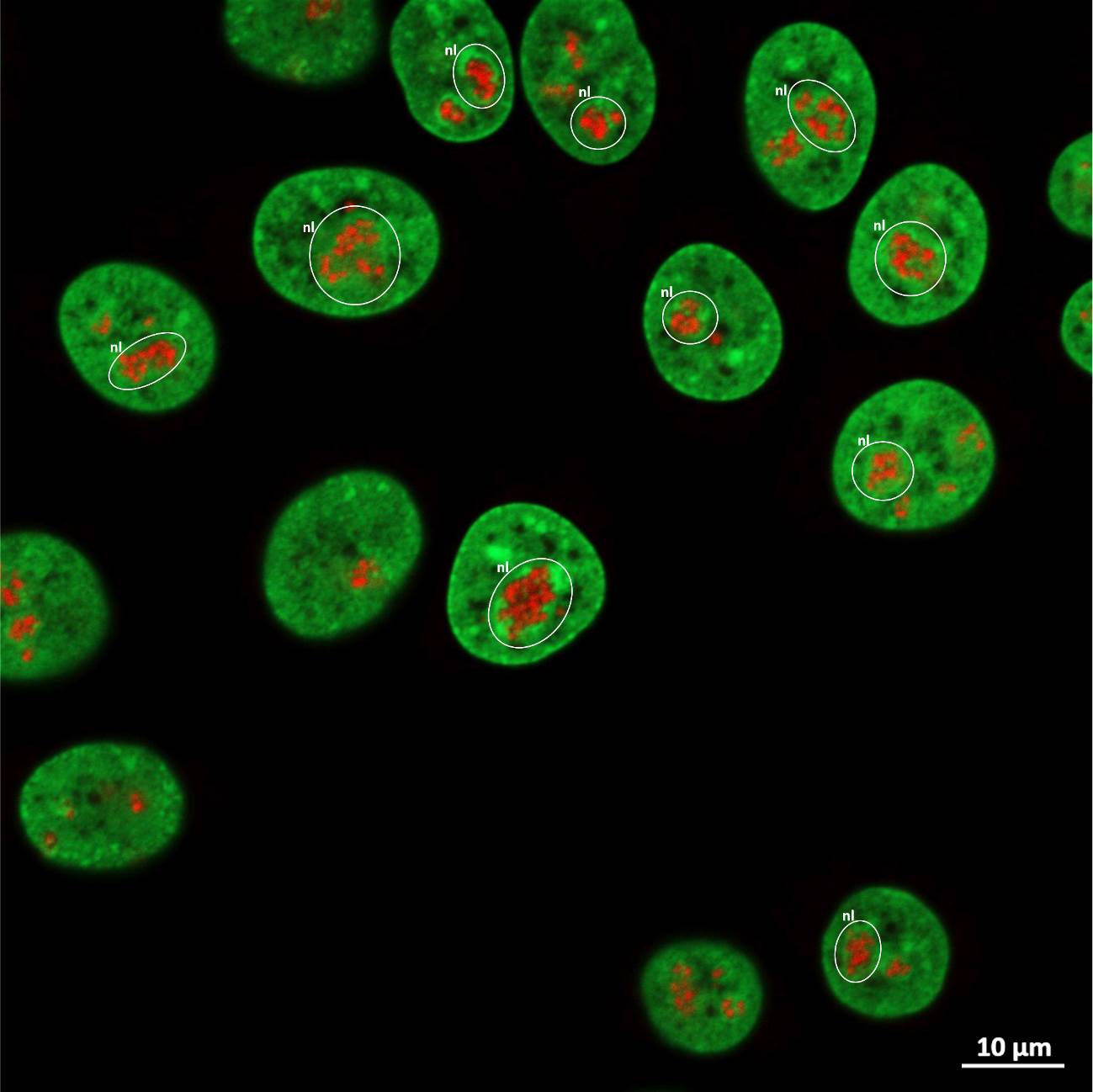

**S8 Figure.** Control: anti-fibrtillarin immunolabeling. Demonstration of nuclear and nucleolar microstructure taken at low magnification to prove that majority of cells are mononuclear, while nuclear contours are smooth. Nucleolar territories are outlined by white circles. Fibrillarin label (red) looks like uniform in all nuclei. Abbreviations as on previous figures.

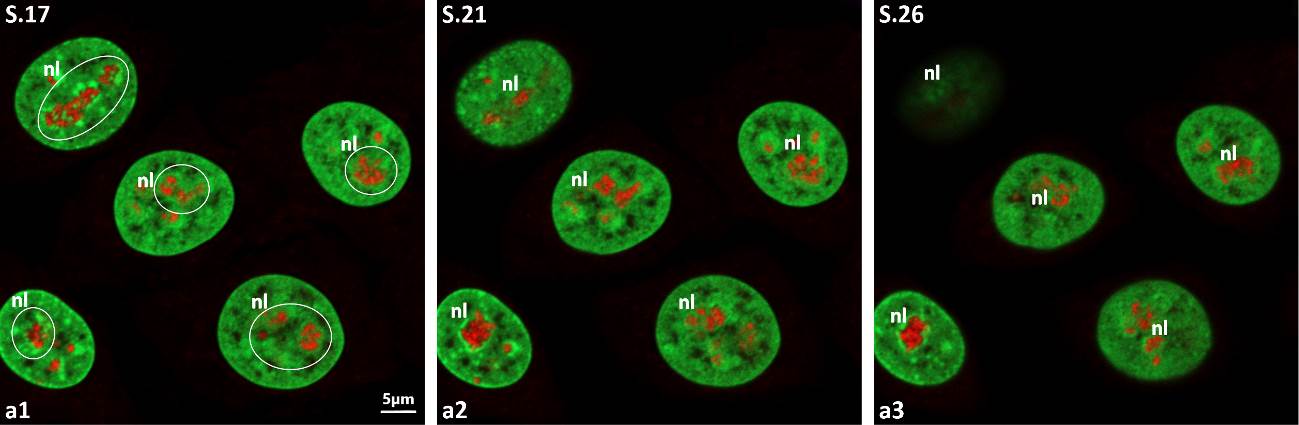

**S9 Figure.** Control: nuclear and nucleolar microstructure after anti-fibrillarin immunolabeling; three non-successive images of virtual planes (S.17, S.21, S.26) extracted from complete series and zoomed for better recognition the uniform spatial arrangement of fibrillary-positive structures (red). Nucleolar territories are outlined by white circles. On all planes anti-fibrillarin label looks like gathered in cord-like fashion. Abbreviations as on previous figures.

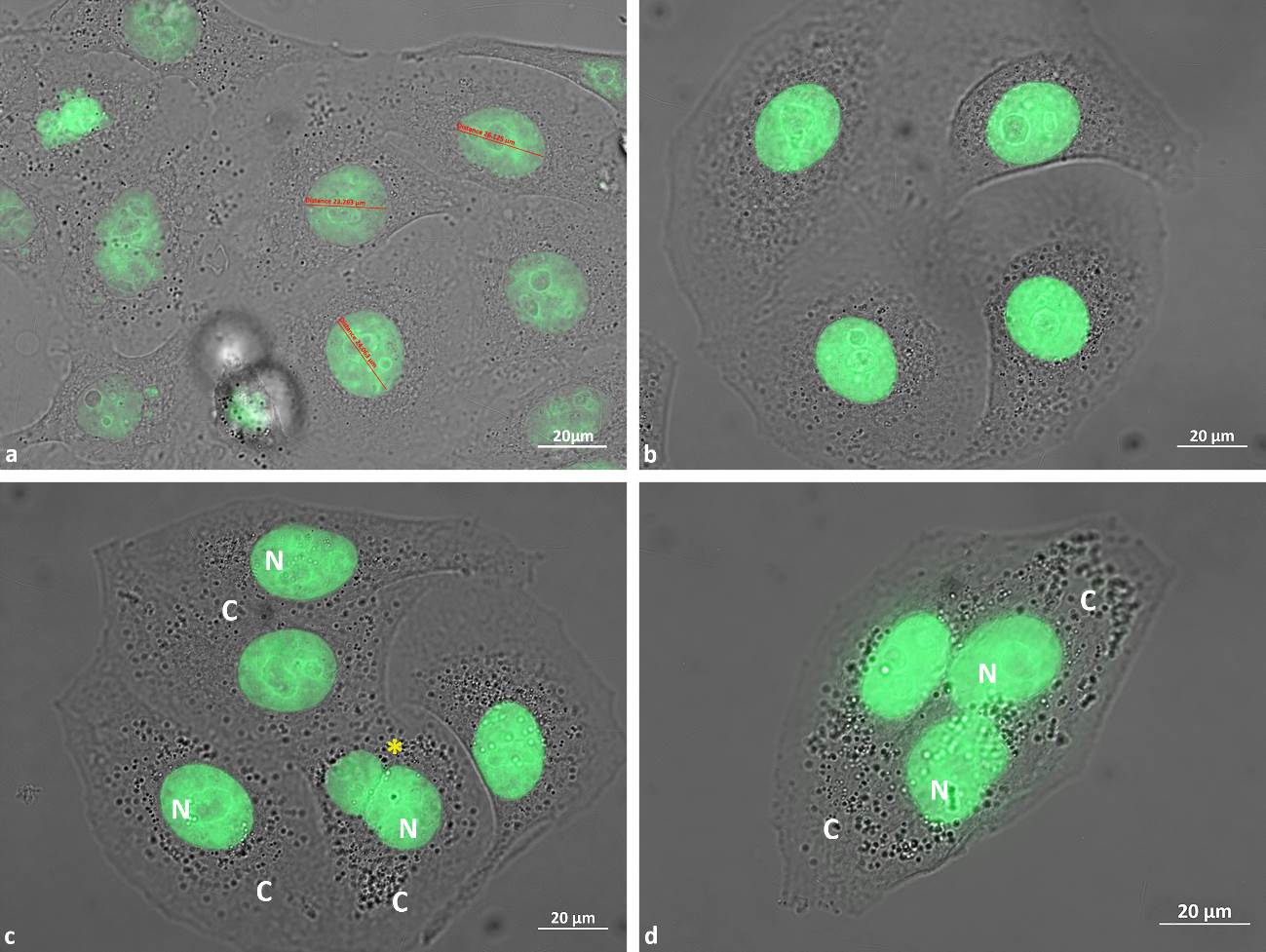

**S10 Figure.** (a-d) 30 Gy post-γ-irradiation image acquisition during 48 hours. These images were taken at low magnification using merged PC and FL regimes to prove mononuclear and multinuclear examples. Importantly, mitotic cells can be seen quite often (10, a; upper left corner of the image). Even after 48 hours of post-irradiation acquisition the plethora of cells retain mononuclear morphology, while nuclear appearance resembling control, despite slightly increased average diameters (10, a); note smooth nuclear outlines (10, a, b) and well distinguishable nucleolar ICC inclusions (10, b; upper right nucleus). (c, d) Emergence of multinucleated cells: demonstration of bi-nuclear (10, c; marked by yellow star) and three-nuclear cells (10, d). In all cases microstructure of the cytoplasm (C) was largely looked like in control samples (compare with S1 Fig.). C – cytoplasm. Other abbreviations as on previous figures.

**S11 Figure.** (a-d) 30 Gy post-γ-irradiation image acquisition during 72 hours. On these figures PC images were also merged with FL ones, hence evidencing presence of mononuclear and multinuclear cells. Mitotic cells can be also seen (11, a; taken in blue circle). Remarkably, after 72 hours of post-irradiation acquisition sufficient number of cells yet retain mononuclear morphology and smooth nuclear outlines (11, b); in parallel, cells with drastically deformed and lobed nuclei (outlined with blue circles) abundantly appeared (11, b, c). Note that same images witnessing micronucleation process and appearance of multinucleated cells (large multinuclear cell on the right side of image). (d) demonstration of giant post-MC multinuclear cells containing simultaneously several larger nuclei with prominently seen nucleoli (outlined by blue circle) and numerous much smaller micronucleoli still emitting GFP signal. Note also, that some cells adopted peculiarly deformed cytoplasmic contours (11, c).

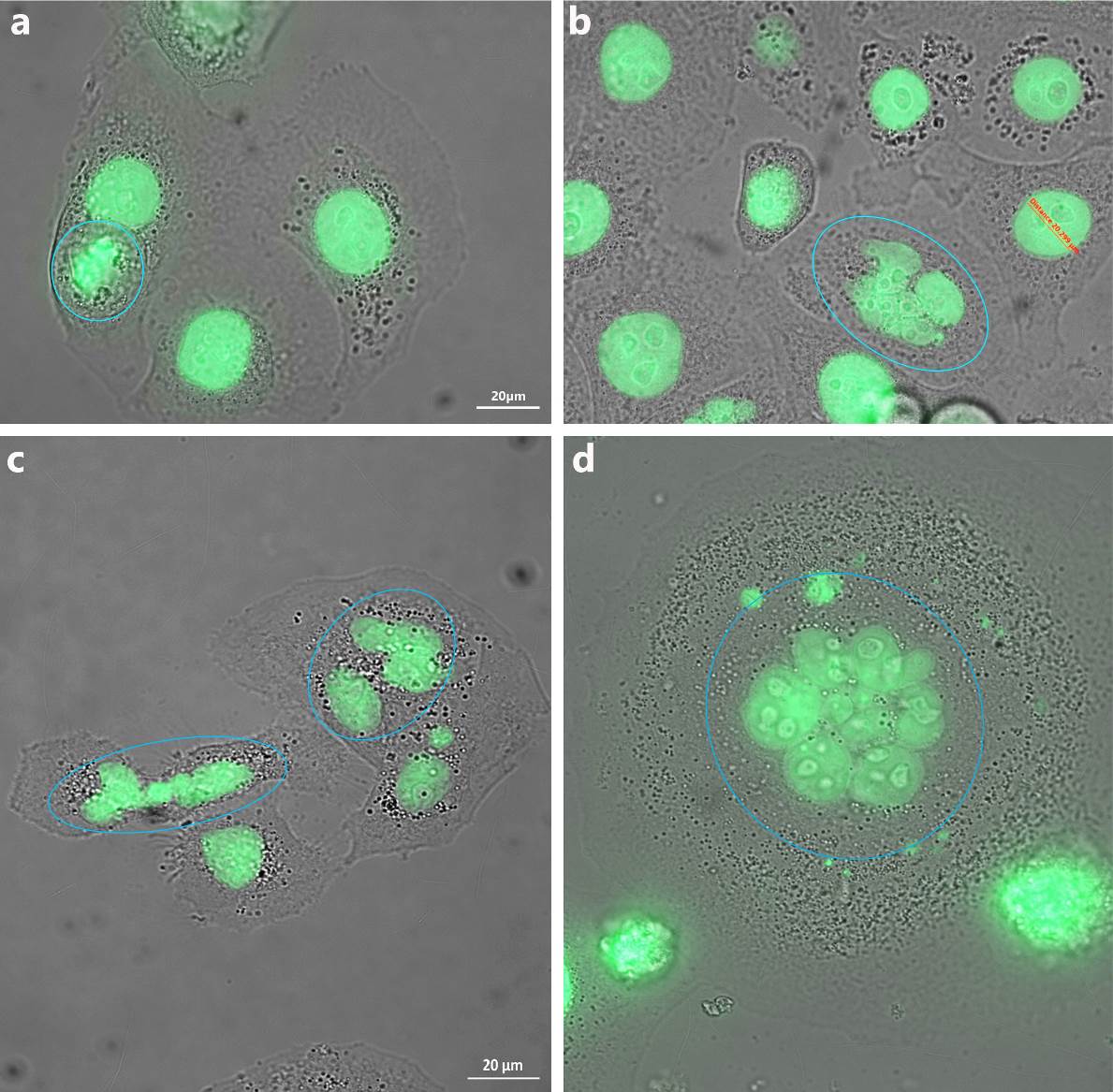

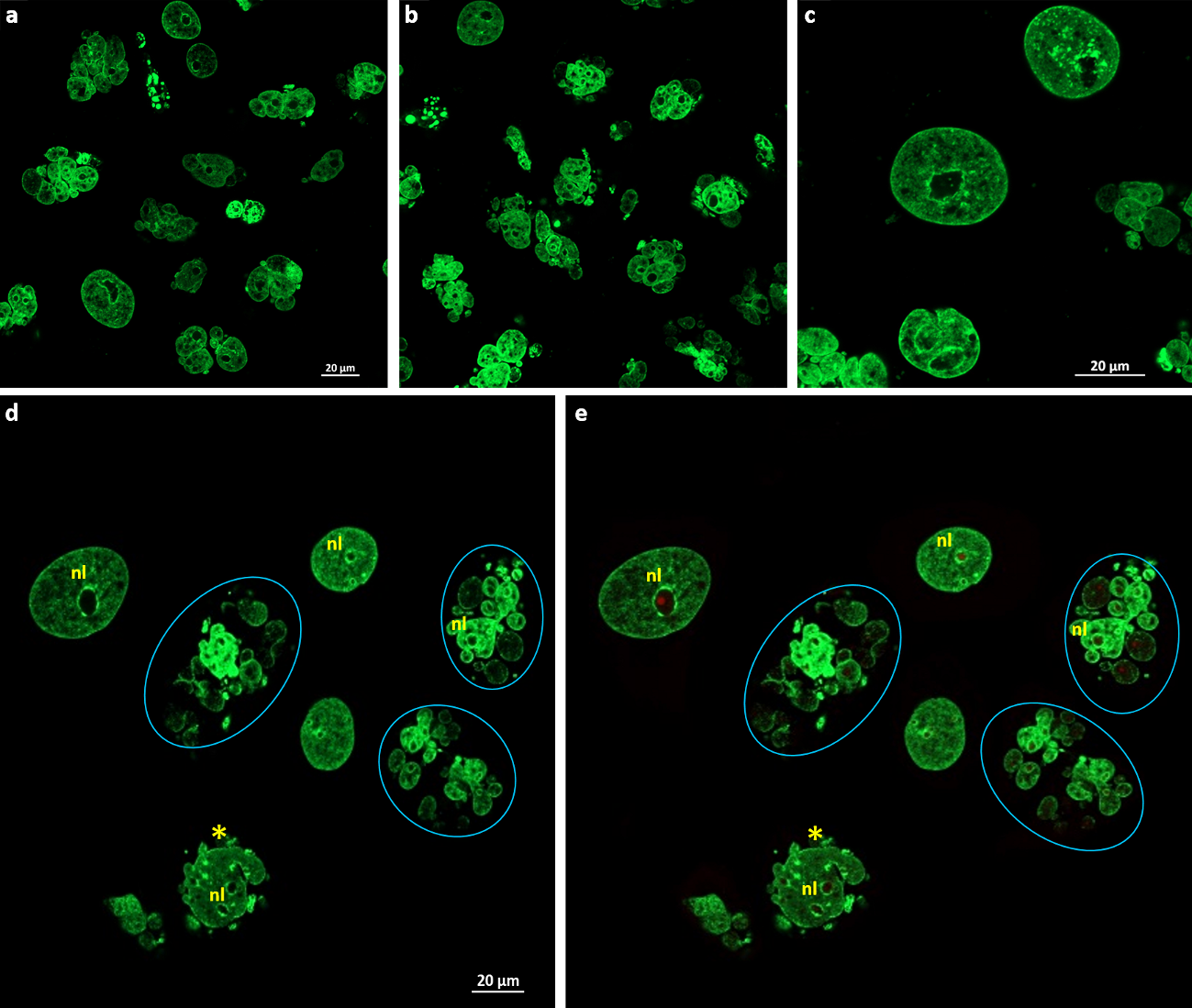

**S12 Figure.** 30 Gy post-γ-irradiation image acquisition during 72 hours. (a-e) Images taken either in FL regime only (12, a-d), or after ant-UBTF immunolabeling (12, e; red). Demonstration of post-MC multinucleation/micronucleation process and massive cellular death. Multinuclear cells outlined by blue circles (12, d, e); note morphological signs of asynchronous apoptosis. Mononuclear cells with roundish, smoothly outlined nuclei also present; meanwhile, some nuclei reveal deep invaginations, witnessing initial amitotic mechanism of multinucleation/micronucleation (12, c-e). Note also large UBTF-positive NCs in mononuclear cells and in post-MC nuclear fragments (12, d, e). Abbreviations as on previous figures.

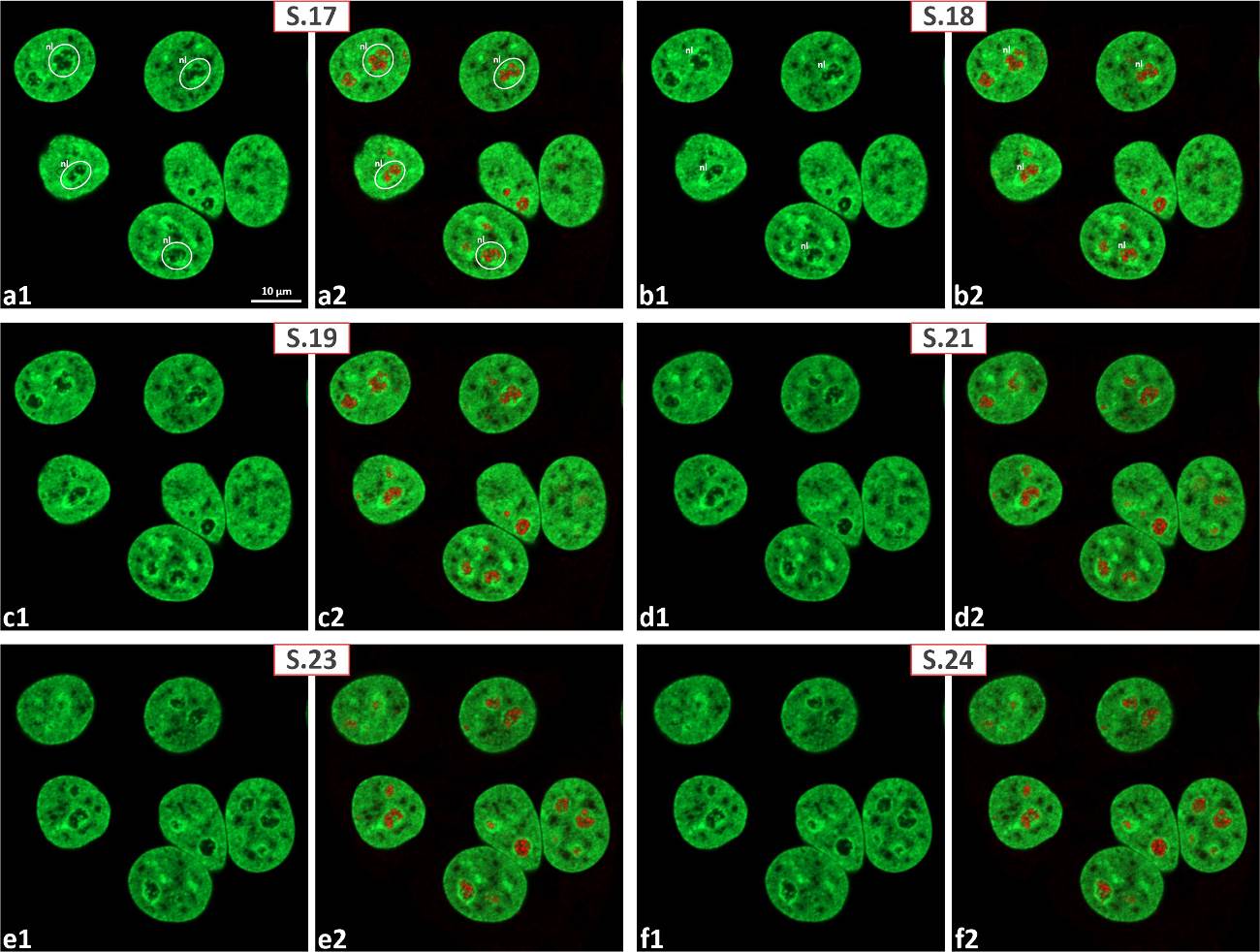

**S13 Figure.** 30 Gy post-γ-irradiation image acquisition during 12 hours. (a1-f2) The gallery of virtual serial (non-successive) sections (S.17-S.19 and S.21-S.24) after anti-UBTF immunolabeling. Nucleolar territory was outlined by white circles. These images were demonstrated without zooming to prove that beside obvious majority of mononuclear cells, three-nuclear cells can be rarely appeared (lower right part of images); presumably a consequence of early irradiation-induced damage and multinucleation. Note close to control nuclear and nucleolar microstructure in majority of mononuclear cells. (a1-f1) GFP fluorescence only: even at low magnification GFP-positive clumps can be easily distinguished, witnessing early signs of irradiation-induced ICC coarsening. (a2-f2) After merging with anti-UBTF immunolabeled images it became clear that chain-like appearance and integration of UBTF-positive structures into NAC system remained largely as in control. Note also that nucleolar microstructure in three-nucleated cell largely resembles those in control preparations. Abbreviations as on previous figures.

**S14 Figure.** 30 Gy post-γ-irradiation image acquisition during 12 hours. (a1-f2) Anti-fibrillarin immunolabeling. The gallery of non-successive virtual serial sections (S.22, 23 and S.25-S.28). Nucleolar territory was outlined by white circles. As in previous figure, these images were demonstrated without zooming to prove presence of bi-nuclear cells (lower left part of images) among majority of mononuclear cells. Demonstration of similar to control nuclear and nucleolar microstructure in majority of mononuclear cells. (a1-f1) Showing GFP fluorescence only. (a2-f2) Merging with anti-fibrillarin immunolabel confirms cord-like appearance of fibrillarin-positive structures (red). In general, nuclear and nucleolar structure in bi-nuclear cell is same as in control samples. Abbreviations as on previous figures.

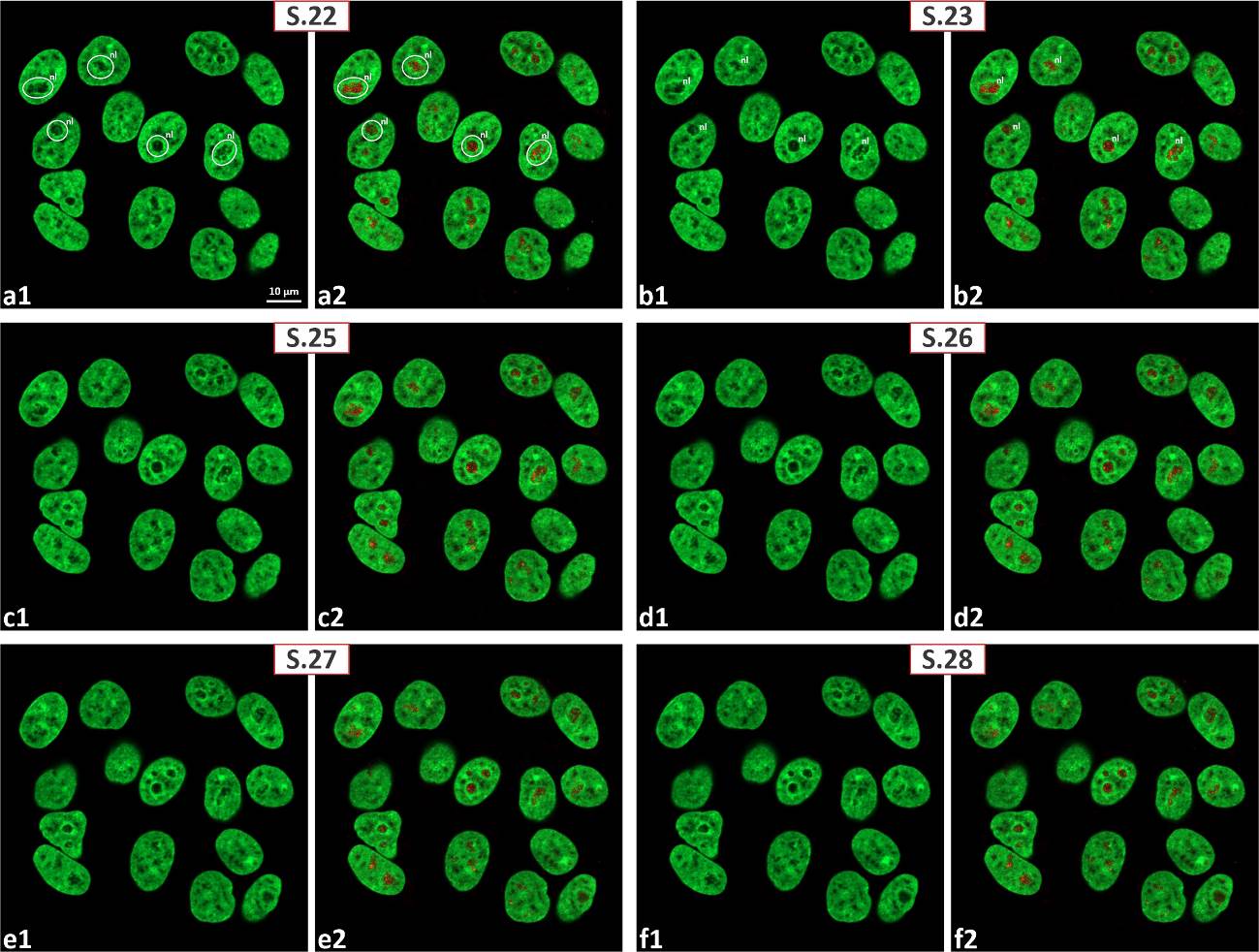

**S15 Figure.** 30 Gy post-γ-irradiation image acquisition during 24 hours. (a1-b2) Anti-UBTF immunolabeling (red). The gallery of non-successive virtual sections extracted from complete series. Nucleolar territory is outlined by white circles. Evidence that even after 24 hours of post-irradiation image acquisition predominance of mononuclear cells is undeniable. (a1, b1) GFP fluorescence only; note clear signs of thickening of PCC and coarsening/clumping of ICC. (a2, b2) After merging with labeled images asymmetrical enlargement of UBTF-positive structures (as consequence of γ-irradiation) became obvious (marked by red arrows). Abbreviations as on previous figures.

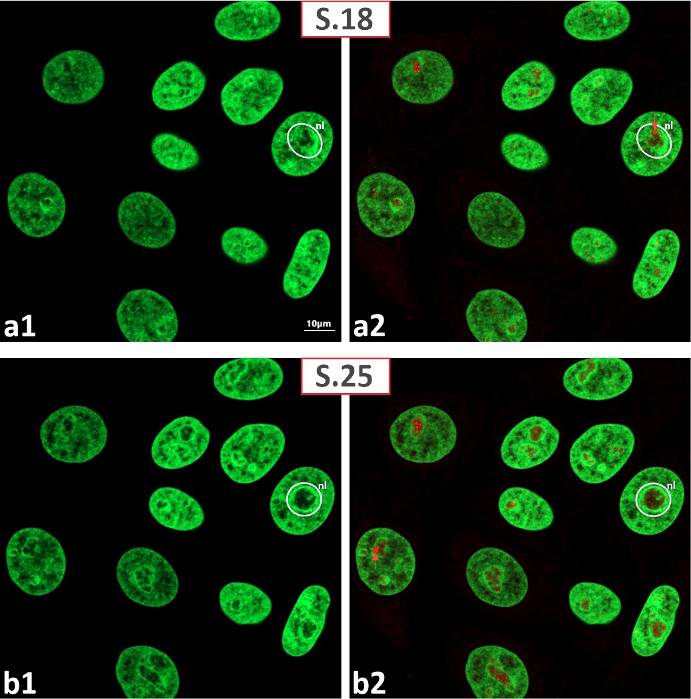

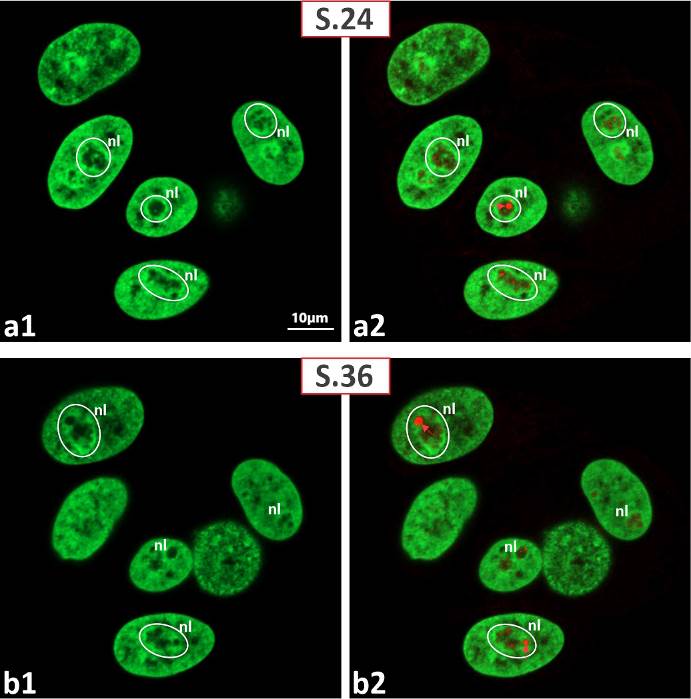

**S16 Figure.** 30 Gy post-γ-irradiation image acquisition during 24 hours. (a1-b2) Anti-UBTF immunolabeling. Zoomed images confirming all previously registered (S15 Fig.) changes at higher magnification. Nucleolar territory was outlined by white circles. Predominance of mononuclear cells was obvious. (a1, b1) GFP fluorescence only; note clear signs of thickening of PCC and coarsening/clumping of ICC. (a2, b2) Merged images prove asymmetrical enlargement of UBTF-positive structures (red, marked by red arrow). Abbreviations as on previous figures.

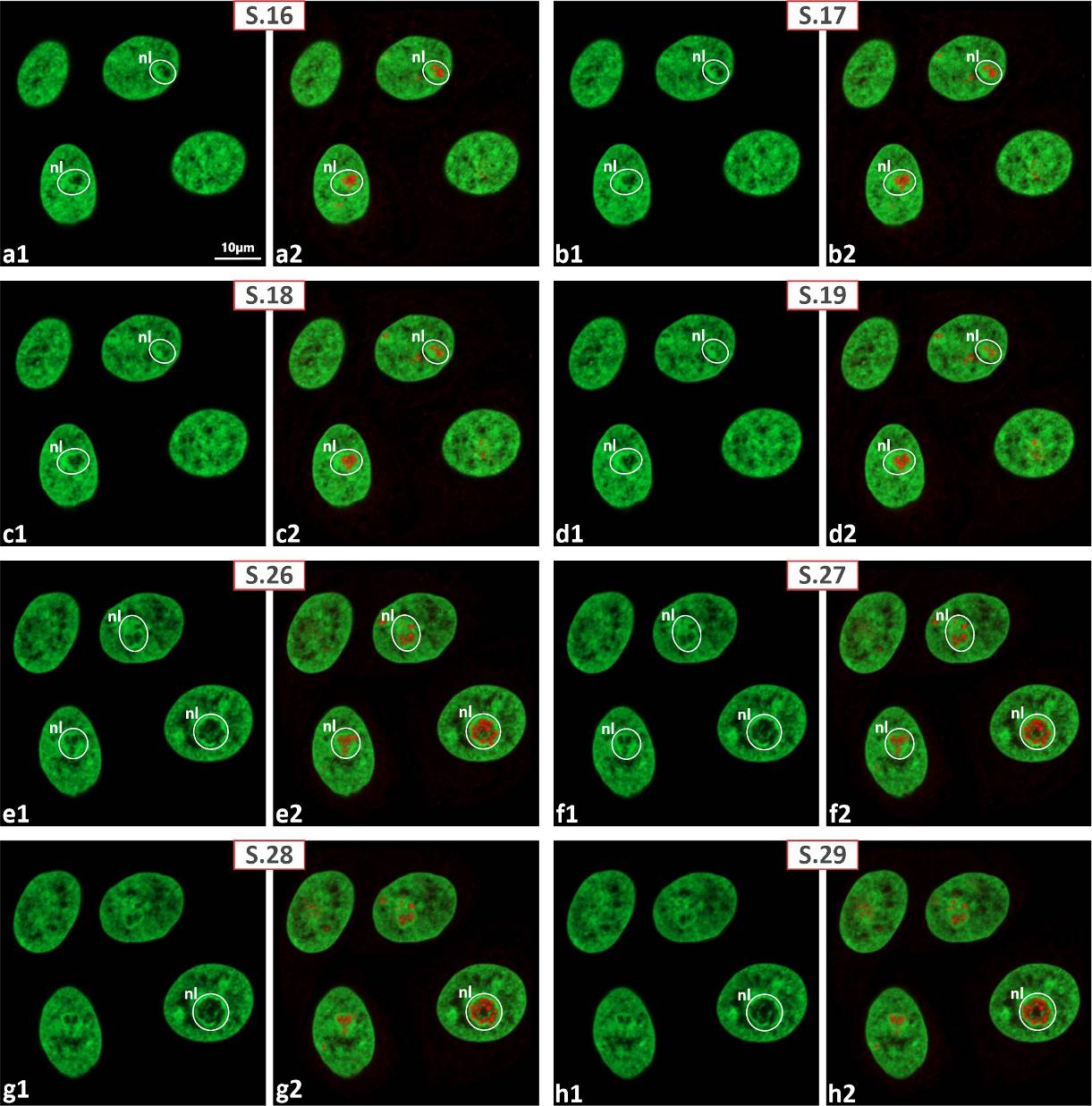

**S17 Figure.** 30 Gy post-γ-irradiation image acquisition during 24 hours. (a1-h2) Anti-fibrillarin immunolabeling. The gallery of non-successful virtual serial sections (S.16-S.19; S.26-S.29). Nucleolar territory is outlined by white circles. This image confirms predominance of mononuclear cells. (a1-h1) GFP fluorescence only; note thickening of PCC and coarsening/clumping of ICC. (a2-h2) Evidence that fibrillarin-positive structures (red) retain cord-like appearance. Abbreviations as on previous figures.

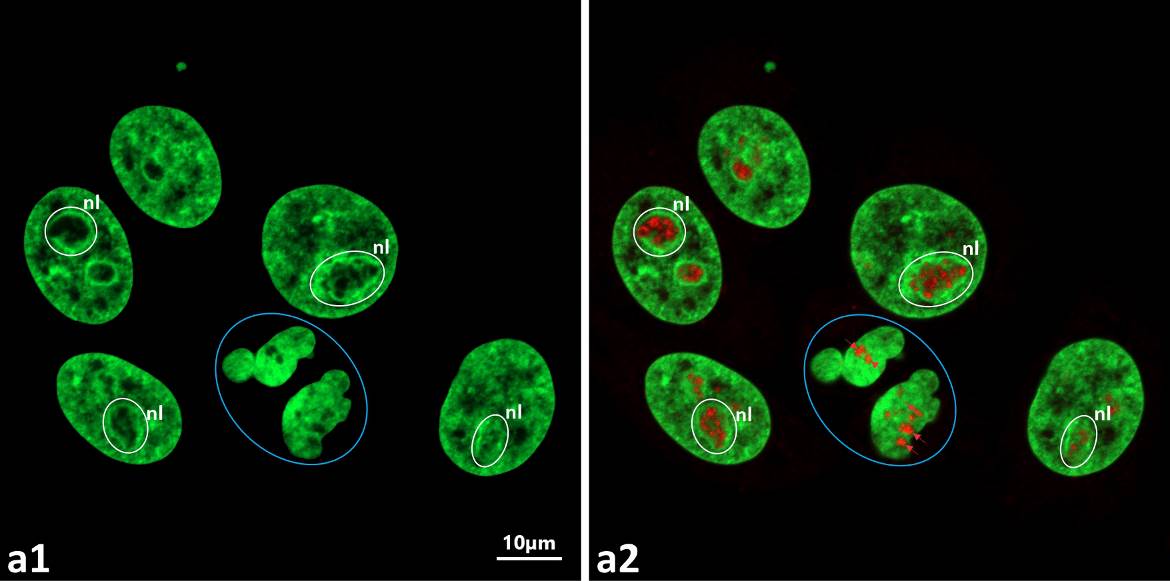

**S18 Figure.** 30 Gy post-γ-irradiation image acquisition during 24 hours. Anti-fibrillarin immunolabeling; zoomed image. Evidence that despite obvious predominance of mononuclear cells with smoothly outlined nuclei, initial signs of multinucleation/micronucleation (outlined by blue circles) also present. (a,1) GFP fluorescence only; nucleolar territory is outlined by white circles. Thickening of PCC and coarsening/clumping of ICC were obvious. (a2) Merging GFP fluorescence with anti-fibrillarin label (red) confirms transformation of typical cord-like organization in multinuclear cells into separate spherical fibrillarin-positive entities. In mononuclear cells anti-fibrillarin labeled retains cord-like organization. Abbreviations as on previous figures.

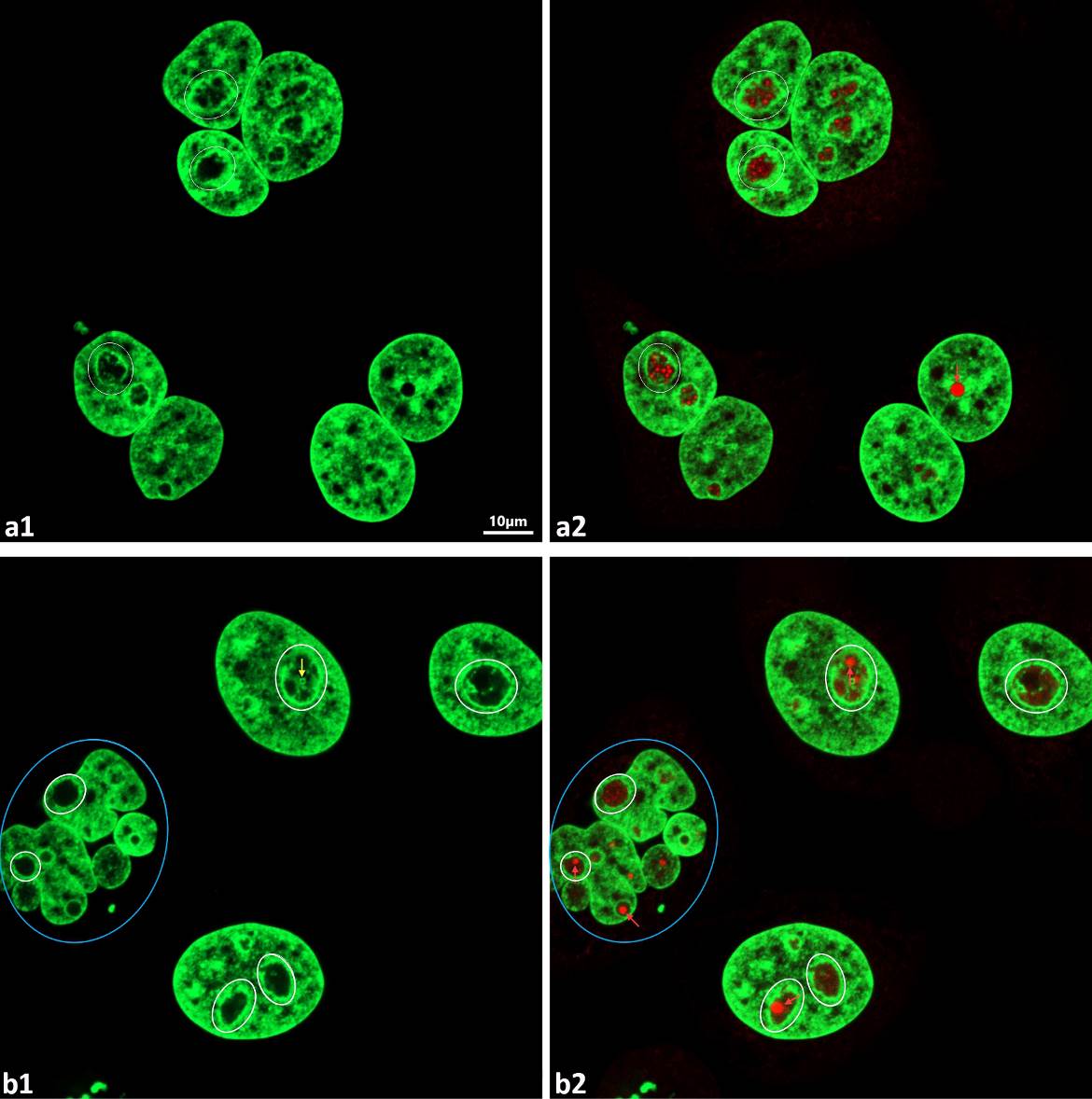

**S19 Figure.** 30 Gy post-γ-irradiation image acquisition during 48 hours. (a1-b2) Anti-UBTF immunolabeling. Nucleolar territory was outlined by white circles. Images reflecting formation of bi-nuclear, three-nucleolar (19, a1, a2) and post-MC multinuclear (19, b1, b2; outlined by blue circles) cells. (a1, b1) GFP fluorescence only; note thickening of PCC and coarsening/clumping of ICC. (a2, b2) Merged images prove presence of giant UBTF-positive structures (red, marked by red arrow) in mononuclear and bi-nuclear cells. In multi-nuclear cells, UBTF-positive spheres (marked by red arrow) look like they are dispersed among nuclear fragments. Abbreviations as on previous figures.

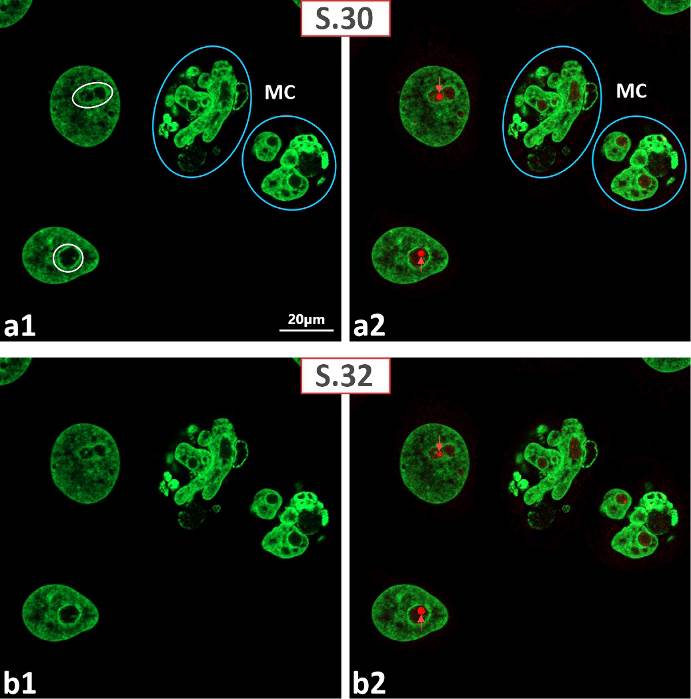

**S20 Figure.** 30 Gy post-γ-irradiation image acquisition during 48 hours. (a1-b2) Anti-UBTF immunolabeling witnessing the same changes as in S19 Fig., but was performed using serial sectioning; demonstration of non-successive sections extracted from a complete series (S.30-S.32). Nucleolar territory was outlined by white circles. Post-MC multinuclear cells (MC) were outlined by blue circles. (a1, b1) GFP fluorescence only. (a2, b2) By merging of GFP and anti-UBTF labeled images the presence of giant UBTF-positive structures (red, marked by red arrow) in mononuclear cells was obvious on all planes of series. In multinuclear cells UBTF-positive label looks like chaotically dispersed in nuclear fragments. Abbreviations as on previous figures.

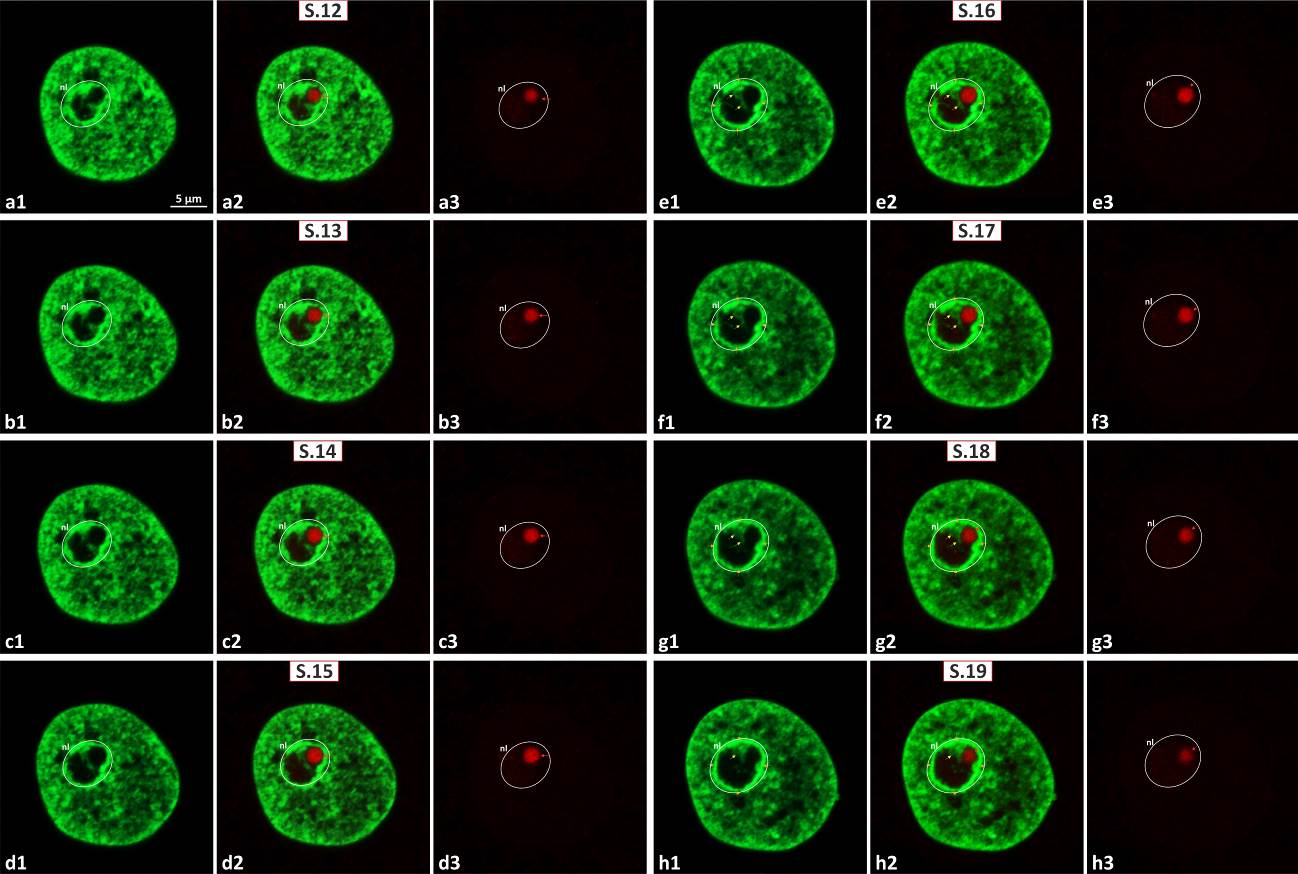

**S21 Figure.** 30 Gy post-γ-irradiation image acquisition during 48 hours. (a1-h3) Anti-UBTF immunolabeling; nucleolar territory was outlined by white circle. Extended demonstration of giant UBTF-positive spheres emerging as a result of γ-irradiation; the gallery of successive virtual serial sections (S.12-S.19) of nucleus showing giant intranucleolar UBTF-positive sphere (red, marked by red arrows) tightly anchored to thick PCC ring. Note also contact between ICC clump (marked by yellow arrows) and UBTF-positive sphere, visible through all series. (a1-h1) GFP label only; structural changes of NAC were clearly depicted. (a2-h2) GFP label merged with anti-UBTF immunolabeling images. Contact of UBTF-positive structure with PCC and ICC was obvious on all planes of series. (a3-h3) Anti-UBTF label only; indeed, serial sectioning of giant UBTF-positive structure (outlined by white circle) confirmed its spherical shape.

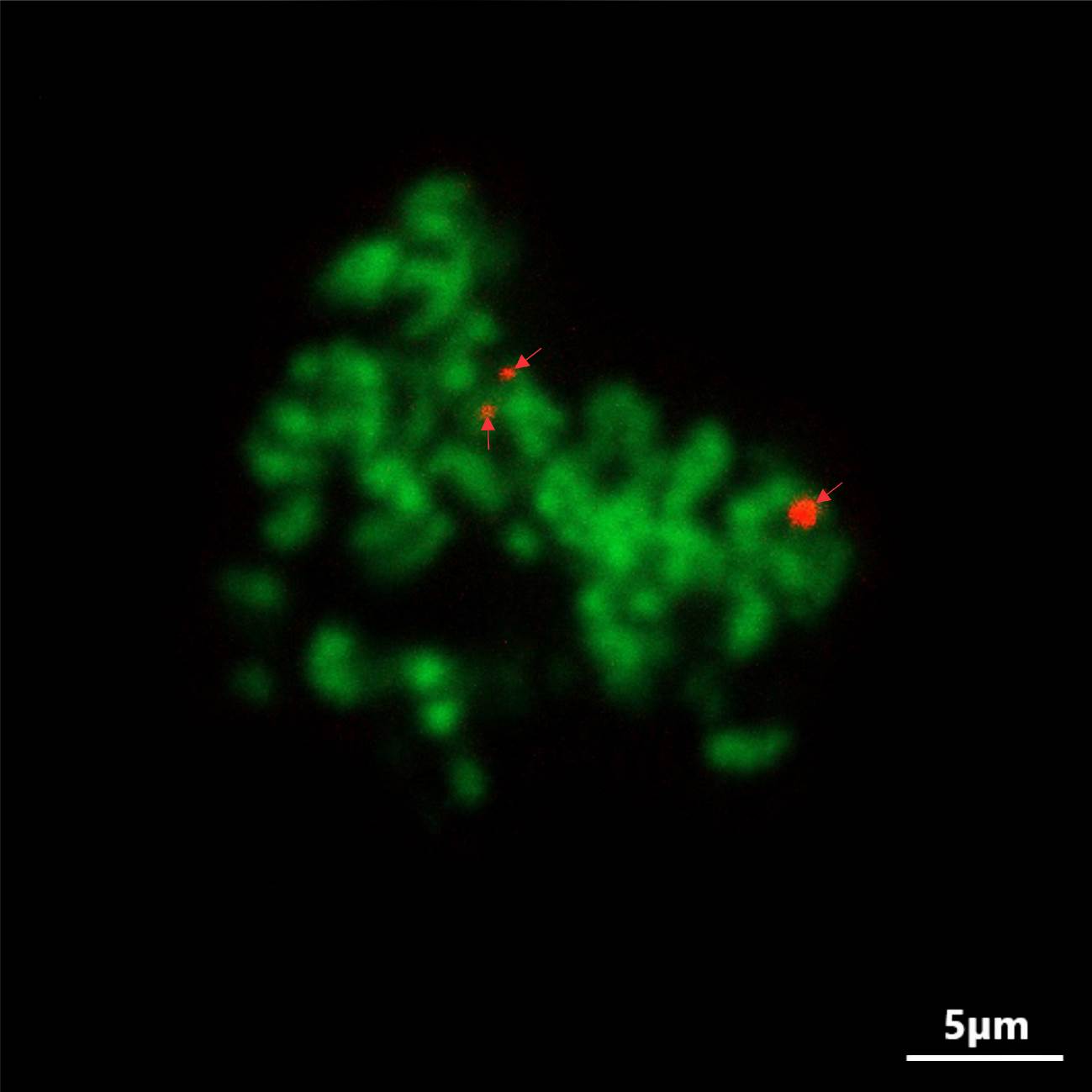

**S22 Figure.** 30 Gy post-γ-irradiation image acquisition during 48 hours. (a1-h3) Anti-UBTF immunolabeling. Mitotic cell revealed well developed UBTF-positive NORs (red, marked by red arrows) even after 48 hours of post-irradiation image acquisition.

**S23 Figure.** 30 Gy post-γ-irradiation image acquisition during 48 hours. (a1) Additional evidence that giant UBTF-positive spheres develop as a result of γ-irradiation. Nucleolar territory (outlined by white circle) contains giant UBTF-positive sphere (red, marked by red arrows). Its integration into NAC system was especially well pronounced. Note contact between ICC clump (marked by yellow arrow) and UBTF-positive sphere. Beside giant spheres, numerous tiny UBTF-positive structures were presented. (a2) 3D model (UCSF Chimera) showing same nucleus as on Fig. 23, a1, containing nucleoli with one giant and numerous tiny UBTF-positive structures. Adherence of the giant UBTF-positive sphere to the PCC shall was clearly seen. (a3) 3D model of extracted nucleolar territory to prove tight integration of UBTF-positive structures into NAC system. (b1, b2) Same as on previous figures. In contrast to previous UCSF Chimera reconstructions, 3D models presented on S23, a3, b2 Figures were generated using solid rendering mode for better demonstration how UBTF-positive structures immersed into NAC system. Abbreviations as on previous figures.

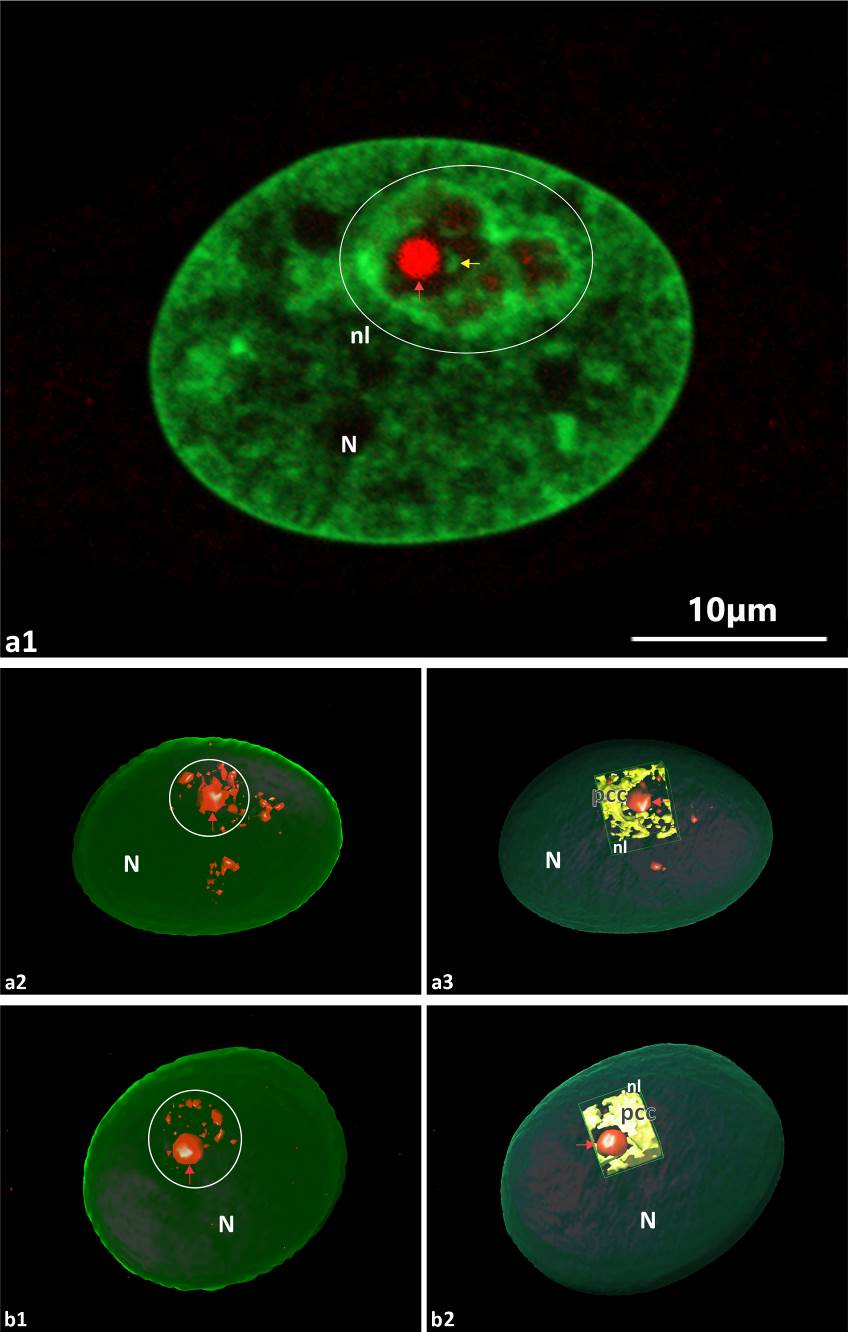

**S24 Figure.** 30 Gy post-γ-irradiation image acquisition during 48 hours. (a1-b2) Anti-fibrillarin immunolabeling (red); two non-successive serial sections (S.35, S.39) extracted from complete series. Evidence that even after 48 hours of image acquisition the majority of cells were mononuclear, while nuclear and nucleolar (outlined by white circles) microstructure resembling control. Note also presence of mitotic cells (outlined by blue circles). (a1, b1) GFP fluorescence only. (a2, b2) Merged GFP and anti-fibrillarin labels.

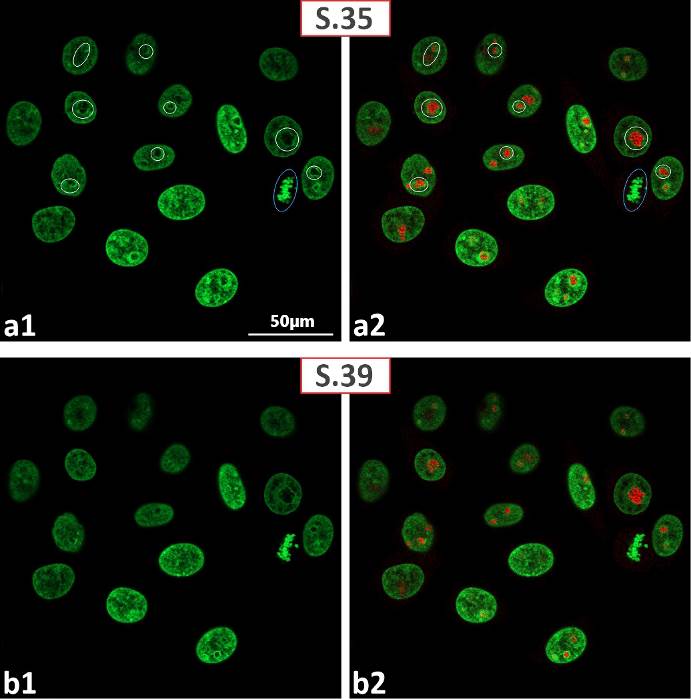

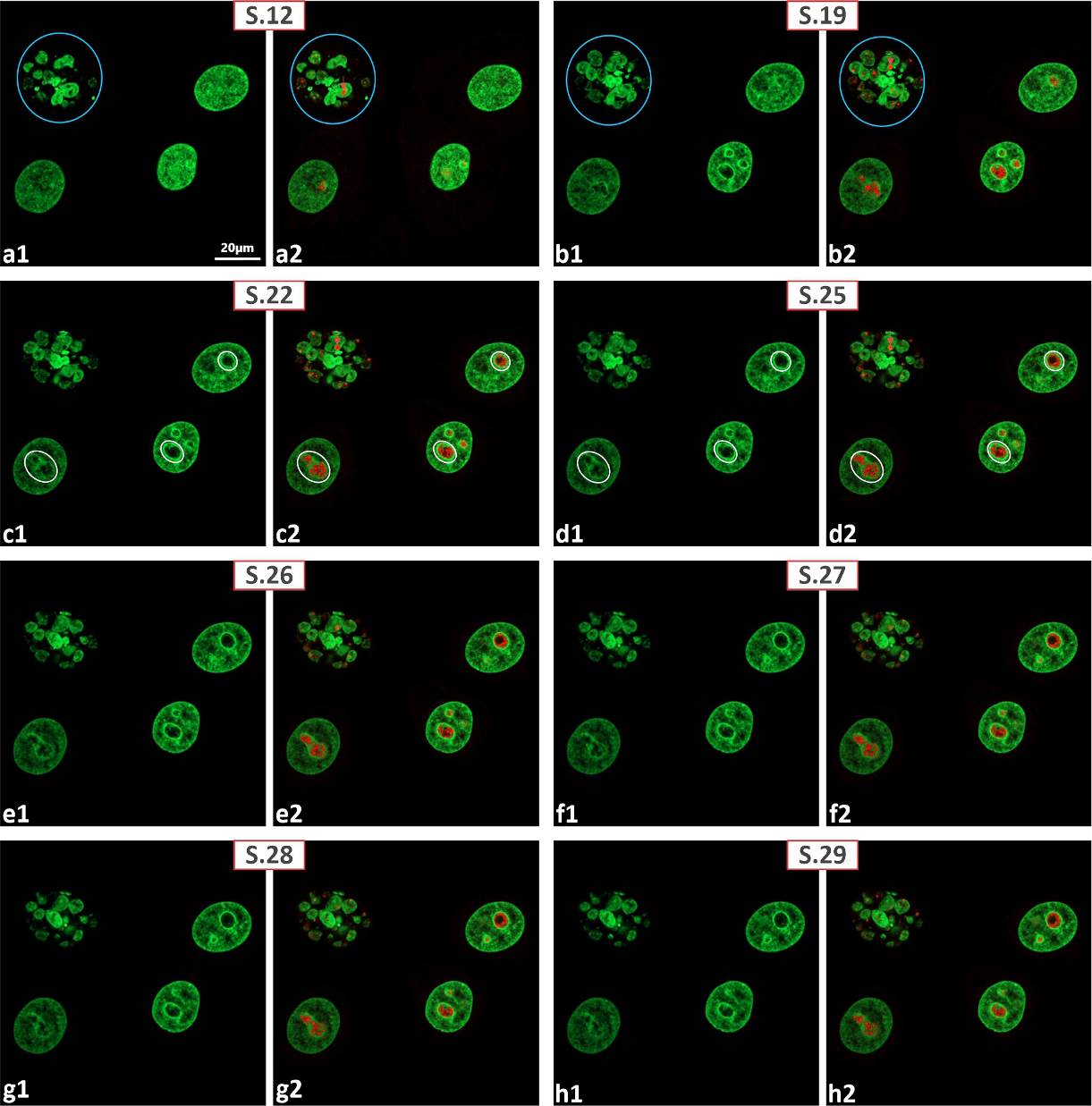

**S25 Figure.** 30 Gy post-γ-irradiation image acquisition during 48 hours. (a1-h2) Anti-fibrillarin immunolabeling; non-successive virtual serial sections (S.12, S.19, S.22, S.25, S.26-29) extracted from complete series. Evidence that despite obvious predominance of mononuclear cells with smoothly outlined nuclei, post MC multinuclear cells (outlined by blue circles) also present. (a1-h1) GFP fluorescence only. (a2-h2) By merging GFP fluorescence with anti-fibrillarin label (red) it became evident that in multinuclear cells cord-like structure transform into spherical fibrillarin-positive entities chaotically dispersed among nuclear fragments. Mononuclear cells maintained cord-like organization of anti-fibrillarin labeled structures.

**S26 Figure.** 30 Gy post-γ-irradiation image acquisition during 48 hours. (a1-h3) Anti-fibrllarin immunolabeling (red); non-successive virtual serial sections (S.12, S.14, S.16, S.18, S.20, S.22, S.26, S.30) extracted from complete series. (a1-h1) GFP fluorescence only. Evidence that even by sufficient compaction of ICC in nucleoli (outlined by white circles) in mononuclear cells, cord-like organization of fibrillarin-positive structures was remained. (a2-h2) Merged GFP fluorescence and anti-fibrillarin labels; obvious integration of fibrillarin-positive structures into NAC unit visible on all planes of series. (a3-h3) Anti-fibrillarin label only; nucleolar territory is outlined by white circles; clear evidence of maintained cord-like structure visible on all planes of series. Abbreviations as on previous figures.

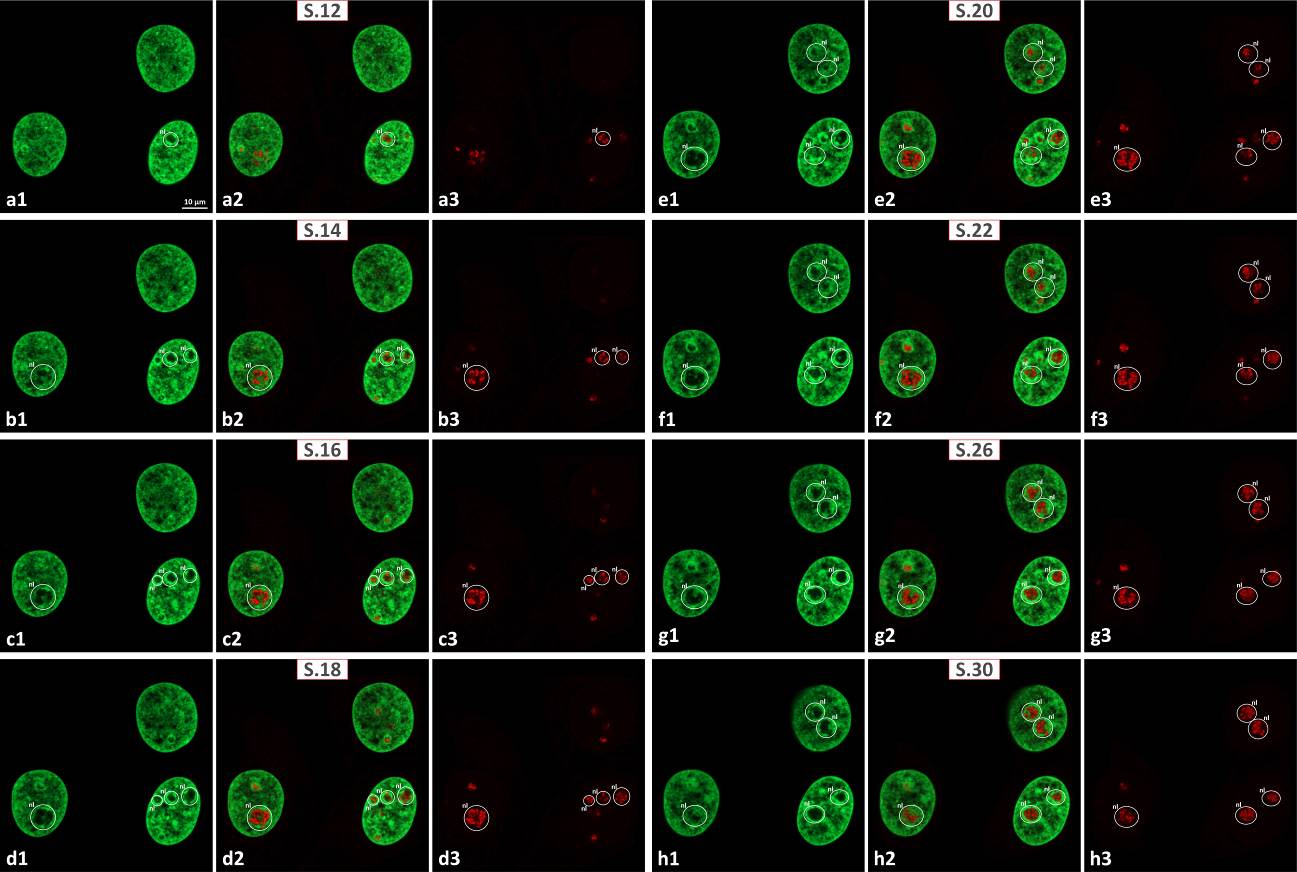

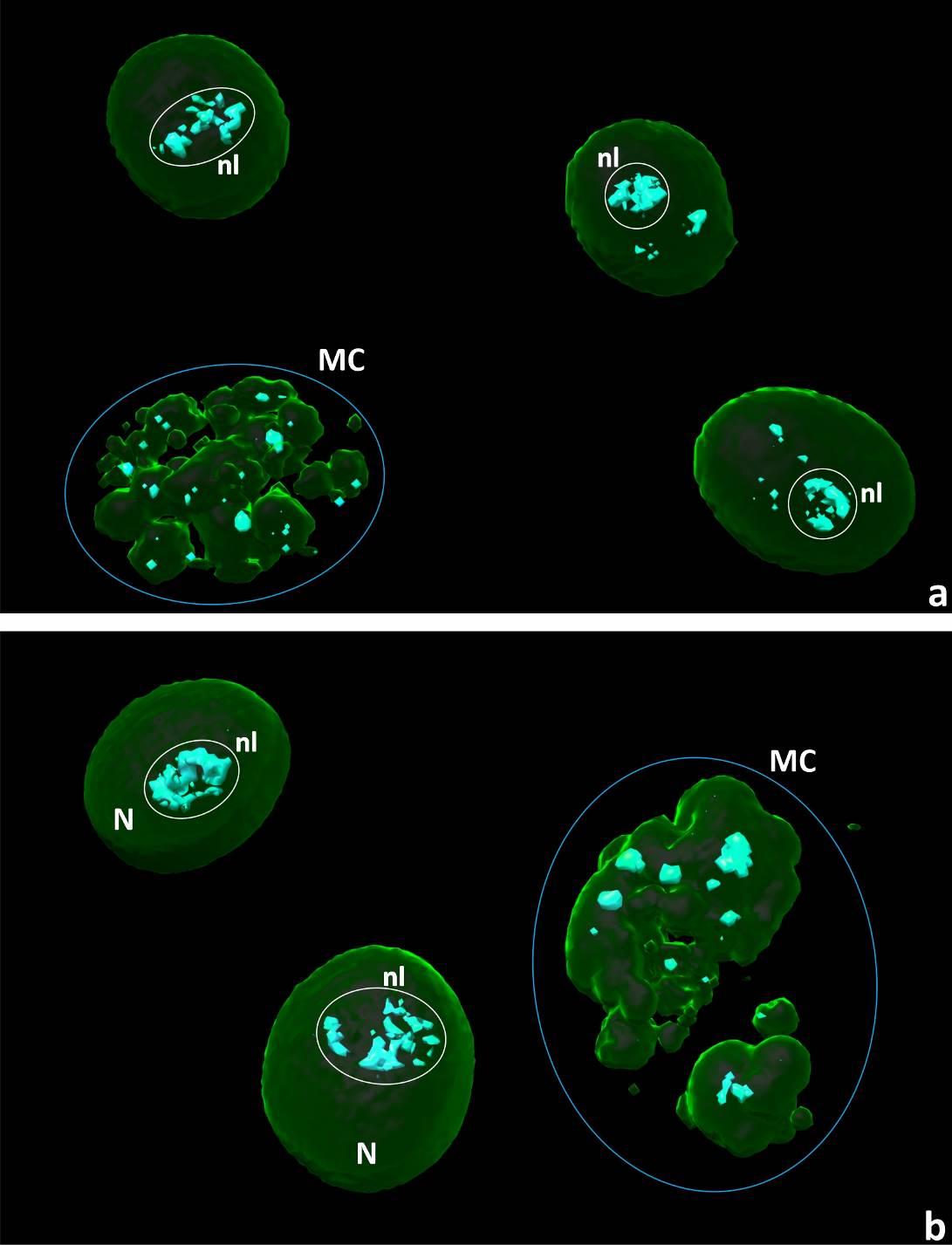

**S27 Figure.** 30 Gy post-γ-irradiation image acquisition during 48 hours. 3D models of anti-fibrillarin labeled (cyan) samples. Evidence of drastic differences between cord-like organization of fibrillarin-positive structures retained in mononuclear cells and those underwent disruption within nuclear fragments of multinuclear (27, a) cells and cells with lobbed nuclei (27, b). Abbreviations as on previous figures.

**S28 Figure.** 30 Gy post-γ-irradiation image acquisition during 72 hours. (a1-j3) Anti-UBTF immunolabeling (red); successive virtual serial sections (S.10-S.19) extracted from complete series. Nucleolar territory was outlined by white circle. Extended evidence that even 72 hours of post-irradiation image acquisition mononuclear cells (probably senescent cells) were presented in sufficient amount. Demonstration of giant UBTF-positive spheres emerged in nucleoli (outlined by white circles) of mononuclear cells as a consequence of γ-irradiation. (a1-j1) GFP label only; note compaction of NAC. (a2-j2) Merged GFP and anti-UBTF labels; integration of UBTF-positive spheres into NAC system. (a3-j3) Anti-UBTF label only; serial sections confirmed the spherical shape of UBTF-positive sphere (outlined by white circle) visible on all virtual planes. Abbreviations as on previous figures.

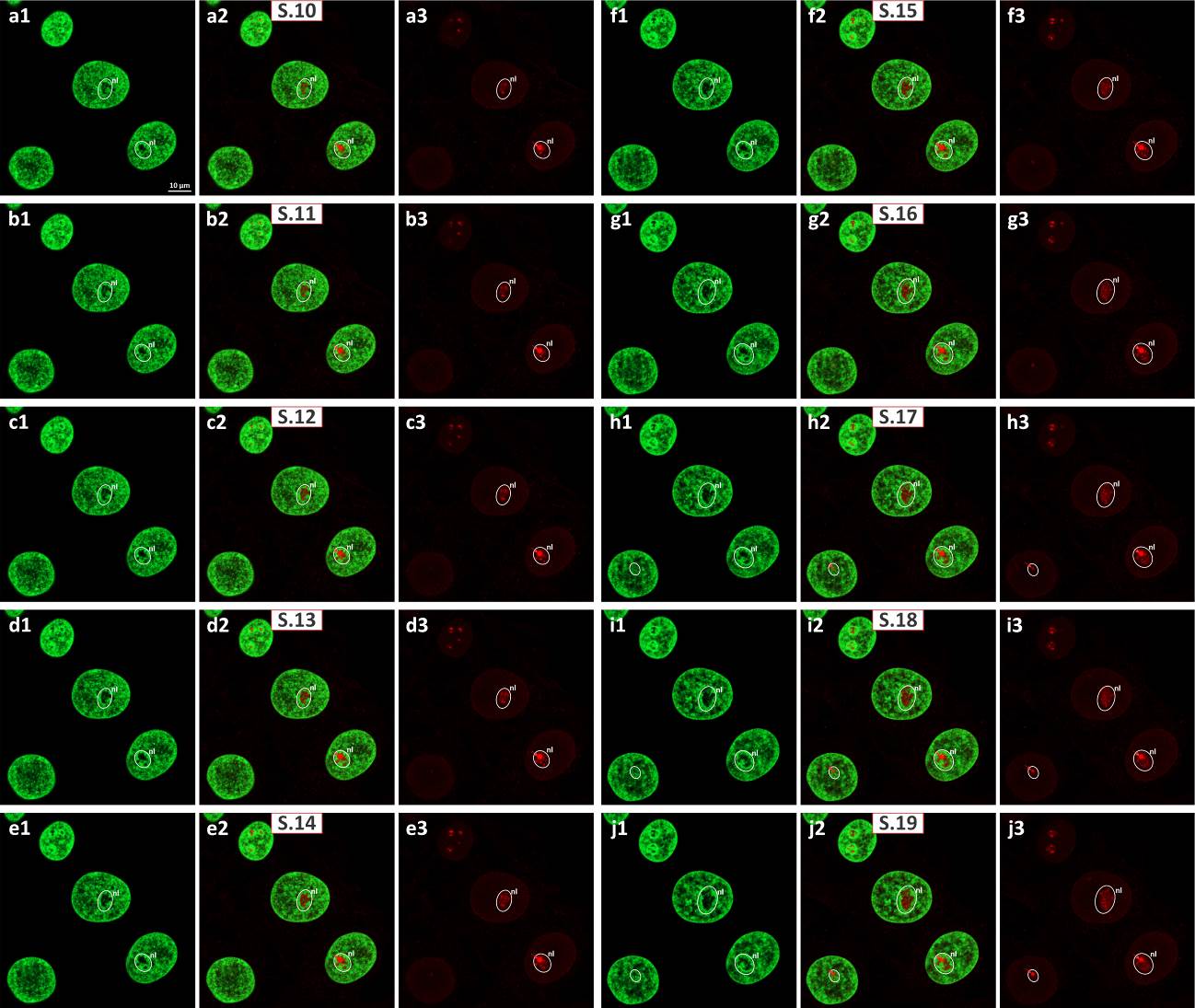

**S29 Figure.** 30 Gy post-γ-irradiation image acquisition during 72 hours. (a1-h3) Anti-UBTF immunolabeling (red); successive virtual serial sections (S.16-S.23) extracted from complete series. Same changes as on previous figure.

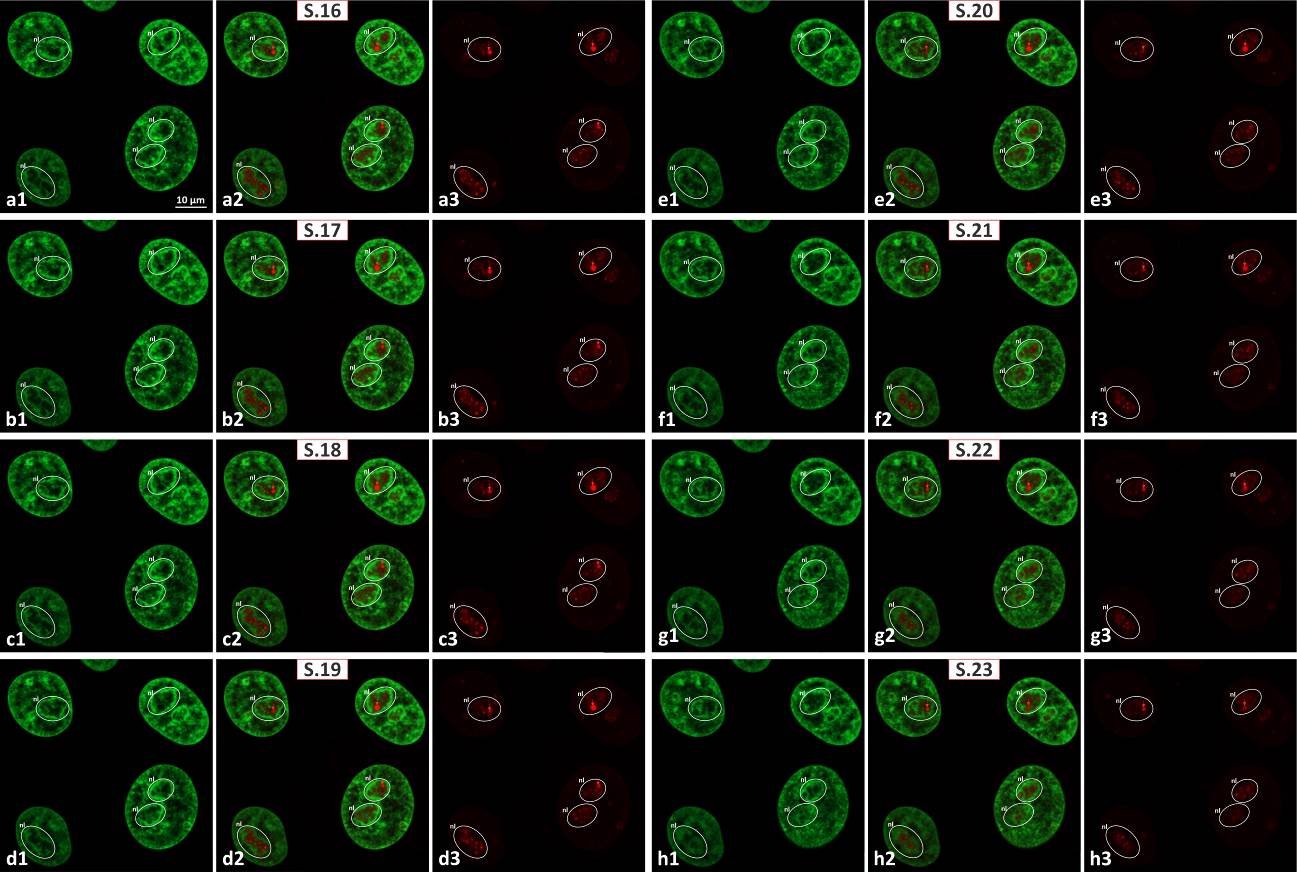

**S30 Figure.** 30 Gy post-γ-irradiation image acquisition during 72 hours. (a1-d2) Anti-UBTF immunolabeling (red). Non-successive virtual serial sections (S.16-S.18, S.26), performed to prove nuclear lobulation/micronucleation process (30, a1-d1, outlined by blue circles). Nucleolar territory was outlined by white circles. Note that three-nuclear cell (30, d1) in fact contained one, but intensively lobed nucleus (30, a1-c1). (a1-d1) GFP fluorescence only; note compaction of NAC system. (a2-d2) Merged GFP and anti-UBTF labels witnessing presence of UBTF-positive spheres inside nuclear lobes. Integration of UBTF-positive spheres into NAC unit was obvious.

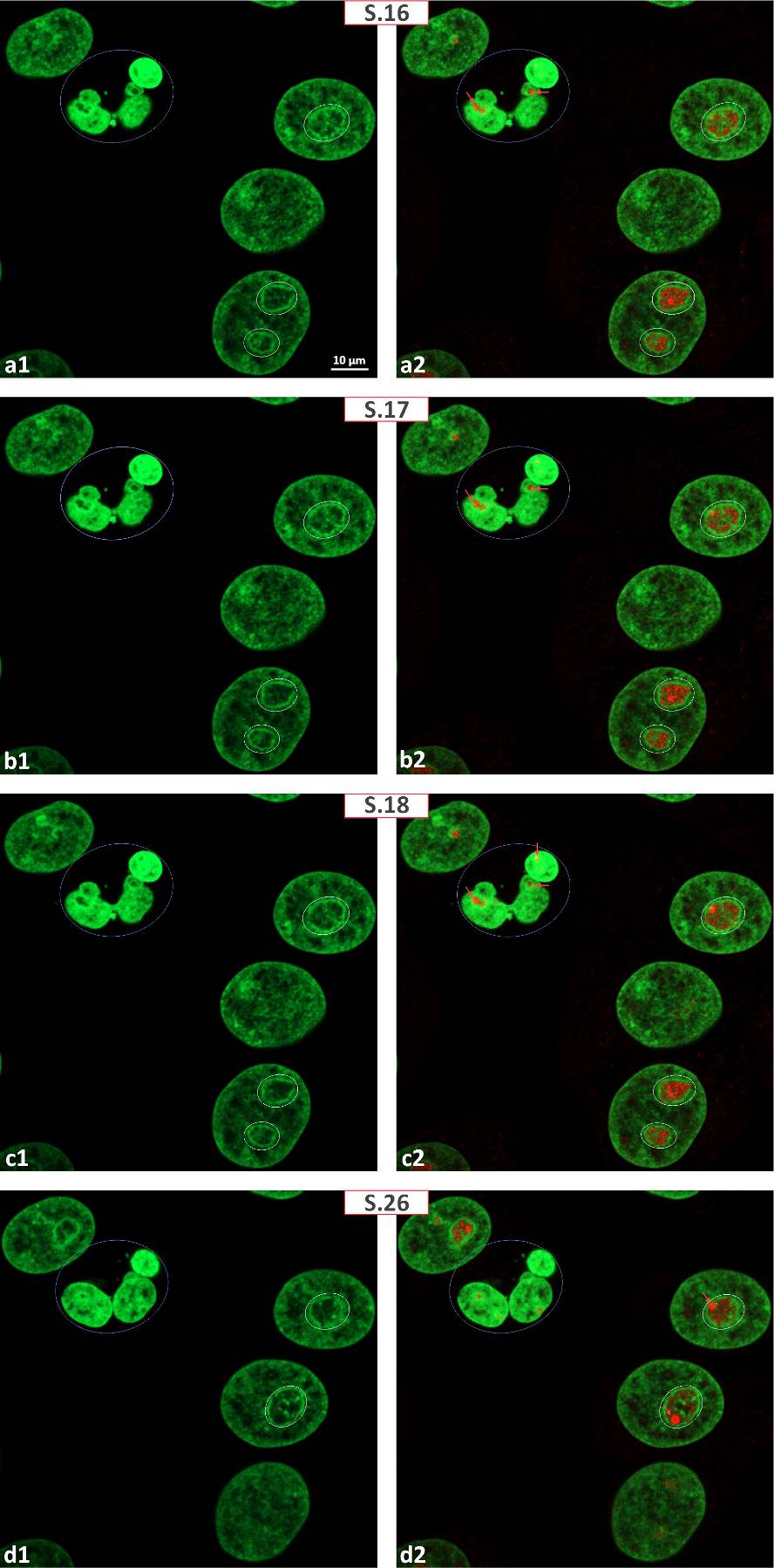

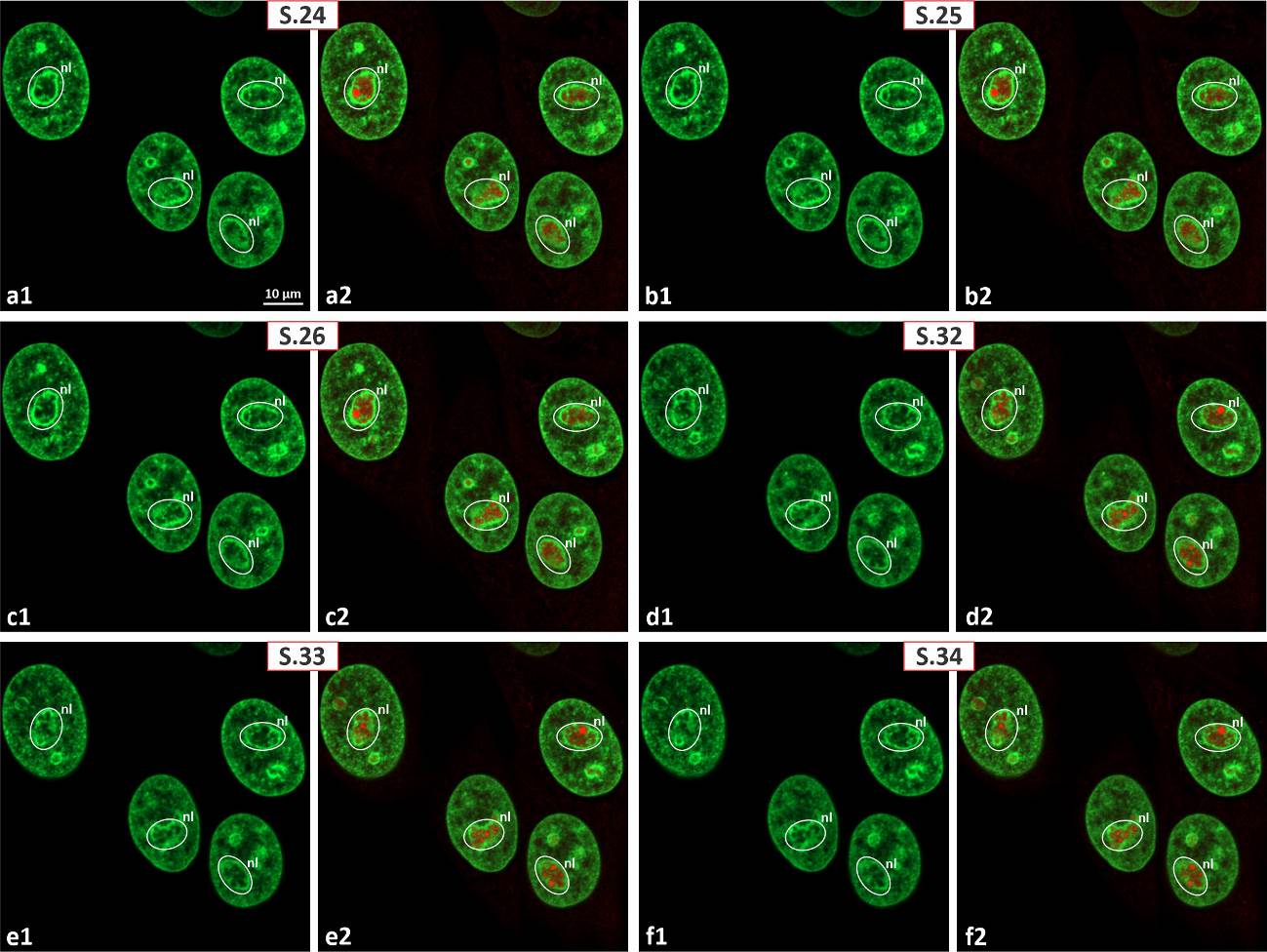

**S31 Figure.** 30 Gy post-γ-irradiation image acquisition during 72 hours. (a1-f2) Anti-UBTF immunolabeling (red). Non-successive virtual sections (S.24-S.34, S.32-S.34) extracted from complete series. Extended evidence of changes as on S28 and S29 Figures.

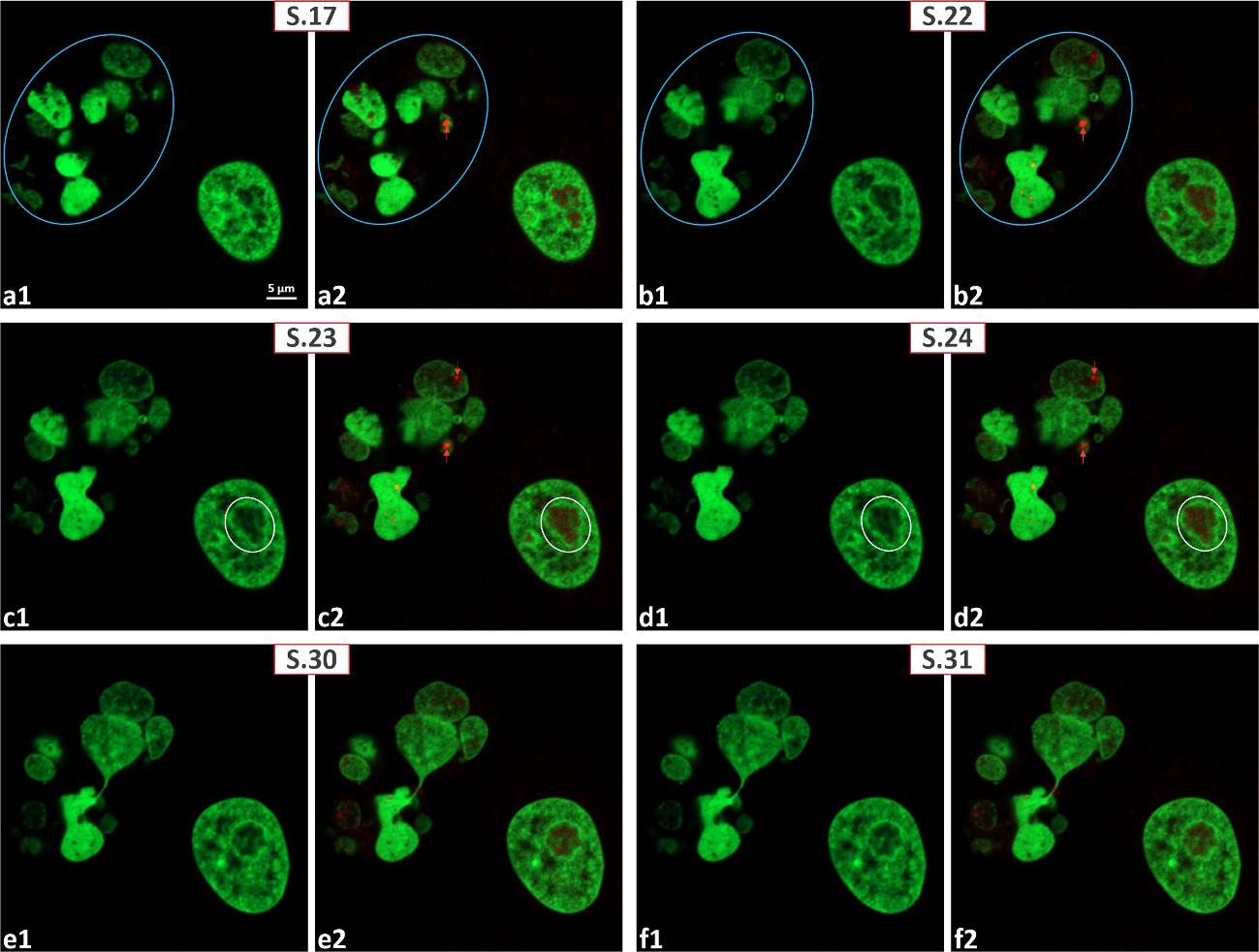

**S32 Figure.** 30 Gy post-γ-irradiation image acquisition during 72 hours. (a1-f2) Anti-UBTF immunolabeling (red). Non-successive virtual sections (S.17, S.22-S.24, S.30, S.31) extracted from complete series. Evidence of presence in nuclear fragments of multinuclear cells UBTF-positive spheres.

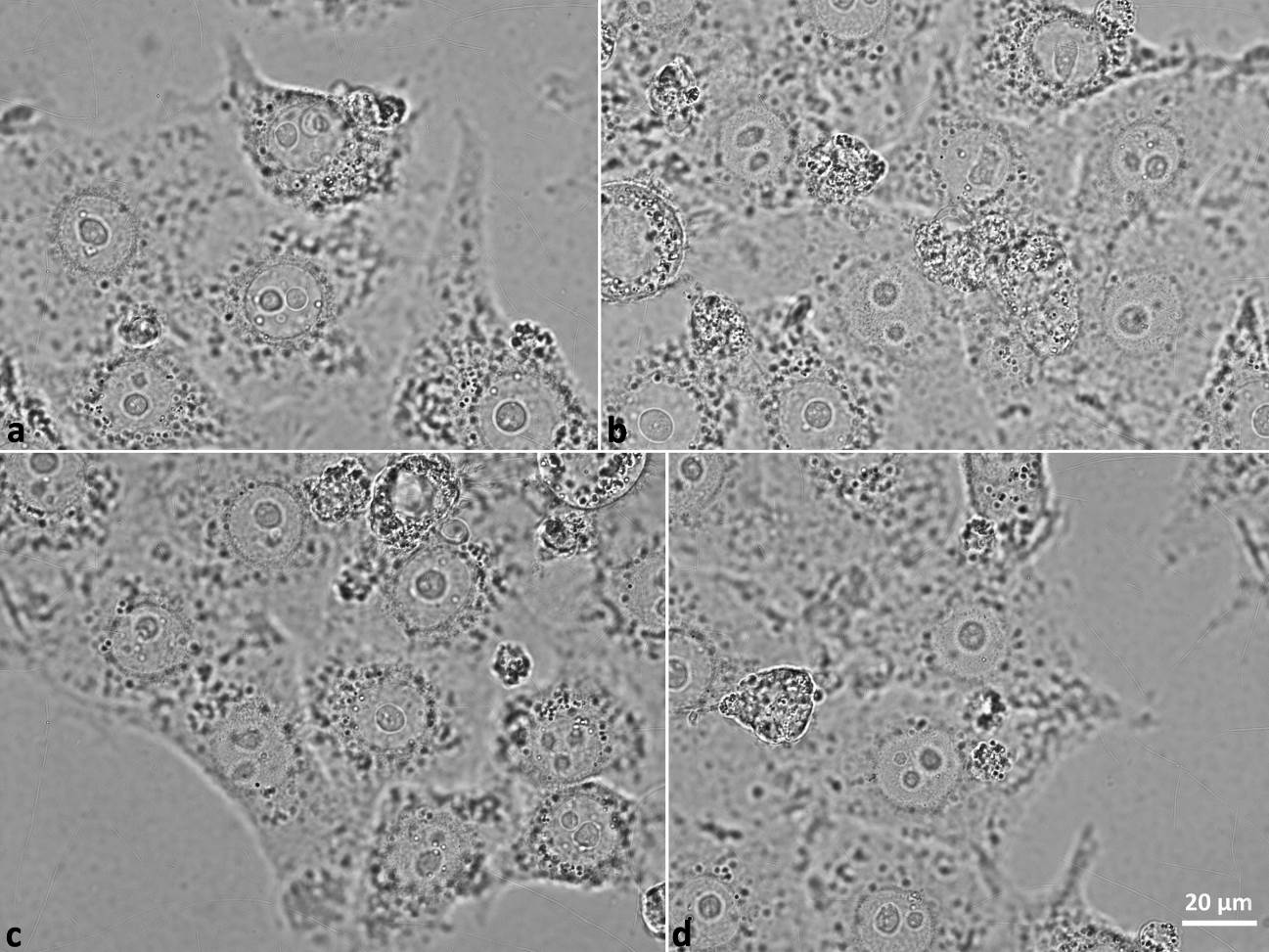

**S33 Figure.** 30 Gy post-γ-irradiation image acquisition during 72 hours. (a-d) PC images evidenced presence of large light zones in nucleoli of irradiated cells defined as nucleolinus [100, 101].

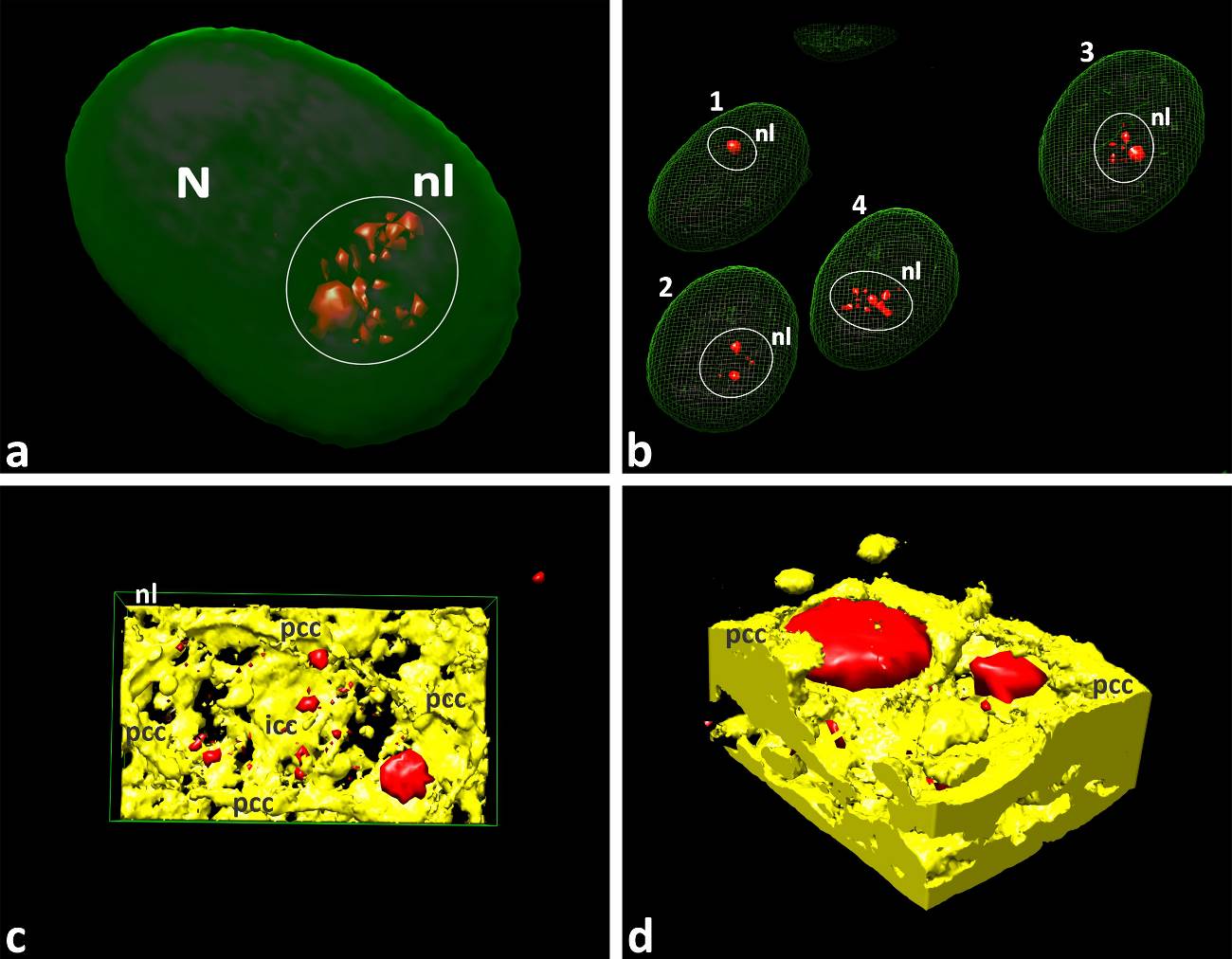

**S34 Figure.** 30 Gy post-γ-irradiation image acquisition during 72 hours. (a-d) 3D models of cells revealing giant UBTF-spheres as a consequence of 72 hours’ post-irradiation time-lapse imaging. (a) 3D model of the nucleus containing nucleolus with one giant sphere (red) and numerous tiny UBTF-positive structures. Nucleolar territory was outlined by white circle. (b) Demonstration of four nuclear 3D models revealing consequently only one (nucleus #1), two (nuclei #2, 3) and three (nucleus #4) giant UBTF-positive spheres. (c) 3D model of intranucleolar content extracted from nuclear volume showed on S34, a Figure. (d) 3D model of intranuclear content extracted from nucleus #3. Models showed on S34, c and S34, d Figures were constructed using solid rendering for better demonstration how UBTF-positive structures was immersed into NAC unit. Abbreviations as on previous figures.

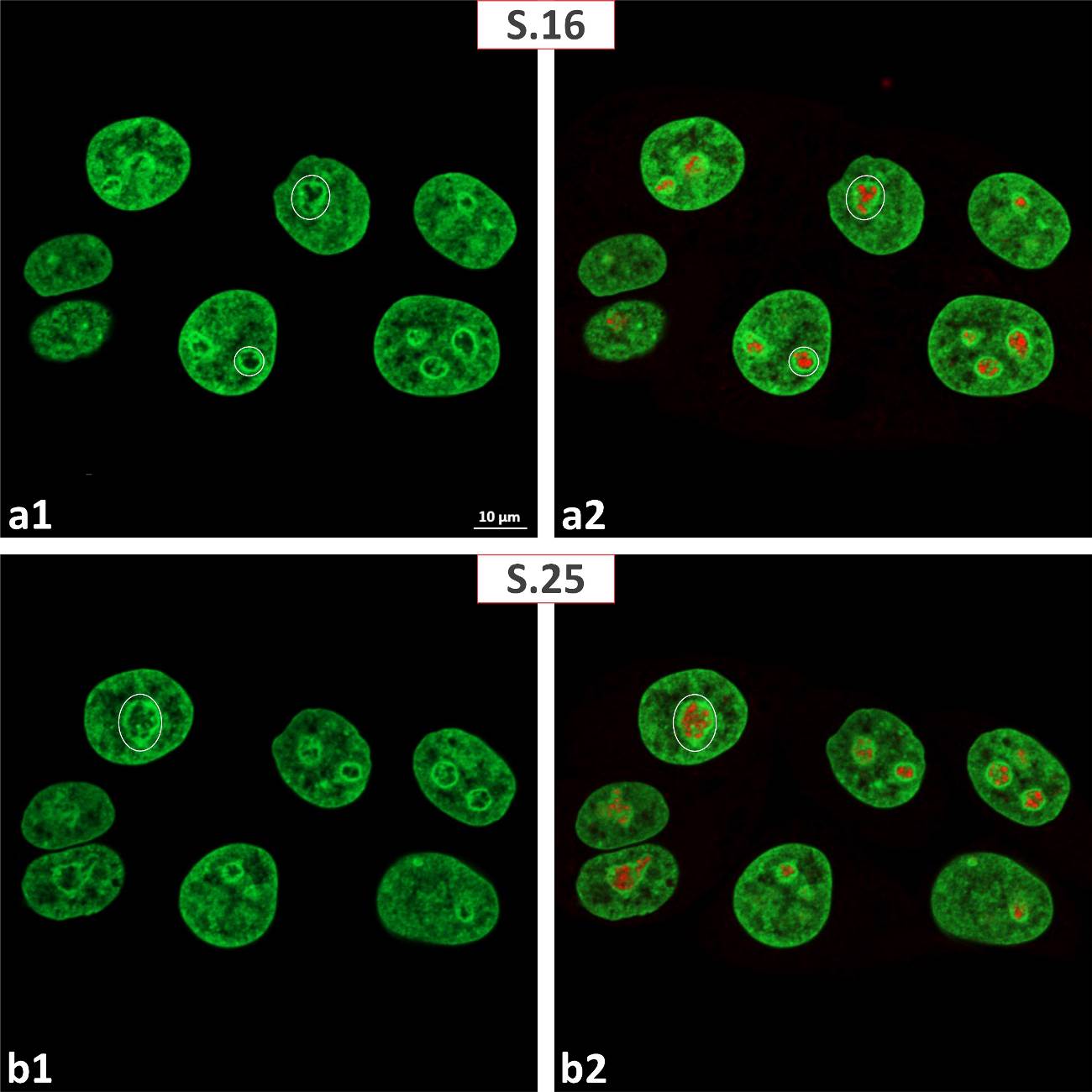

**S35 Figure.** 30 Gy post-γ-irradiation image acquisition during 72 hours. (a1-b2) Anti-fibrllarin immunolabeling (red); non-successive virtual serial sections (S.16 and S.25) extracted from complete series. Nucleolar territory was outlined by white circles. Extended evidence that even after 72 hours of post-irradiation image acquisition fibrillarin-positive structures in mononuclear cells retain cord-like appearance. (a1-b1) GFP fluorescence only. (a2-b2) Merged GFP fluorescence and anti-fibrillarin labels.

**S36 Figure.** 30 Gy post-γ-irradiation image acquisition during 72 hours. (a- d2) 3D models of anti-fibrillarin labeled (cyan) samples. Nucleolar territories were outlined by white circles. (a, b) Evidence that even after 72 hours of post-irradiation fibrillarin-positive structures retain cord-like organization. These nuclear models were constructed using transparent (36, a) and outlined (36, b) rendering. (c1) 3D model presenting position of nucleolus and NAC complex (taken in rectangle) inside nuclear volume. (c2) Zoomed 3D model showing integration of fibrillarin-positive cords into NAC system. (d1, d2) 3D models showing same structural interplay as on S36, c1, c2 Figures. Abbreviations as on previous figures.

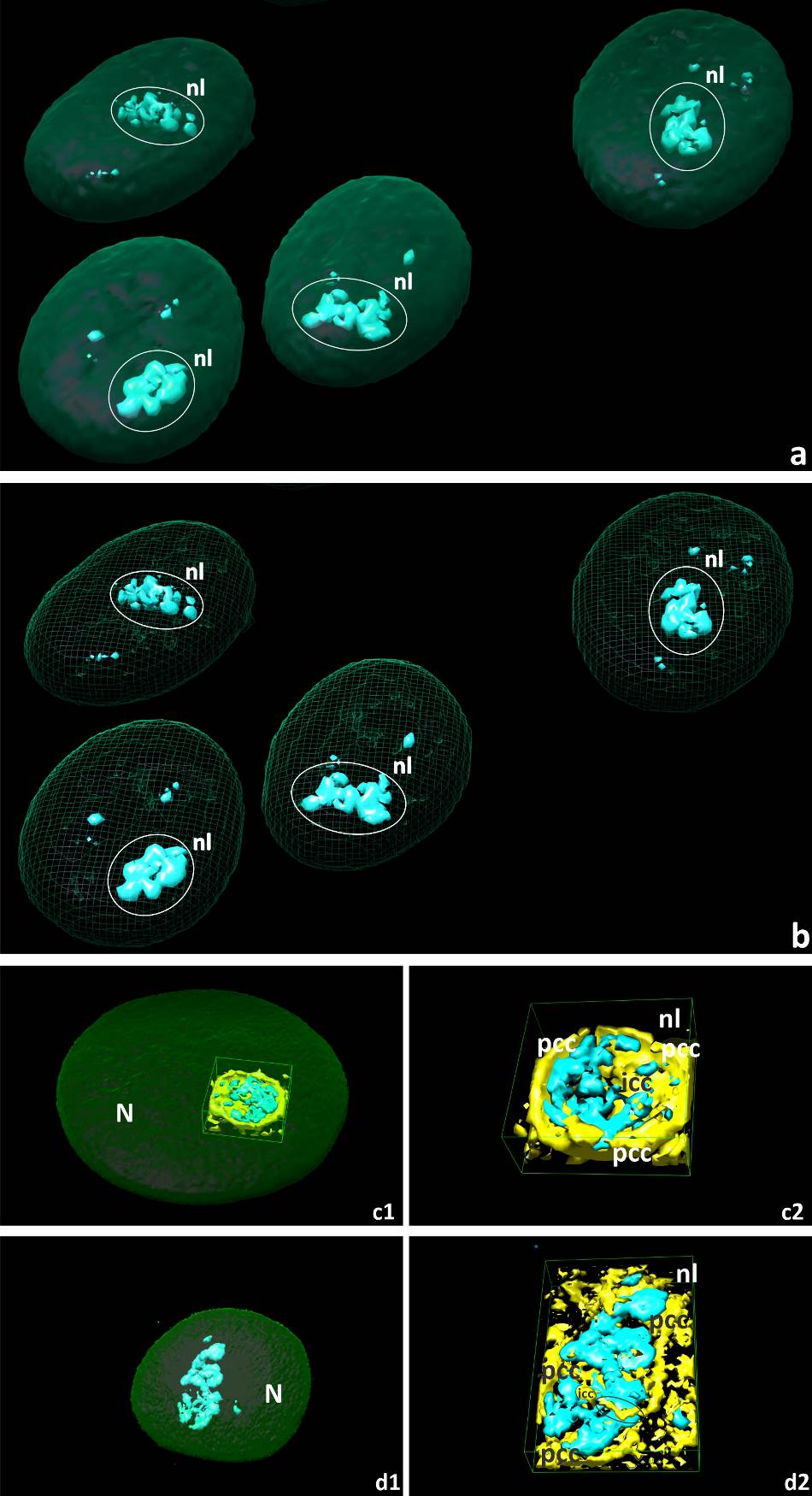
